# Development and utility of a PAK1-selective degrader

**DOI:** 10.1101/2022.05.12.491715

**Authors:** Hoi-Yee Chow, Sofiia Karchugina, Brian J. Groendyke, Sean Toenjes, John Hatcher, Katherine A. Donovan, Eric S. Fischer, Gleb Abalakov, Bulat Faezov, Roland Dunbrack, Nathanael S. Gray, Jonathan Chernoff

## Abstract

Amplification and/or overexpression of the *PAK1* gene is common in several malignancies, and inhibition of PAK1 by small molecules has been shown to impede the growth and survival of such cells. Potent inhibitors of PAK1 and its close relatives, PAK2, and PAK3, have been described, but clinical development has been hindered by recent findings that PAK2 function is required for normal cardiovascular function in adult mice. A unique allosteric PAK1-selective inhibitor, NVS-PAK1-1, provides a potential path forward, but has relatively modest potency in cells. Here, we report the development of BJG-05-039, a PAK1-seletive degrader consisting of the allosteric PAK1 inhibitor NVS-PAK1-1 conjugated to lenalidomide, a recruiter of the E3 ubiquitin ligase substrate adaptor Cereblon (CRBN). BJG-05-039 induced degradation of PAK1, but not PAK2, and displayed enhanced anti-proliferative effects relative to its parent compound in PAK1-dependent, but not PAK2-dependent, cell lines. Notably, BJG-05-039 promoted sustained PAK1 degradation and inhibition of downstream signaling effects at ten-fold lower dosage than NVS-PAK1-1. Our findings suggest that selective PAK1 degradation may confer more potent pharmacological effects compared with catalytic inhibition and highlight the potential advantages of PAK1-targeted degradation.

## INTRODUCTION

p21-activated kinases (PAKs) have been considered as potential drug targets in a variety of cancers ^1–4^. The PAK family is comprised of two groups: group A (PAK1, -2, and -3) and Group B (PAK4, -5-, and -6)^5^. The three members of the Group A PAKs are all closely related in sequence and structure, whereas the three Group B PAKs are distinct from the Group A proteins as well as more distantly related to one another. In addition to their structural differences, the six isoforms have distinct though sometimes overlapping expression patterns. For example, PAK1 is primarily expressed in brain, muscle, and blood cells, PAK2 is ubiquitous, and PAK3 is primarily expressed in neuronal cells. Of the Group B PAKs, PAK4 is ubiquitous, PAK5 is expressed mainly in neuronal cells, and PAK6 is expressed in neuronal cells, skin, prostate and testes ^4, 6^. Genetic loss-of-function analyses in animal models has shown a variety of different phenotypes, ranging from embryonic lethality (PAK2, PAK4), to cognitive dysfunction (PAK3), to minimal effects (PAK1, PAK5, PAK6) ^7, 8^.

As effectors of the small GTPase RAC, the PAK enzymes regulate several key proliferative and survival pathways including the RAF-MEK-ERK, the PI3K-AKT-mTORC, and the β-catenin pathways ^2^. While rarely subject to mutational activation, certain PAK isoforms, in particular PAK1 and PAK4, are frequently expressed at high levels in various tumor types due to chromosomal amplifications of their corresponding genes at chromosome 11q13 and 19q13, respectively ^2, 3^, or are activated by mutations in RAC1 ^9^. Reducing PAK activity, via gene knockouts, RNA interference, or small molecule inhibitors, has been shown to be of benefit in many cell-based and *in vivo* cancer models ^2^. These factors led several pharmaceutical companies to develop various PAK inhibitors for cancer therapy ^10–14^. One of these, Pfizer’s pan-PAK inhibitor PF3758309, was evaluated in a human Phase 1 clinical trial, but was withdrawn due to a combination of poor pharmaceutical properties and toxicity ^2^. Afraxis and Genentech described a series of increasingly specific Group A PAK inhibitors, which were found to be effective in preclinical models of NF2, KRAS-driven squamous cell carcinoma, and HER2- driven breast cancer ^11, 15–17^. However, development of this series of compounds was halted in 2016 due to evidence of on-target toxicity related to cardiovascular events ^14^. Similar findings were described in *Pak2* knockout mice, using a tamoxifen regulated CAGG-Cre-ERT gene to delete the floxed *Pak2* gene in adult mice ^18^. In contrast, deletion of the closely related genes for *Pak1* and *Pak3* was not found to be required for viability, development, longevity, or fertility ^7, 19^. These combined findings suggested that PAK2 function is required in adult mice and that small molecules that inhibit PAK2 might be toxic or even lethal in humans, muting enthusiasm for further development of Group A PAK inhibitors.

In 2015, Novartis described a dibenzodiazepine-based small molecule inhibitor, NVS-PAK1-1, that showed excellent specificity for PAKs over other kinases and a ∼50x selectivity for PAK1 over PAK2 ^20^. Given the extensive similarity between the catalytic domains of PAK1 and PAK2 (93% identical), this selectivity was unexpected, as was the co-crystal structure which revealed that the molecule occupied a space in the catalytic cleft underneath the αC-helix rather than in the hinge region of the ATP binding pocket. Selective blockade of PAK1 might be clinically useful in an animal model of NF2, as we recently reported that deletion of the *Pak1*, but not the *Pak2* gene, was effective in slowing hearing loss and schwannoma growth in these mice, and that treatment with NVS-PAK1-1 showed a similar trend without obvious systemic toxicity ^21^. As several cancer cells are PAK1-dependent, these findings suggested a path forward for PAK1- selective inhibitors. However, NVS-PAK1-1 has a short half-life in rat liver microsomes and, *in vivo*, is metabolized by the cytochrome P450 system ^20, 21^. To our knowledge, only one other class of small molecule inhibitors has displayed a similar selectivity for PAK1 over PAK2, which is achieved by binding to the less conserved p21-binding domain at the N-terminus of PAK1, as opposed to the highly conserved kinase domain ^22^. However, these compounds are only effective at micromolar doses.

Given that the signaling activity of PAK1 is mediated both by enzymatic and scaffolding functions ^6, 23^, we considered that a degrader derived from NVS-PAK1-1 might prove more potent than the parental molecule, while simultaneously retaining its selectivity for PAK1 over PAK2. We therefore synthesized and characterized a small collection of bivalent degraders (PROTACs) derived from conjugating NVS-PAK1-1 to lenalidomide which facilitate recruitment of the CRL4^CRBN^ ubiquitin ligase for substrate ubiquitination and eventual proteosome-mediated degradation. We anticipated that this bivalent degrader would preserve the unique isoform-specific PAK1 inhibitor activity while simultaneously being capable of inducing PAK1 protein degradation. In this report, we describe the properties of an optimized PAK1- specific degrader, as compared to the parental inhibitor, a negative degrader (analog disabled for binding to CRBN), and to shRNA-mediated gene knockdown. We found that the degrader form of NVS-PAK1-1 could be used at ten-fold lower doses to reduce growth signaling and proliferation of PAK1-dependent, but not PAK2-dependent cell lines. These effects could be enhanced by co-treatment with a drug efflux inhibitor. These results represent the first degrader form of a PAK inhibitor, and could form the basis for future development of PAK1-specific inhibitors.

## RESULTS AND DISCUSSION

### Design of a Selective PAK1 Degrader

To develop a PAK1-selective heterobifunctional degrader, we designed compounds based on NVS-PAK1-1, a unique allosteric inhibitor which displays marked selectivity for PAK1 over PAK2 (Figure 1A) ^20^. This compound is based on a dibenzodiazepine scaffold, which is uncommon for kinase inhibitors. Co-crystals of close relatives of NVS-PAK1-1 show that these molecules bind beneath the αC helix in PAK1 in a pocket formed in the DFG-out conformation of PAK1 analogous to the well-characterized allosteric inhibitors of MEK1/2 ^24^ and EGFR ^25^. It has been speculated that they derive specificity for PAK1 over PAK2 due to a steric clash between the molecule with Leu301 in PAK2 *vs.* the equivalent Asn322 in PAK1 ^20^ (Figure 1B). Such specificity is unusual and has rarely been described for other inhibitors of Group A PAK. For example, we tested NVS-PAK1- 1 and over 2000 hinge-region kinase inhibitors against PAK1 and PAK2 using an in vitro kinase assay. With the exception of NVS-PAK1-1, none of the other compounds displayed differential sensitivities between these two kinases (Figure 1C).

**Figure 1.**
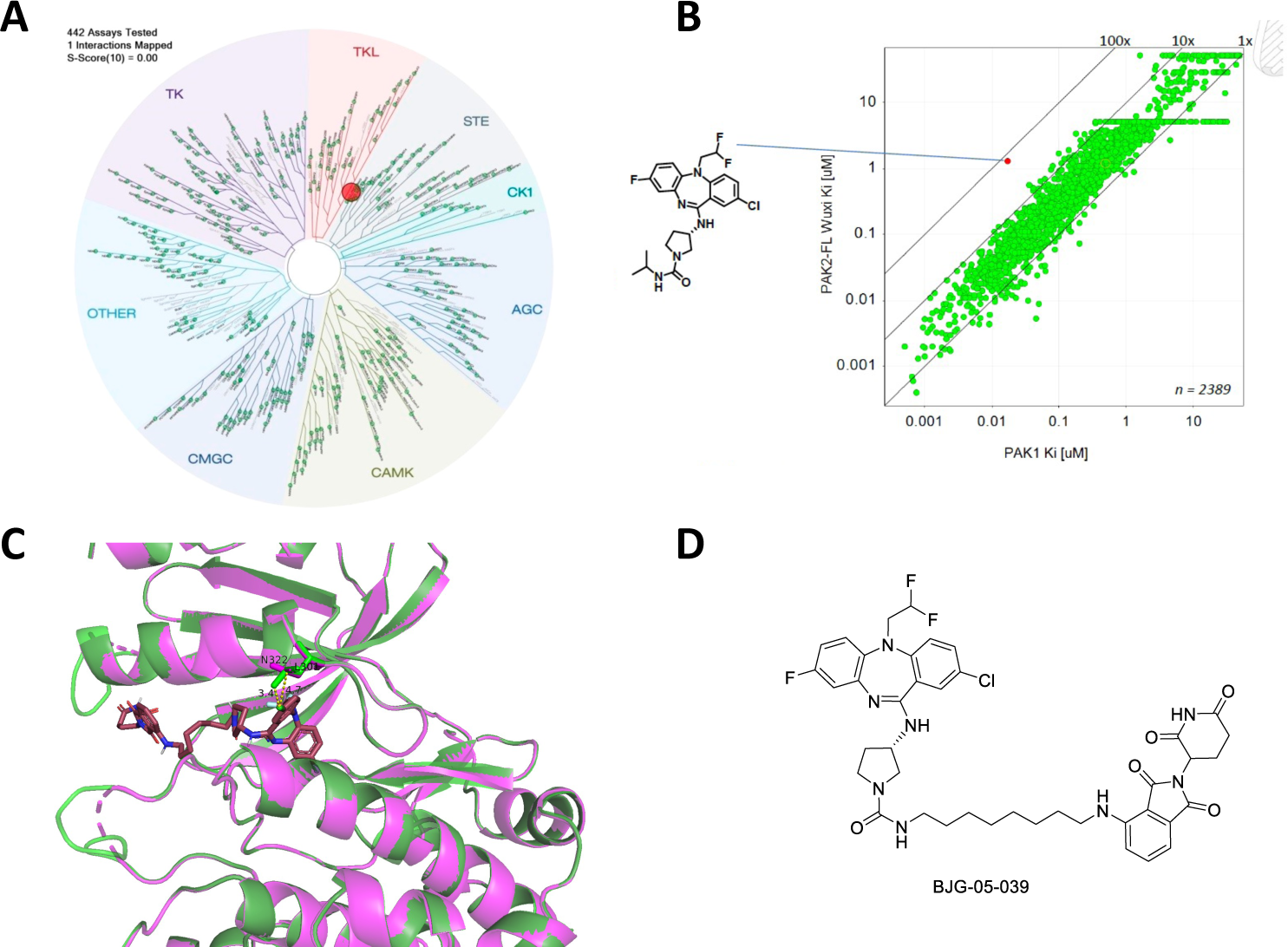
Structure and Selectivity of PAK1 Degrader. **A)** TREEspot visualization of the biochemical kinome selectivity of 1 μM NVS-PAK1-1. **B)** Proposed structural basis for isoform specificity. The crystal structure of PAK1 is depicted in magenta (PDB entry, 4ZJI, chain A). PAK2 in green was modeled using Swiss-Model using 4ZJI chain A as a template. BJG-05- 039 was modeled using Open Babel from its SMILES string and docked with AutoDock Vina. Amino acids that contact the ligand and differ between PAK1 and PAK2 are shown in sticks. N322 in PAK1 and L301 in PAK2 are labeled; the side chain of L301 forms a very close contact with the ligand (3.4 Å) compared to N322 in PAK1 (4.7 Å). **C)** Specificity of NVS-PAK1-1 compared to other Group A PAK inhibitors. *In vitro* kinase assays were performed using recombinant PAK1 and PAK2, using a library of 2389 hinge-binding small molecule protein kinase inhibitor compounds from Genentech’s PAK1 inhibitor program, plus NVS-PAK1-1. **D)** Chemical structure of BJG-05-039.

As NVS-PAK1-1 has a short *in vivo* half-life and has shown marginal effects in cancer cell lines ^20^, we attempted to create degrader forms of this compound. Such compounds might have the added advantage over conventional PAK inhibitors because, in addition to blocking kinase enzymatic activity, they also have the potential to reduce signaling effects that emanate from the scaffold functions of PAK1. A co-crystal structure of a close analogue of NVS-PAK1-1 bound to PAK1 (PDB: 4ZJJ) revealed that the isopropyl urea is solvent exposed, suggesting that the carbonyl (either as a urea or an amide) could serve as a suitable attachment site for linkers without adversely affecting affinity to PAK1 (Figure 1E).

Hydrocarbon and polyethylene glycol (PEG) linkers of varying lengths were used to conjugate NVS-PAK1-1 with either a CRBN ligand (lenalidomide) or a VHL ligand, respectively. To verify that conjugation of the linker and lenalidomide or VHL ligand did not affect the ability of NVS-PAK1-1 to bind to PAK1, ten different NVS-PAK1-1 conjugates were tested in a commercially available fluorescence resonance energy transfer-based assay (Invitrogen, Z′-Lyte) for PAK1, PAK2, and PAK4 inhibition (Table S1). Three of these – BJG- 05-014, BJG-05-027, and BJG-05-039 – retained PAK1 inhibitory activity and selectivity and were therefore selected for more detailed studies. While BJG-05-039 had lower inhibitory activity against PAK1 (half maximal inhibitory concentration (IC50) = 233 nM) compared to NVS-PAK1-1 (IC50 = 29.6 nM), it displayed greater specificity (PAK2 IC50 > 10,000 nM for BJG-05-039 *vs.* 824 nM for NVS-PAK1-1), demonstrating that BJG-05-039 had improved selectivity for PAK1 over PAK2 compared to the parent molecule. Molecular modeling suggests that the addition of the linker and degrader moieties would not impair selective binding of the inhibitory ligand in the ATP cleft of PAK1 (Figure S1).

BJG-05-014, BJG-05-027, and BJG-05-03 were tested in Panc1 cells for their ability to degrade PAK1. Of these three molecules, BJG-05-039, which uses an 8-carbon linker to conjugate NVS-PAK1-1 with lenalidomide, had the most promising profile (Figure S2).

We evaluated the biochemical selectivity of BJG-05-039 against a panel of 468 kinases at 10 μM (KINOMEscan) and observed that BJG-05-039 had a similar selectivity profile as 10 μM NVS-PAK1-1 (Figure S3 and Table S2).

### BJG-05-039 is a Highly Selective PAK1 Degrader

After verifying that BJG-05-039 retained specificity for PAK1, we sought to characterize its degradation activity in cells. We first chose to evaluate the PAK1 degrader in breast cancer cell line MCF7 and the ovarian cancer cell line OVCAR3 due to their high expression of both PAK1 and PAK2, and their known dependence on PAK1 ^26, 27^. We found that BJG-05-039 induced degradation of PAK1 but not PAK2 in a dose-dependent manner after a 12 h treatment, with maximal degradation observed at 10 nM (Figure 2A). At concentrations of 1 μM and greater, we observed diminished PAK1 degradation, consistent with the hook effect, in which independent engagement of PAK1 and CRBN by BJG-05-039 prevents formation of a productive ternary complex ^28^. PAK1 degradation was not observed in cells treated with the parent molecule, NVS-PAK1-1, or with BJG-05-098, a negative control compound with an N-methylated glutarimide that weakens CRBN binding (Figure 2A), demonstrating that BJG-05-039-induced PAK1 degradation was CRBN dependent^29^. Time course treatment of MCF7 cells with 250 nM BJG-05-039 revealed partial degradation of PAK1 within 4 h and progressive loss out to 24 h (Figure 2B). Co-treatment of cells with BJG-05-039 and bortezomib, a proteasome inhibitor, or MLN4924, an NAE1 (NEDD8- activating enzyme 1) inhibitor that prevents the neddylation required for the activation of cullin RING ligases, such as CRL4^CRBN^ ^30^, prevented PAK1 destabilization, indicating that degradation was dependent on the ubiquitin-proteasome system (Figure 2C).

**Figure 2.**
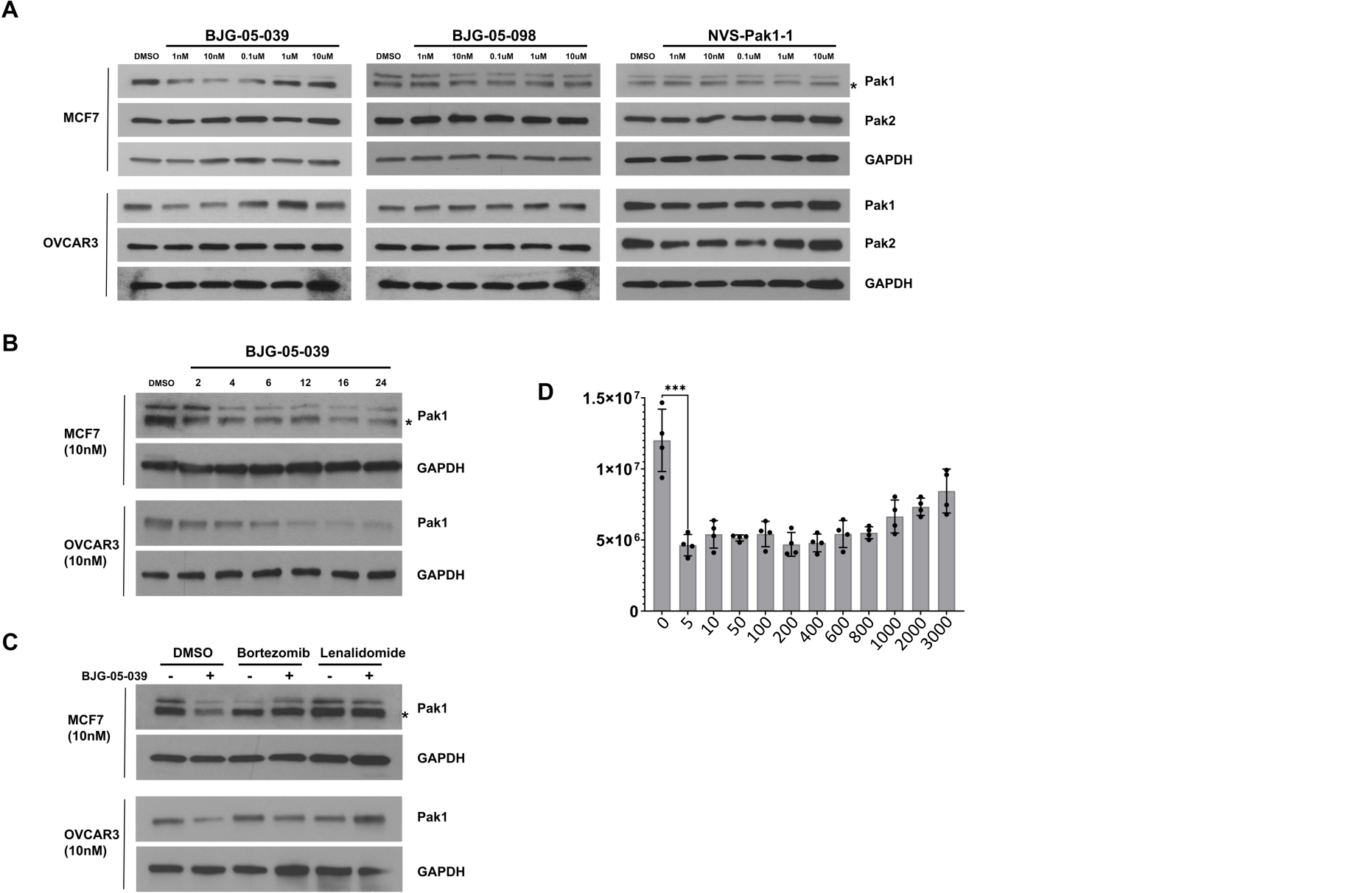
BJG-05-039 Induces Selective Degradation of PAK1 Dependent on CRBN, Neddylation, and the Proteosome. **A)** Effects of BJG-05-039, BJG-05-098, and NVS-PAK1-1 on PAK1 and PAK2 expression. MCF7 and OVCAR3 cells were treated with increasing concentrations of the indicated compounds for 24 h and protein lysates were analyzed by immunoblot. **B)** Time course of PAK1 degradation. **C)** Effect of Bortezomib and Lenalidomide, respectively, on degrader capacity of BJG-05-039. Asterisk indicates PAK1 signal on immunoblot. **D)** Quantitation of PAK1 expression by luminescence assay. HEK293 cells stably expressing near-endogenous levels of Nluc-PAK1 were treated with the indicated concentrations of BJG-05-039. 24 h post treatment the cells were lysed an analyzed for luciferase activity as described in the Methods section.

To assess degrader selectivity across the proteome, MOLT4 cells were treated with 1 µM BJG-05-039 for 5 h and global multiplexed mass spectrometry-based proteomic analysis was performed ^31^. As expected for lenalidomide-based degraders, the ring-finger protein RNF166 as well as other known IMiD off target proteins, Ikaros (IKZF1) and Aiolos (IKZF3), were strongly downregulated by BJG-05-039 treatment (Figure S4) ^32^. Interestingly, while BJG-05-039 showed potent *in vitro* inhibition of PAK1 (IC50 = ss nM) and a ∼50% reduction in PAK1 levels as determined by immunoblot (Figure 2A), significant downregulation of this kinase was not apparent in the proteomic analysis (Table S3).

To better quantify the effect of degrader treatment, we transfected HEK293 cells with an expression vector encoding nano-luciferase (Nluc)-tagged PAK1. The Nluc tag allows for luciferase-based quantitation of protein expression ^33^. Nluc-PAK1 cells were treated with BJG- 05-039 or its N-methylated analog, BJG-05-098 and PAK1 expression was assessed (Figure 2D). This experiment showed that half-maximal degradation of PAK1 was achieved at low nM concentrations of BJG-05-039, with approximately 70% reduction in PAK1 expression following treatment with 10 nM of BJG-05-039.

### BJG-05-039 Exhibits Enhanced Effects on Signaling Compared with NVS-PAK1-1

In certain cell lines PAK1 has well-characterized functions in regulating proliferative signaling. We compared the activity of BJG-05-039 against NVS-PAK1-1 in two such cell lines, OVCAR3 and MCF7. As a readout for anti-PAK activity, we assessed phosphorylation of MEK1 at S298, a direct target site for Group A PAKs ^34^, as well as phosphorylation of the downstream target of MEK, ERK. At a 10 nM dose, BJG-05-039 inhibited phosphorylation of MEK S298, as well as inhibiting phosphorylation of ERK, however an equivalent dose of NVS-PAK1-1 had no visible impact on phosphorylation status of these proteins (Figure 3A). Selective knockdown of PAK1 showed similar effects to BJG-05-039 in these cell lines, substantially reducing levels of phospho-MEK and phospho-ERK, whereas knockdown of PAK2 showed little effect (Figure 3B).

**Figure 3.**
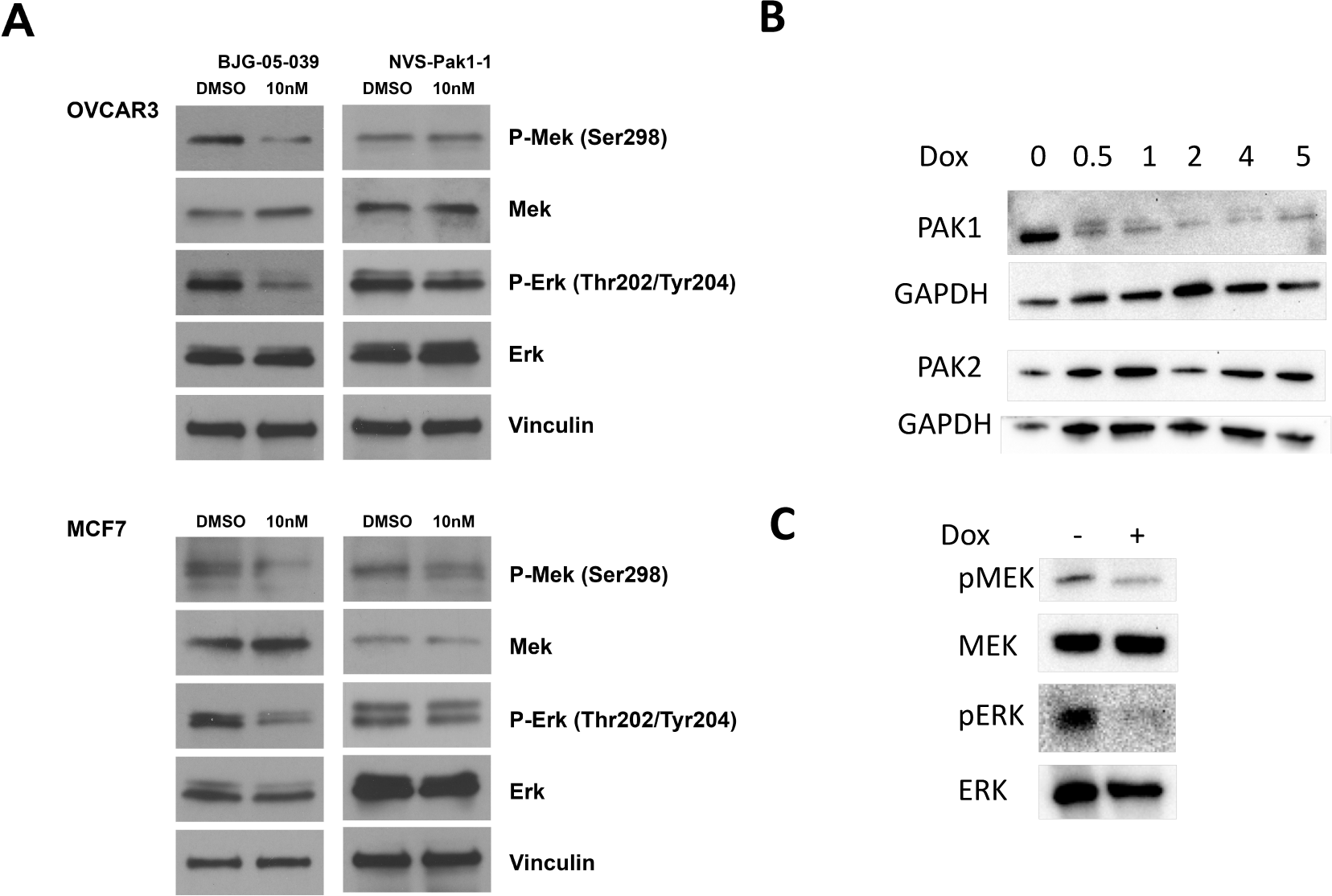
PAK1 Degrader Potently Suppresses Proliferative Signals. **A)** OVCAR3 and MCF7 cells were treated for 24 h with DMSO, 10 nM BJG-05-039, or NVS-PAK1-1 as indicated. MEK and ERK phosphorylation was assessed by immunoblot with the indicated phosphorylation-specific antibodies. **B)** MCF7 cells were stably transduced with a doxycycline-regulated shRNA against PAK1 ^27^. shRNA expression was induced by the indicated amounts of doxycycline (mg/ml) and immunoblots were performed using cell lysates 24 h post doxycycline addition. **C)** MCF7 cells were treated with vehicle or 1 mg/ml doxycycline. Immunoblots were performed using cell lysates 24 h post doxycycline addition.

### BJG-05-039 is Effective in Reducing Proliferation in PAK1-Dependent, but not PAK2- Dependent Cell Lines

We next compared the anti-proliferative effects of PAK1 degradation and inhibition. Using the same two cell lines, we found that BJG-05-039 was far more potent than NVS-PAK1-1, with EC50 values of 3 and 8.4 nM in MCF7 and OVCAR3 cells, respectively, compared to 11.9 μM and 3.68 μM, respectively, for the parental compound NVS-PAK1-1 (Figure 4A and 4B). In contrast, BJG-05-039 did not show increased efficacy relative to NVS- PAK1-1 in OMM1 and HEY-A8 cells, which genome-wide CRISPR-screens suggest are highly dependent on PAK2 (Figure 4A and 4B) (https://depmap.org/portal/gene/PAK2?tab=dependency). These large differences in EC50 values in PAK1-dependent cells are likely to be ddirectly related to the degrader properties of BJG-05- 039, as an isomeric non-degrader control (BJG-05-098) shoed no activity in any of the cell lines (Fig. 4C). In addition, the effects of BJG-05-039 are unlikely to be caused by non-specific degradation of proteins (*e.g*., IKZF1 or IKZF3), as lenalidomide lacked significant antiproliferative effects in either cell line (Figure 4D). These data suggest that the combined selective inhibition and degradation of PAK1 by BJG-05-039 is more potent at reducing proliferative signaling in PAK1-dependent cells than what can be achieved by inhibiting catalytic activity alone. Consistent with this view, NVS-PAK1-1 showed much more potent anti-proliferative effects in cells in which PAK1 levels were reduced ∼50% using shRNA (Figure 4E). These results also support a mechanism of action related to combined PAK1 inhibition/degradation as opposed to “off-target” degradation effects.

**Figure 4.**
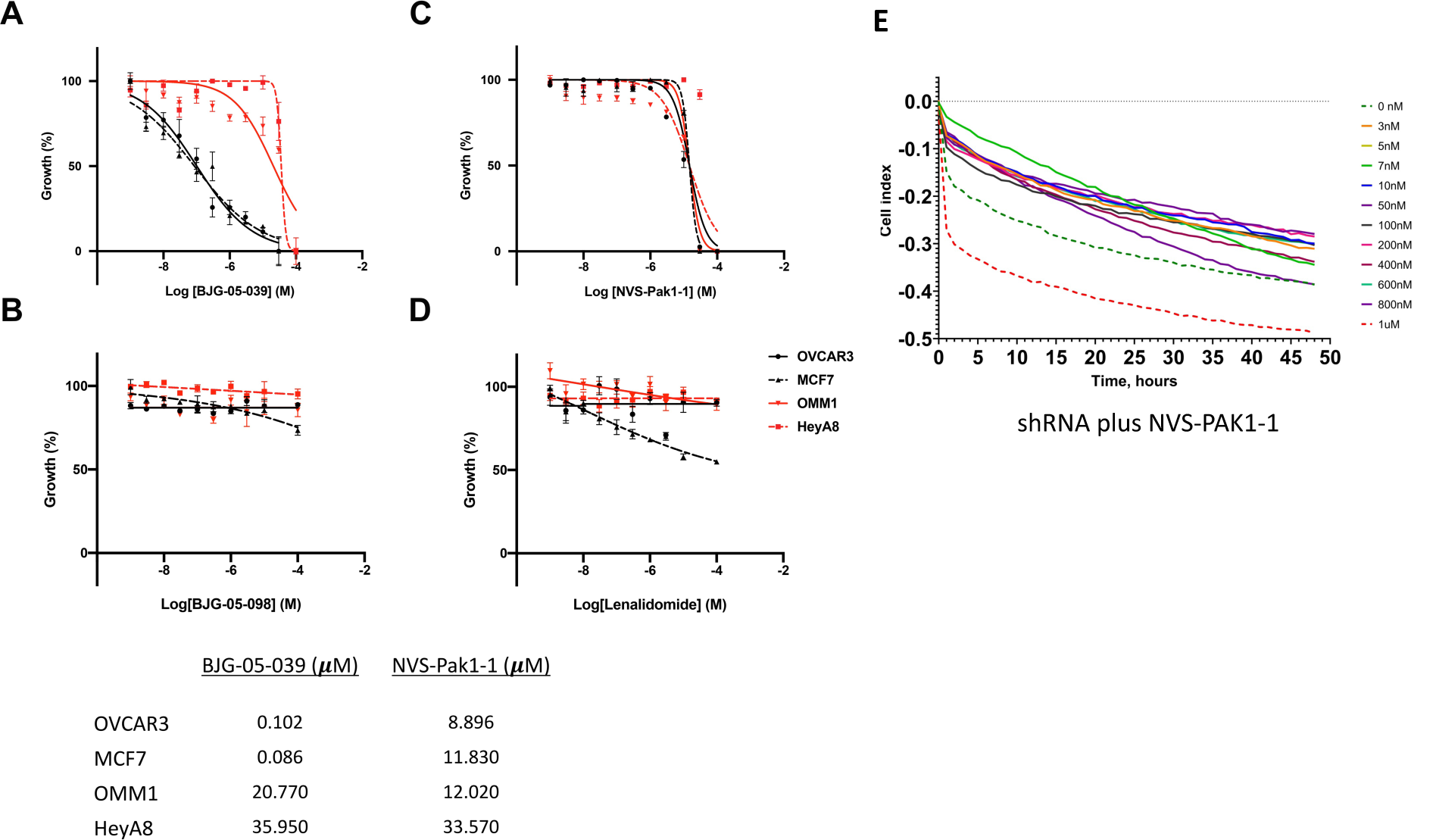
PAK1 Degrader Selectively Suppresses Proliferation of PAK1-Dependent Cells. PAK1-dependent (MCF7 and OVCAR3) and PAK2-dependent (OMM1 and HeyA8) cells were treated for 96 h with varying concentrations of **A)** BJG-05-039, **B)** BJG-05-098, **C)** NVS-PAK1- 1, or **D)** lenalidomide. Cell proliferation was assessed by MTT assay and EC50 values were determined. EC50 values are shown below the graphs. **E)** The effect of reducing PAK1 expression on the potency of NVS-PAK1-1 was assessed by treating control MCF7 cells or MCF7 cells in which PAK1 expression was reduced ∼50% via induction of a PAK1-specific shRNA at 0.5 mg/ml doxycycline.

In many cancer cells, Group A PAKs act as signaling hub, coordinating the activation of various central proliferative, survival, and motility pathways ^2^. As such, there has been interest in targeting these enzymes with small molecule inhibitors ^10, 12, 35^. One pan-PAK inhibitor, PF3758309 ^10^, was evaluated in a phase 1 clinical trial, but poor pharmacologic properties and excessive toxicity led to its withdrawal. More selective inhibitors, such as FRAX-597, FRAX- 1036, and G5555 showed efficacy in cell-based models, in particular, in cells in which the *PAK1* gene was amplified or in which *RAC1* was a driving oncogene ^11–13, 16, 27, 36^. However, progress in clinical development has been hampered by genetic and pharmacologic evidence suggesting a vital role for PAK2 in cardiovascular function and vascular integrity in adult mammals ^14, 18^.

Acute cardiotoxicity upon inhibitor treatment or gene loss is thought to be due to a unique role for PAK2 in regulating ER stress and oxidative stress in cardiomyocytes ^37^.

While the vast majority of Group A PAK inhibitors equally affect PAK1, -2, and -3, NVS- PAK1-1 - an allosteric inhibitor that binds beneath the αC helix rather than in the hinge region of the ATP binding pocket of PAK1 - exhibits an approximately 50-fold specificity for PAK1 over PAK2. This unique specificity prompted us to ask if we could improve upon this molecule by imparting it with the ability to degrade its target. As with FAK degraders ^38^, PAK1 degraders have the potential to incite more benefit than standard enzymatic inhibitors because removal of PAK1 would not only reduce its kinase enzymatic activity, but also its scaffolding function, both of which mediate signaling activity. For example, PAK1 has been shown to be required for AKT activation ^15, 39^, but these effects map to the N-terminus of PAK1 and appear to be independent of kinase activity ^39^. Degraders also offer the potential for prolonged efficacy, which is driven by target half-life, and this is an important consideration given the short half-life of NVS-PAK1-1 ^20, 21^. While adding the linker and lenalidomide moiety to NVS-PAK-1 reduced the potency of BJG-05-039 as a PAK1 catalytic inhibitor, it also increased its selectivity over PAK2 more than ten-fold (Table S1). Given the necessity for isoform selective targeting, this improved selectivity for PAK1 could reduce the potential toxicity of this molecule relative to the parental NVS- PAK1-1.

While we were not able to achieve more than 70% loss of the PAK1 protein using BJG-05- 039, the combination of catalytic inhibition and lowered protein expression had dramatically improved inhibitory effects on signaling and proliferation in PAK1-dependent cell lines (Figures 3A and 4A). In this respect, the degrader may be mimicking the effects of genetic manipulations such as RNAi-mediated gene knock down. This property can be used, in conjunction with the parental inhibitor NVS-PAK1-1, to tease apart kinase *vs.* scaffolding effects of PAK1 in cells. In fact, our data suggests that reducing the total level of PAK1 expression synergizes with catalytic expression, as evidenced by the increased potency of the non-degrader NVS-PAK-1 when used in conjunction with partial knockdown of PAK1 with shRNA (Figure 4C). Thus, in addition to its potential role in reigniting interest in clinical development of PAK inhibitors, the degrader compound described herein provides a useful tool compound for signaling analysis.

## CONCLUSION

Given their role in regulating the ERK, AKT, and b-catenin pathways, Group A PAKs have been considered as potential therapeutic targets in cancer. However, the PAK2 isoform plays a key role in normal cardiovascular function in adult mammals, and this factor has impeded further preclinical development of anti-PAK agents. Selective PAK2-sparing molecules present a potential path forward and we previously showed that one such inhibitor, NVS-PAK1-1, has promising effects in a mouse model of NF2. Given that PAK1 has significant scaffolding activity in addition to its catalytic activity, we considered that a degrader based on NVS-PAK1-1 might provide considerable benefits while avoiding toxicities associated with PAK2 inhibition. Here, we show that an optimized heterobifunctional degrader, BJG-05-039, promoted rapid and selective degradation of PAK1 but not PAK2, and had more potent anti-proliferative effects than NVS-PAK-1. Given that many cancer cell types display elevated PAK1 expression and activity, and knockdown of PAK1 has been shown to have considerable anti-proliferative and/or anti-survival effects in these settings, isoform-specific inhibition and degradation of PAK1 may reignite interest in targeting this kinase in cancer.

## EXPERIMENTAL SECTION

### Chemical Synthesis

Detailed information on chemical synthesis analytical data as well as supplementary data can be found in the Supporting Information.

### Cell Lines

MCF7 (female, CVCL_0031) and HEK293 cells were cultured in high-glucose DMEM medium (Gibco) supplemented with 10% fetal bovine serum (HyClone), 2 mM L-glutamine and 100U/ml penicillin/streptomycin (Gibco). MOLT4 (male, CVCL_0013) were cultured in RPMI 1640 medium (Gibco) supplemented with 10% fetal bovine serum and 2 mM L-glutamine. OVCAR3 (female, CVCL_0465), Hey A8 (female, CVCL_8878) and OMM1 (male, CVCL_6939) were cultured in RPMI 1640 medium (Gibco) supplemented with 10% fetal bovine serum, 2 mM L-glutamine and 100U/ml penicillin/streptomycin (Gibco). All cell lines were cultured at 37℃ in a humidified 5% CO2 incubator. HEK293 cells stably expressing Nluc- PAK1 were constructed by transfecting with the pFN31K Nluc-PAK1 expression vector (0.5 *μ*g DNA per well) in 12-well plates using Lipofectamine 3000 (Invitrogen) according to the manufacturer’s protocol. Transfected Nluc-PAK1 cells were cultured for 1 week in DMEM medium containing G418 (2 mg ml^−1^) to select stable clones.

A doxycycline (Dox)-inducible shRNA-bearing retrovirus against PAK1 was previously described^27^ and oligonucleotide used in this study are as follows: PAK1 shRNA-1 5′-GAT CCCCGAAGAGAGGTTCAGCTAAATTCAAGAGATTTAGCTGAACCTCTCTTCTTTTTTGGAAA-3′; Recombinant viruses were generated using the Phoenix amphotropic packaging system (Orbigen). the ΦNX cells were transfected using Lipofectamine 3000 (Invitrogen). Viral supernatants were harvested 72 hr post-transfection and filtered. Ovarian (OVCAR3) and breast (MCF7) cancer cells were incubated with retroviral supernatant supplemented with 8 μg/ml polybrene for 4 h at 37°C, and then were cultured in growth media for 48 h for viral integration. Green fluorescent protein (GFP)-positive infected cells were selected by flow cytometry.

### Plasmids

The pFN31K-Nluc-PAK1 vector was constructed as follows: The gene sequence encoding PAK1 from pCMV6M-PAK1 (Plasmid #12209 Addgene) was PCR-amplified using the following oligonucleotide pair: TTCTGGCGGGCTCGAGCGTCGACATGGAACAGAAACT (Forward), TACCGAGCCCGAATTGAATTCCTCGAGGCCACGAAG (Reverse), designed with recognition sites for XhoI and EcoRI restriction enzymes. The PCR product was subcloned into the expression vector pFN31K-Nluc) using the XhoI and EcoRI restriction endonucleases (NEB, USA) and In-Fusion HD Enzyme (Takara, Japan).

### Retroviral transductions

The ΦNX packaging cell line (Orbigen) was transfected using Lipofectamine 2000 (ThermoFisher Scientific) according to the manufacturer’s instruction.

Viral supernatants were harvested 48 hours post-transfection and filtered. Cells were incubated with retroviral supernatant supplemented with 4 μg/ml polybrene for 4 hours at 37°C, and then were cultured in growth media for 48 hours for viral integration. Green fluorescent protein (GFP)-positive infected cells were selected by flow cytometry.

### Drug Treatment with PAK1 Degraders

NVS-PAK1-1, BJG-05-039, BJG-05-098, bortezomib (Selleckchem), and lenalidomide (Selleckchem) were dissolved in DMSO at 10 mM. Cells were seeded in 6-well plate at 250,000 cells per mL in 2 mL per well. Cells were incubated overnight then treated with various concentrations of PAK degraders alone or together with bortezomib or lenalidomide for 24 hours. Protein lysates were harvested at the times specified.

### Immunoblotting

Cells were washed once in PBS and lysed in RIPA buffer (50 mM Tris-HCl, 150 mM NaCl, 0.25% (w/v) sodium deoxycholate, 1% (w/v) NP-40, pH 7.5) containing protease inhibitor cocktail (Roche) and phosphatase inhibitor cocktail (Roche) for 30 minutes on ice. Cell lysates were obtained by centrifugation at 13,500 rpm for 15 minutes at 4°C. BCA protein assay (Thermo Scientific) was used to determine protein concentration then equal amounts of total proteins were separated by SDS-PAGE (Bio-Rad) and transferred to PVDF membrane (Thermo Scientific) at 100V for 2 hours. Membranes were blocked in 5% (w/v) non-fat dry milk in tris-buffered saline with 0.1% Tween-20 (TBS-T) for 1 hour and incubated with primary antibody at 4°C for overnight. Membranes were washed with TBS-T on the next day and incubated with HRP-conjugated secondary antibodies (Millipore) at room temperature for 1 hour and exposed to films after washing.

| Antibodies | Company | Catalog | Dilution |
| --- | --- | --- | --- |
| Rabbit PAK1 | Cell Signaling Technology | 2602 | 1:1000 |
| Rabbit PAK2 | Cell Signaling Technology | 2608 | 1:1000 |
| Rabbit MEK | Cell Signaling Technology | 9122 | 1:1000 |
| Rabbit Phospho-MEK | Cell Signaling Technology | 9128 | 1:1000 |
| Rabbit ERK | Cell Signaling Technology | 4695 | 1:2500 |
| Rabbit Phospho-ERK | Cell Signaling Technology | 9101 | 1:2500 |
| Rabbit GAPDH | Cell Signaling Technology | 2118 | 1:2500 |
| Mouse Vinculin | Sigma-Aldrich | V9131 | 1:2000 |

### Cell viability assay

Cells were plated at 2 x 10^3^ cells per well in 96-well plates overnight and treated with various concentrations of Pak degraders for 72 hours. Cell viability was measured by MTT assay and the half maximal inhibitory concentration (IC50) was calculated using GraphPad Prism. Triplicates were performed for each sample and medium alone was used as a blank.

### Luciferase Assays

HEK293 cells stably expressing pFN31K-Nluc-PAK1 were assayed for luciferase activity according to the manufacturer’s Nano-Glo® Live Cell Assay System protocol (Promega). In brief, 25μl of Nano-Glo Live Cell Reagent was added per well and the plate was gently mixed by hand, then placed in a 37°C luminometer for 10 minutes.

### KinomeScan

The kinase engagement assay (KINOMEscan) was performed by DiscoverX assessing binding abilities toward a set of protein kinases. NVS-PAK1-1 was screened at a concentration of 1 μM and BJG-05-039 was screened at a concentration of 10 μM.

### Kinase Activity Assay

Kinase activity assays were performed by Reaction Biology Corp.

Compounds were tested in 10-dose IC50 duplicate mode with a 3-fold serial dilution starting at 1 μM. The control compound, staurosporine, was tested in 10-dose IC50 mode with 4-fold serial dilution starting at 20 μM. Reactions were carried out at 10 μM ATP. IC50 values were calculated using Prism 7.0 (GraphPad)

### Biochemical Assay Protocol for PAK1 and PAK2

The activity/inhibition of human recombinant PAK1 (kinase domain) or PAK2 (full length) was determined by measuring the phosphorylation of a FRET peptide substrate (Ser/Thr19) labeled with Coumarin and Fluorescein using Z’-LYTE™ assay (Invitrogen). The 10 µL assay mixtures contained 50 mM HEPES (pH 7.5), 0.01% Brij-35, 10 mM MgCl2, 1 mM EGTA, 2 µM FRET peptide substrate, and PAK enzyme (20 pM PAK1; 50 pM PAK2). Incubations were carried out at 22°C in black polypropylene 384-well plates (Corning Costar). Prior to the assay, enzyme, FRET peptide substrate and serially diluted test compounds were preincubated together in assay buffer (7.5 µL) for 10 minutes, and the assay was initiated by the addition of 2.5 µL assay buffer containing 4x ATP (160 µM PAK1; 480 µM PAK2). Following the 60-minute incubation, the assay mixtures were quenched by the addition of 5 µL of Z’-LYTE™ development reagent, and 1 hour later the emissions of Coumarin (445 nm) and Fluorescein (520 nm) were determined after excitation at 400 nm using an Envision plate reader (Perkin Elmer). An emission ratio (445 nm/520 nm) was determined to quantify the degree of substrate phosphorylation.

### TMT LC-MS Sample Preparation

MOLT4 cells were treated with DMSO in biological triplicate and 1 µM BJG-05-039 for 5 h and harvested by centrifugation. Cell lysis was performed by the addition of Urea buffer (8 M Urea, 50 mM NaCl, 50 mM 4-(2-hydroxyethyl)- 1-piperazineethanesulfonic acid (EPPS) pH 8.5, Protease and Phosphatase inhibitors) followed by manual homogenization by 20 passes through a 21-gauge (1.25 in. long) needle. Lysate was clarified by centrifugation at 4 °C and protein quantified using bradford (Bio-Rad) assay. 100 µg of protein for each sample was reduced, alkylated and precipitated using methanol/chloroform as previously described ^31^. The resulting precipitated protein was resuspended in 4 M Urea, 50 mM HEPES pH 7.4, buffer for solubilization, followed by dilution to 1 M urea with the addition of 200 mM EPPS, pH 8. Proteins were digested for 12 hours at room temperature with LysC (1:50 ratio), followed by dilution to 0.5 M urea and a second digestion step was performed by addition of trypsin (1:50 ratio) for 6 hours at 37 °C. Anhydrous ACN was added to each peptide sample to a final concentration of 30%, followed by addition of Tandem mass tag (TMT) reagents at a labelling ratio of 1:4 peptide:TMT label. TMT labelling occurred over a 1.5 h incubation at room temperature followed by quenching with the addition of hydroxylamine to a final concentration of 0.3%. Each of the samples were combined using adjusted volumes and dried down in a speed vacuum followed by desalting with C18 SPE (Sep-Pak, Waters). The sample was offline fractionated into 96 fractions by high pH reverse-phase HPLC (Agilent LC1260) through an aeris peptide xb-c18 column (phenomenex) with mobile phase A containing 5% acetonitrile and 10 mM NH_4_HCO_3_ in LC-MS grade H_2_O, and mobile phase B containing 90% acetonitrile and 5 mM NH4HCO3 in LC-MS grade H_2_O (both pH 8.0). The resulting 96 fractions were recombined in a non-contiguous manner into 24 fractions and desalted using C18 solid phase extraction plates (SOLA, Thermo Fisher Scientific) followed by subsequent mass spectrometry analysis.

Data were collected using an Orbitrap Fusion Lumos mass spectrometer (Thermo Fisher Scientific, San Jose, CA, USA) coupled with an Proxeon EASY-nLC 1200 LC lump (Thermo FIsher Scientific, San Jose, CA, USA). Peptides were separated on a 50 cm 75 μm inner diameter EasySpray ES903 microcapillary column (Thermo Fisher Scientific). Peptides were separated over a 190 min gradient of 6 - 27% acetonitrile in 1.0% formic acid with a flow rate of 300 nL/min.

Quantification was performed using a MS3-based TMT method as described previously. The data were acquired using a mass range of m/z 340 – 1350, resolution 120,000, AGC target 5 x 105, maximum injection time 100 ms, dynamic exclusion of 120 seconds for the peptide measurements in the Orbitrap. Data dependent MS2 spectra were acquired in the ion trap with a normalized collision energy (NCE) set at 35%, AGC target set to 1.8 x 104 and a maximum injection time of 120 ms. MS3 scans were acquired in the Orbitrap with HCD collision energy set to 55%, AGC target set to 2 x 105, maximum injection time of 150 ms, resolution at 50,000 and with a maximum synchronous precursor selection (SPS) precursors set to 10.

### LC-MS data analysis

Proteome Discoverer 2.4 (Thermo Fisher Scientific) was used for .RAW file processing and controlling peptide and protein level false discovery rates, assembling proteins from peptides, and protein quantification from peptides. The MS/MS spectra were searched against a Swissprot human database (December 2019) containing both the forward and reverse sequences. Searches were performed using a 20 ppm precursor mass tolerance, 0.6 Da fragment ion mass tolerance, tryptic peptides containing a maximum of two missed cleavages, static alkylation of cysteine (57.02146 Da), static TMT labelling of lysine residues and N-termini of peptides (304.2071 Da), and variable oxidation of methionine (15.99491 Da). TMT reporter ion intensities were measured using a 0.003 Da window around the theoretical m/z for each reporter ion in the MS3 scan. The peptide spectral matches with poor quality MS3 spectra were excluded from quantitation (summed signal-to-noise across channels < 100 and precursor isolation specificity < 0.5), and the resulting data was filtered to only include proteins with a minimum of 2 unique peptides quantified. Reporter ion intensities were normalized and scaled using in-house scripts in the R framework ^41^. Statistical analysis was carried out using the limma package within the R ^42^.

### Cell Viability and Luciferase Assays

All experiments were performed at least three times. Results were reported as means ± SD. The significance of the data was determined by two-tailed, unpaired Student’s *t-*test with *p* < 0.05 considered statistically significant.

### Data and code availability

The accession number for the proteomic data reported in this paper in Pride: PXcccc

## ASSOCIATED CONTENT

### Supporting Information

Additional tables and figures illustrating PAK1/NVS-PAK1-1 interaction, profiles of NVS-based (allosteric) degraders, KinomeSCAN data, degrader effects on total proteome, and biochemical data. This material is available free of charge via the Internet at http://pubs.acs.org

## AUTHOR INFORMATION

### Corresponding Author

*. Phone: (215) 728-5319. Fax: (215) 728-3616.

### Author Contributions

J.C. and N.S.G conceived of the study. B.J.G., S.T., J.H, K.A.D, and E.S.F. designed and synthesized the compounds, profiled the degrader against the kinome, and/or carried out the proteomic analysis in MOLT4 cells. S.K., B.A., and R.D, performed the structural analysis shown in Figure 1. H.-Y.C., S.K, and G.A, designed and conducted the biological profiling of the degraders.

### Notes

N.S.G. is a founder, science advisory board member (SAB) and equity holder in Syros, C4, Allorion, Jengu, B2S, Inception, EoCys, CobroVentures (advisor), GSK (advisor), Larkspur (board member) and Soltego (board member). The Gray lab receives or has received research funding from Novartis, Takeda, Astellas, Taiho, Jansen, Kinogen, Arbella, Deerfield, Springworks, Interline and Sanofi. K.A.D is a consultant to Kronos Bio. E.S.F. is a founder, science advisory board member (SAB) and equity holder in Civetta Therapeutics, Jengu Therapeutics (board member) and Neomorph Inc. An advisor for EcoR1 capital, Deerfield, Sanofi and Avilar Therapeutics. The Fischer lab receives or has received research funding from Novartis, Astellas, Ajax, Voronoi, Interline and Deerfield.

## ACKNOWLEDGMENTS

We thank Joachim Rudolph and Christopher Heise (Genentech) for data regarding the specificity of NVS-PAK1-1. This work was supported by grants from the NIH (J.C. CA227184 and CA148805; N.S.G and E.S.F. CA218278), and NCI Core Grant P30 CA06927 to FCCC.

## ABBREVIATIONS USED

CDI, carbonyldiimidazole; CRBN, cereblon; DCM, dichloromethane; DIPEA, diisopropylethylamine; DMF, N,N-dimethylformamide; DMP, dess-martin periodane; DMSO, dimethyl sulfoxide; EtOAc, ethyl acetate; HATU, hexafluorophosphate azabenzotriazole tetramethyl uronium; HPLC, high-performance liquid chromatography; MeCN, acetonitrile; MeOH, methanol; MeI, methyl iodide, Pd2(dba)3, tris(dibenzylideneacetone)dipalladium(0); PAK, p21-activated kinase; PROTAC, Proteolysis Targeting Chimera; TEA, triethylamine; TFA, trifluoroacetic acid; UPLC-MS, ultra-performance liquid chromatography-mass spectrometry; XPhos, 2-dicyclohexylphosphino-2′,4′,6′-triisopropylbiphenyl

## KEY RESOURCES TABLE

| REAGENT or RESOURCE | SOURCE | IDENTIFIER |
| --- | --- | --- |
| Antibodies |  |  |
| Rabbit monoclonal anti PAK1 | Cell Signaling Technology | Cat# 2602; 1:1000 |
| Rabbit monoclonal anti PAK2 | Cell Signaling Technology | Cat# 2608; 1:1000 |
| Rabbit polyclonal anti phospho-PAK1/2/3 (S141) | Invitrogen | Cat# 44940G; 1:1000 |
| Rabbit anti ERK1/2 | Cell Signaling Technology | Cat# 4695; 1:2500 |
| Rabbit anti phospho-ERK1/2 (T202/Y204) | Cell Signaling Technology | Cat# 9101; 1:2500 |
| Rabbit anti MEK1/2 | Cell Signaling Technology | Cat# 9122; 1:1000 |
| Rabbit anti phospho-MEK1 (S298) | Cell Signaling Technology | Cat# 9128; 1:1000 |
| Rabbit anti GAPDH | Cell Signaling Technology | Cat# 2118; 1:2500 |
| Chemicals, Cloning Materials, Plasmids, Peptides, and Recombinant Proteins |  |  |
| DMSO | Fisher | Cat# BP231-100 |
| Bortezomib (PS-341) | Selleckchem | Cat# S1013 |
| Lenalidomide (CC-5013) | Selleckchem | Cat# S1029 |
| NVS-PAK1-1 | Sigma-Aldrich | Cat# SML1867 |
| Valspodar | Sigma-Aldrich | Cat# SML0572 |
| DMEM Medium | Gibco | Cat# 10566-016 |
| RPMI Medium | Gibco | Cat# 72400-047 |
| Fetal Bovine Serum | HyClone | Cat# SH30071.03 |
| Penicillin Streptomycin | Gibco | Cat# 15150-122 |
| PAK1 Protein | This study | NA |
| PAK2 Protein | This study | NA |
| Doxycycline Hyclate | Sigma-Aldrich | Cat# D9891-1G |
| Viral Boost Reagent | Alstem | Cat# VB100 |
| Lipofectamine™ 3000 | Invitrogen | Cat# L3000001 |
| Lipofectamine™ RNAiMAX | Invitrogen | Cat# 13778100 |
| ON-TARGETplus HUMAN PAK1 (5062) siRNA-SMARTpool | Dharmacon | Cat# L-003521-00-005 |
| ON-TARGETplus HUMAN PAK2 (5058) siRNA-SMARTpool | Dharmacon | Cat# L-003597-00-0005 |
| ON-TARGETplus Non-Targeting Pool | Dharmacon | Cat# D-001810-10-05 |
| EcoRI restriction enzyme | NEB | Cat# R3101 |
| XhoI restriction enzyme | NEB | Cat# R0146 |
| In-Fusion HD Enzyme | Takara | Cat# 639649 |
| Stellar competent cells | Takara | Cat# 636766 |
| pFN31K-Nluc | Promega | Cat# N1321 |
| pFN31K-Nluc-PAK1 | This study | NA |
| pLenti-BFP | <sup>40</sup> | NA |
| pCMV6M-Pak1 | Addgene | Cat# 12209 |
| Nano-Glo® Endurazine™ Live Cell Substrate | Promega | Cat# N2570/1/2 |
| Critical Commercial Assays |  |  |
| Z-Lyte (PAK1) | Invitrogen | PV2830 |
| Z-Lyte (PAK2) | Invitrogen | PV4565 |

| Deposited Data |  |  |
| --- | --- | --- |
| BGY-05-039 Proteomics in MOLT4 cells | This study | Pride: PXccccc |
| Experimental Models: Cell Lines |  |  |
| MOLT4 | ATCC | CRL-1582; RRID: CVCL_0013 |
| HEK293 | ATCC | CRL-1573; RRID: CVCL_0031 |
| OVCAR3 | ATCC | HTB-161; RRID: CVCL_0465 |
| MCF7 | ATCC | HTB-22; RRID: CVCL_0031 |
| HeyA8 | MDACC | N/A |
| OMM1 | Hans Grossniklaus, Emory University School of Medicine | N/A |
| Software and Algorithms |  |  |
| GraphPad Prism | GraphPad Software, Inc. | www.graphpad.com/ |
| Adobe Illustrator | Adobe Creative Cloud | https://www.adobe.com/creativecloud.html |
| Proteome Discoverer 2.2 | Thermo Fisher Scientific | https://www.thermofisher.com/order/catalog/product/OPTION-30795 |
| R Framework | Team RCR: A Language and Environment for Statistical Computing | http://www.R-project.org/ |
| Python 3.7.3 | Python Software Foundation | https://www.python.org/ |

**Figure S1.**
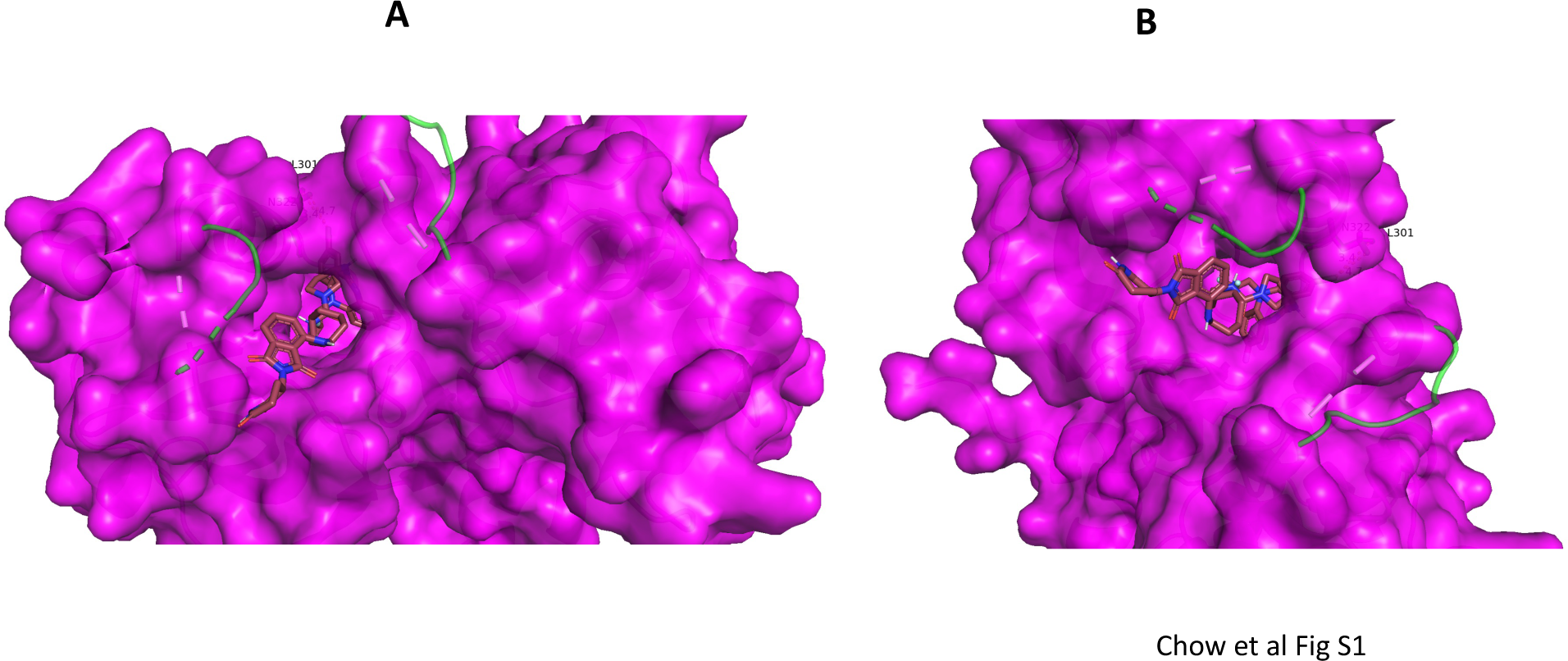
Surface representations of PAK1 and BJG-05-039 docked into the kinase. For clear demonstrations that the ligand occupies ATP pocket region of PAK1, S1A and S1B were taken from 90-degree apart angles. All figures were visualized in PyMOL 2.3.5

**Figure S2.**
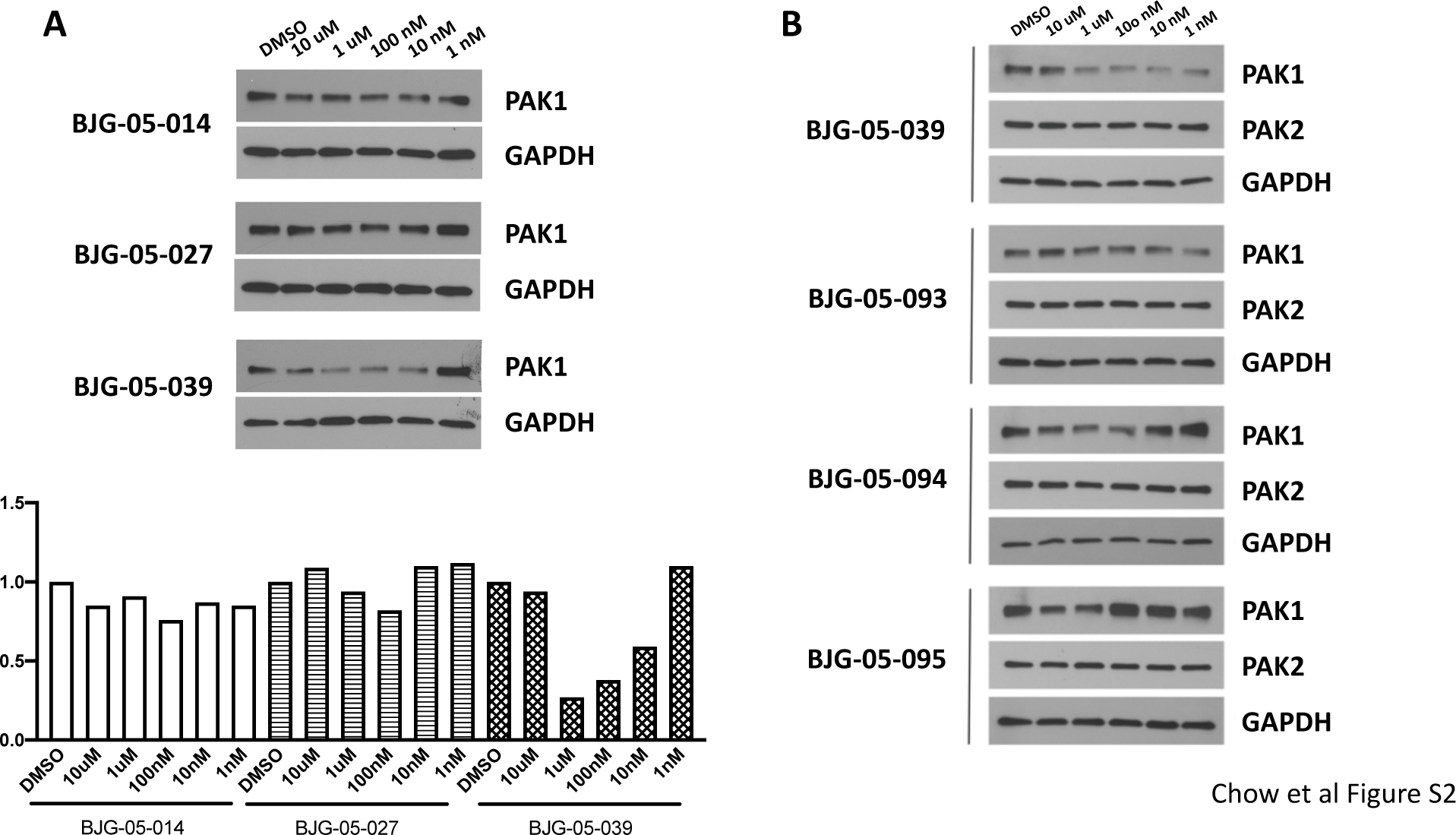
A) Profiling NVS-based (allosteric) degraders BJG-05-014, BJG-05-027, and BJG- 05-039 via western blots in Panc1 cells. Quantification is shown below. B) Profiling NVS-based (allosteric) degrader BJG-05-039 and ATP-competitive degraders BJG-05-093, BJG-05-094, and BJG-05-095 via western blots in Panc1 cells.

**Figure S3.**
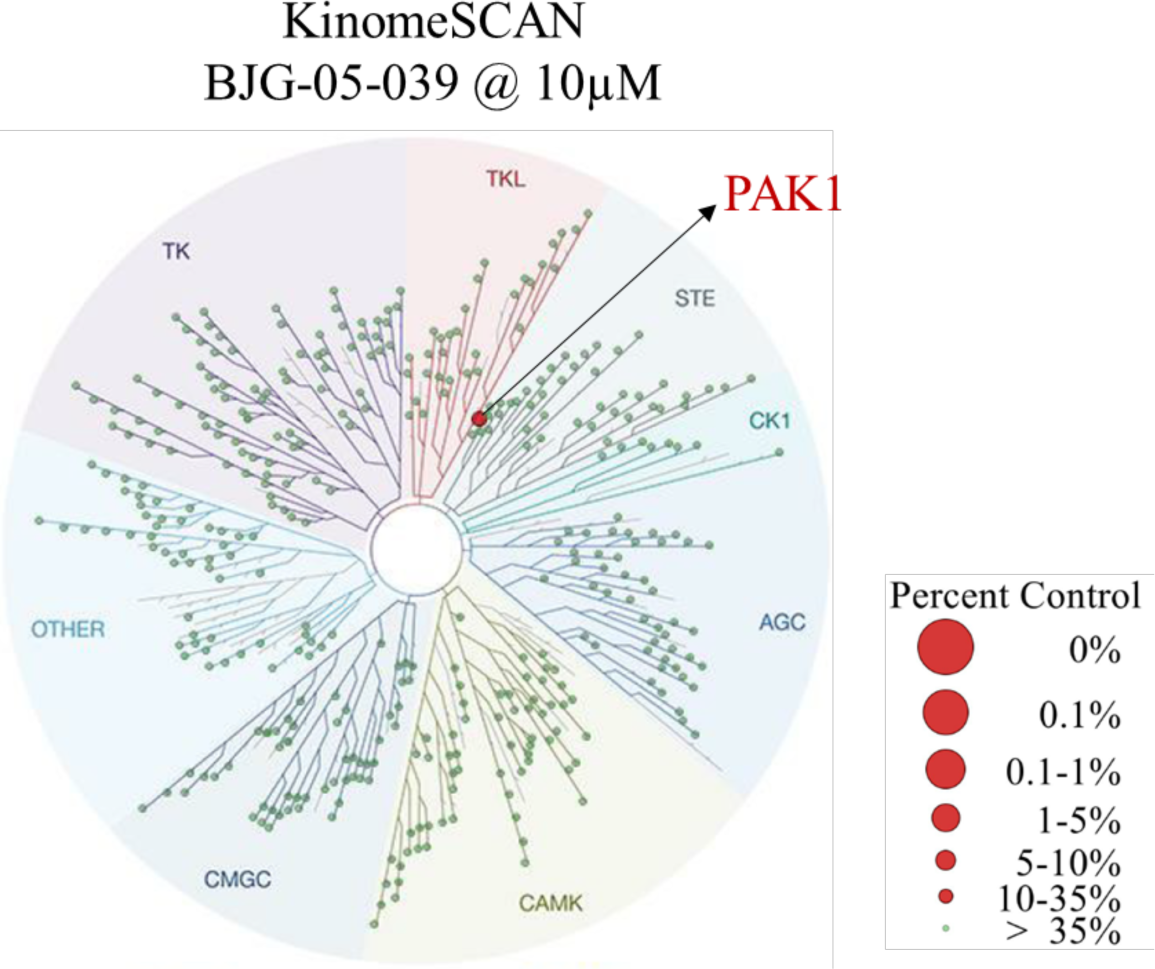
KINOMEscan Profiling of BJG-05-039 @ 10μM. Image generated using TREEspot Software Tool and reprinted with permission from KINOMEscan, a division of DiscoveRx Corporation.

**Figure S4.**
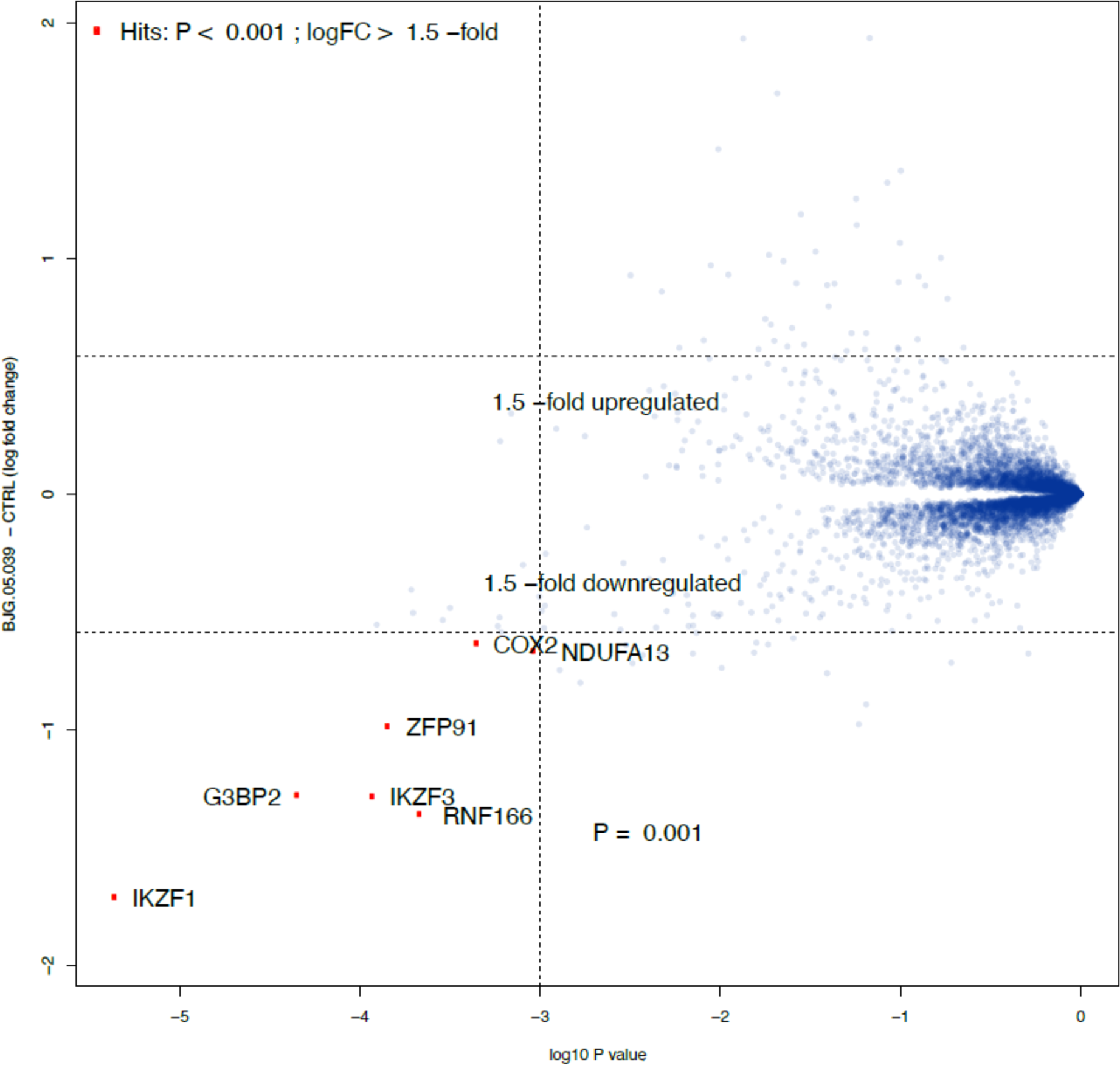
Effect of BJG-05-039 on Proteome. Scatterplot depicts the change in relative protein abundance of MOLT cells treated with BJG-05-039 (5 h, @ 1μM) compared with DMSO vehicle control-treated cells. Protein abundance measurements were made using tandem mass spectrometry and significant changes were assessed by moderated t test as implemented in the limma package (Ritchie et al., 2015). The log2 fold change (log2 FC) is shown on the y-axis and negative log10 p value (-log10 p value) on the x-axis for three independent biological replicates of each treatment.

**Table S1A.** Biochemical IC50 data of allosteric PAK1 degraders.

| Compound | R | PAK1 IC<sub>50</sub> (nM) | PAK2 IC<sub>50</sub> (nM) | PAK4 IC<sub>50</sub> (nM) |
| --- | --- | --- | --- | --- |
| NVS-PAK-1 |  | 29.6 | 824 | >10000 |
| BJG-05-014 |  | 432 | 724 | 152 |
| BJG-05-022 |  | 1490 | 6750 | 3860 |
| BJG-05-019 |  | 6300 | >10000 | >10000 |
| BJG-05-017 |  | >1110 | >3330 | >1000 |
| BJG-05-018 |  | >3330 | >10000 | >10000 |
| BJG-05-027 |  | 594 | >10000 | >10000 |
| BJG-05-039 |  | 233 | >10000 | >10000 |
| BJG-05-020 |  | >10000 | >10000 | >10000 |
| BJG-05-052 |  | n/a | n/a | n/a |
| BJG-05-053 |  | n/a | n/a | n/a |
| BJG-05-098 |  | n/a | n/a | n/a |
| BJG-05-026 |  | n/a | n/a | n/a |

**Table S1B.** Biochemical IC50 data of ATP-competitive PAK1 degraders.

| Compound | R | PAK1 IC<sub>50</sub> (nM) | PAK2 IC<sub>50</sub> (nM) | PAK4 IC<sub>50</sub> (nM) |
| --- | --- | --- | --- | --- |
| BJG-05-093 |  | 32.5 | n/a | n/a |
| BJG-05-094 |  | 103 | n/a | n/a |
| BJG-05-095 |  | 36.6 | n/a | n/a |

**Table S2.**
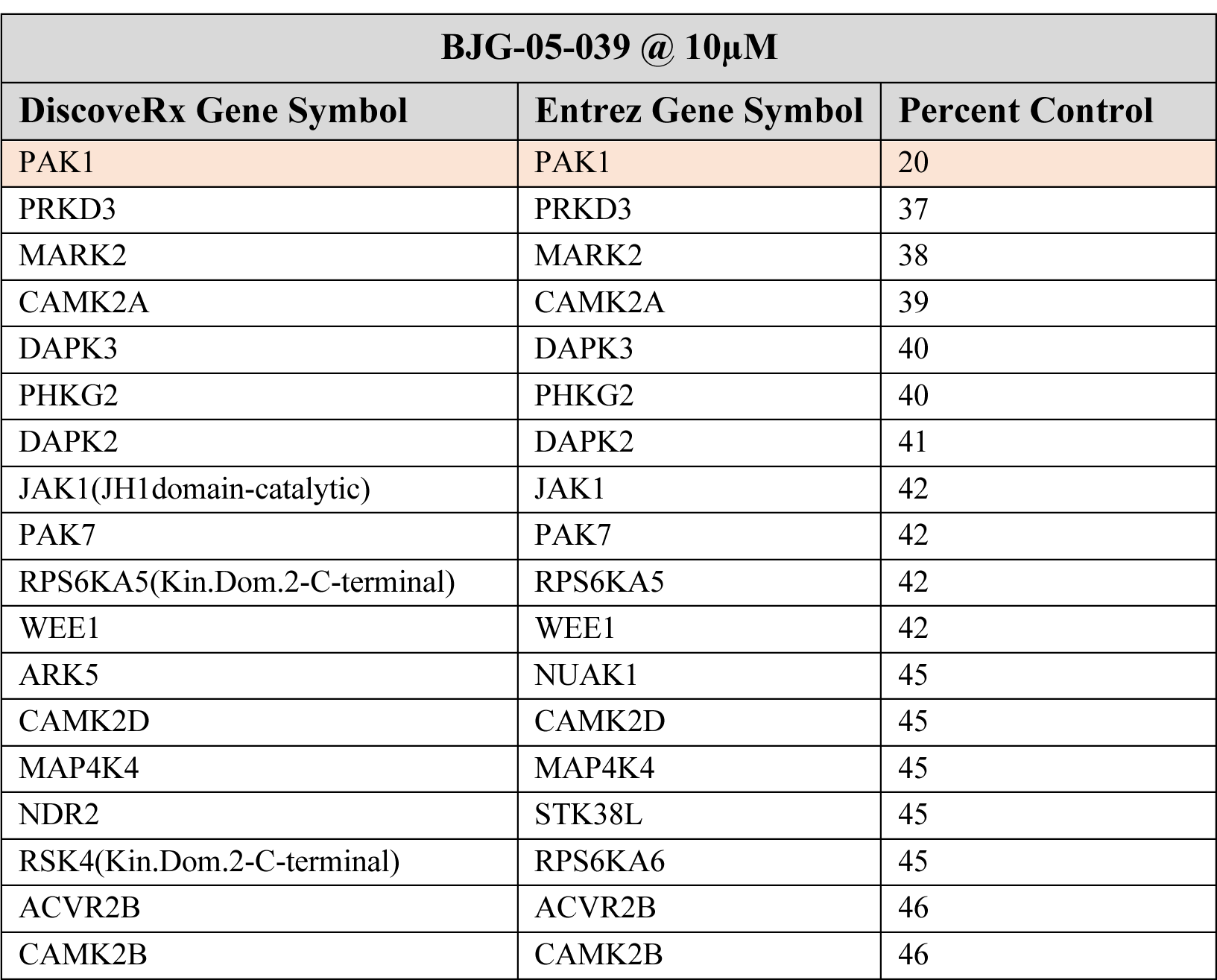
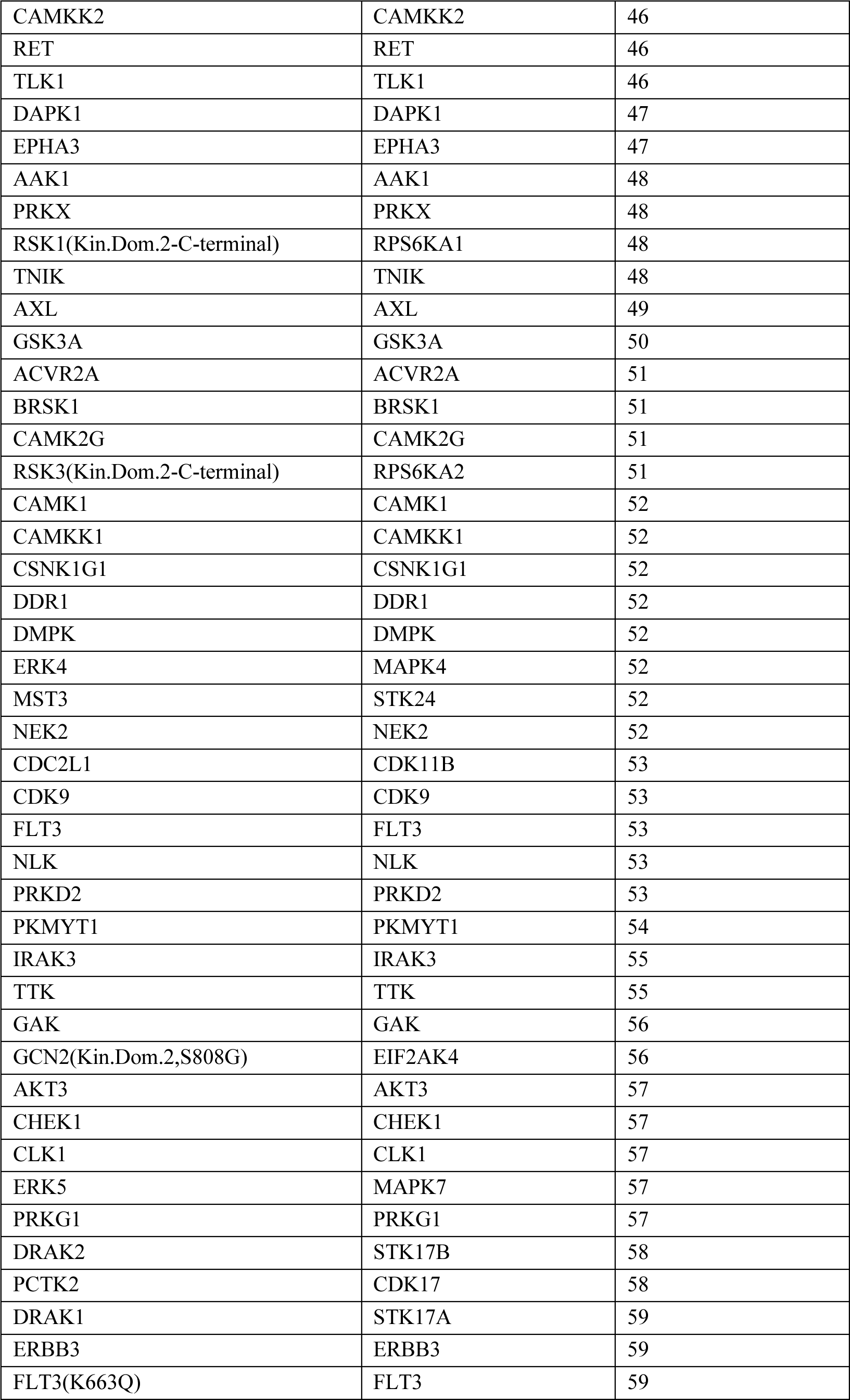
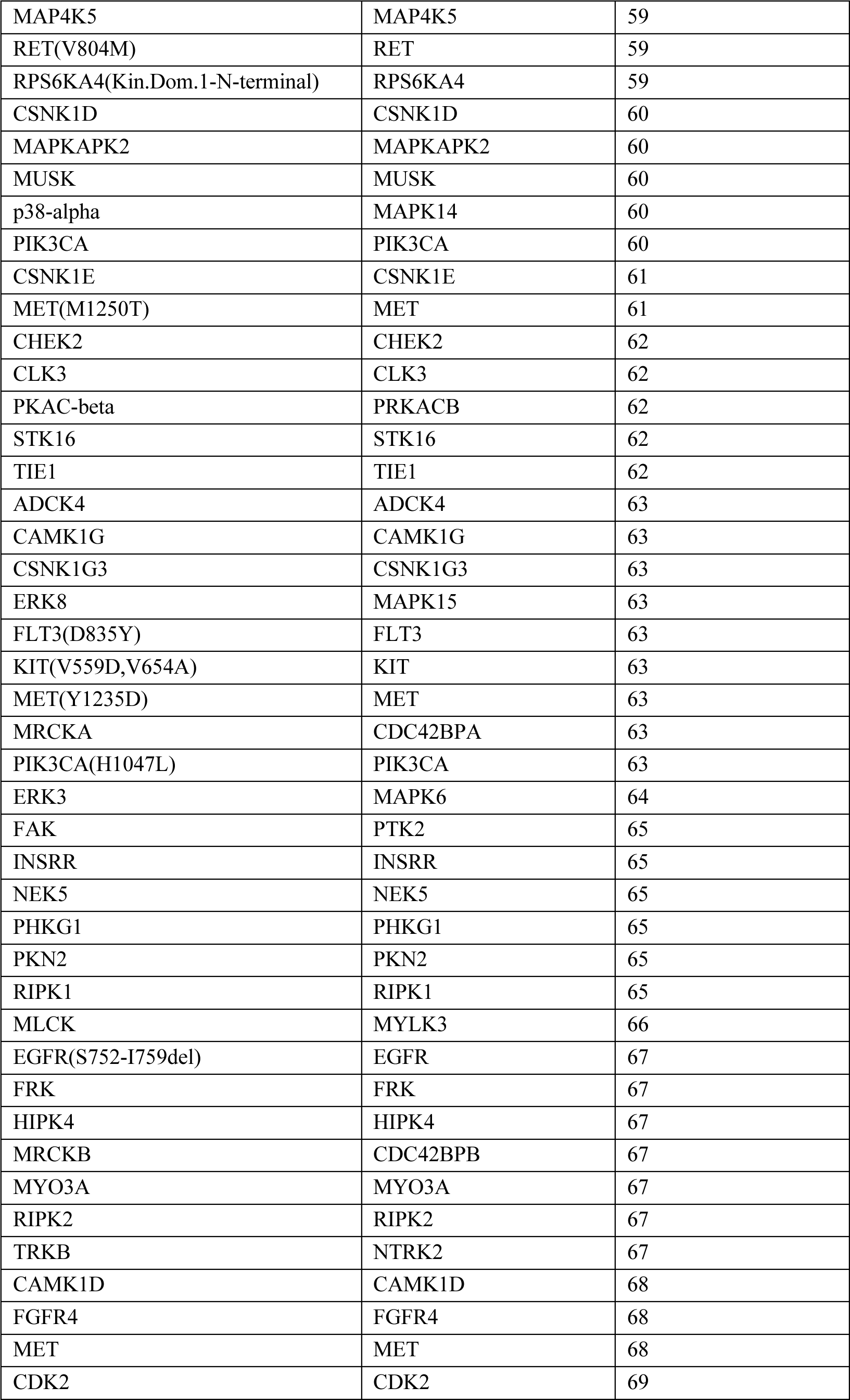
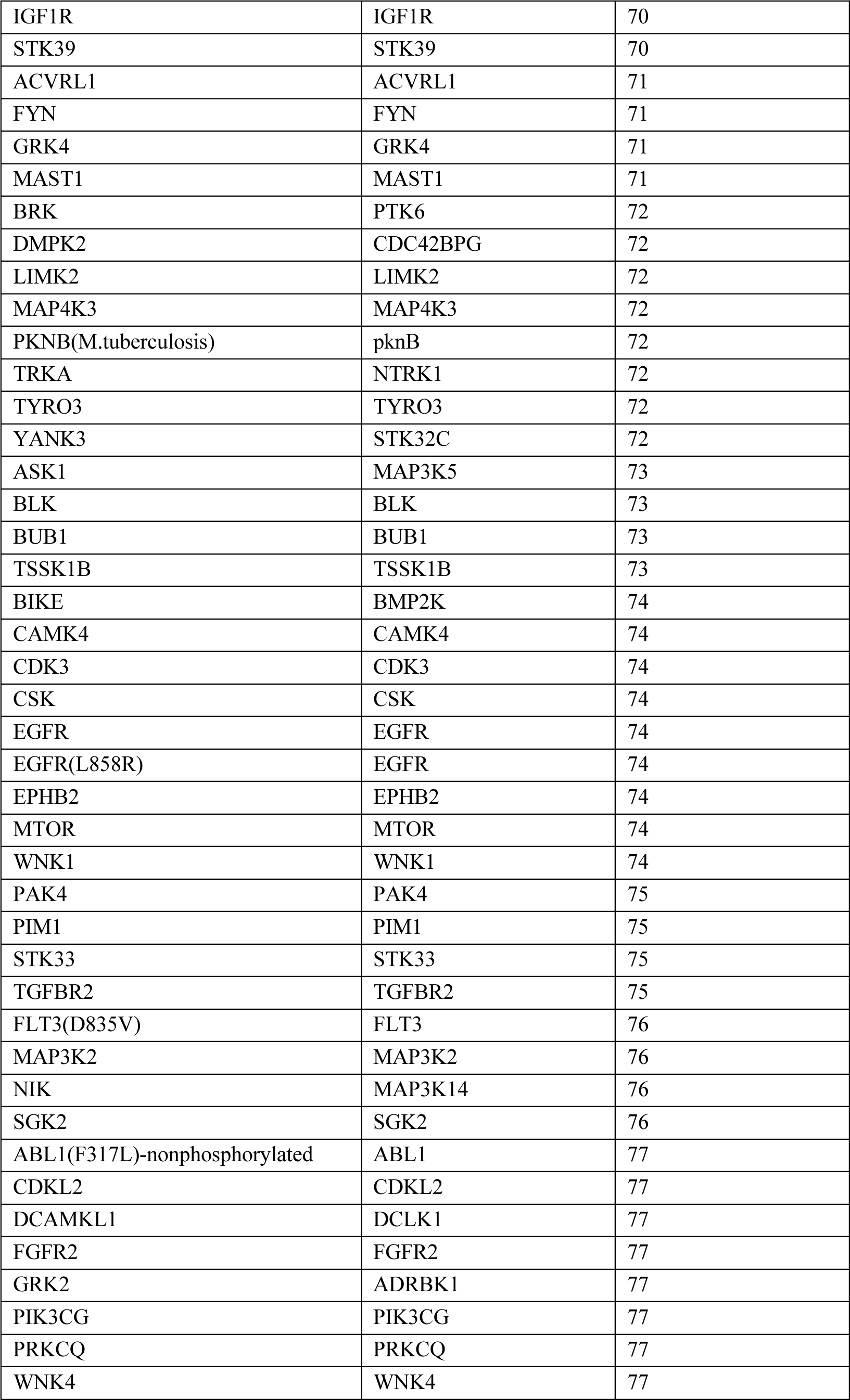
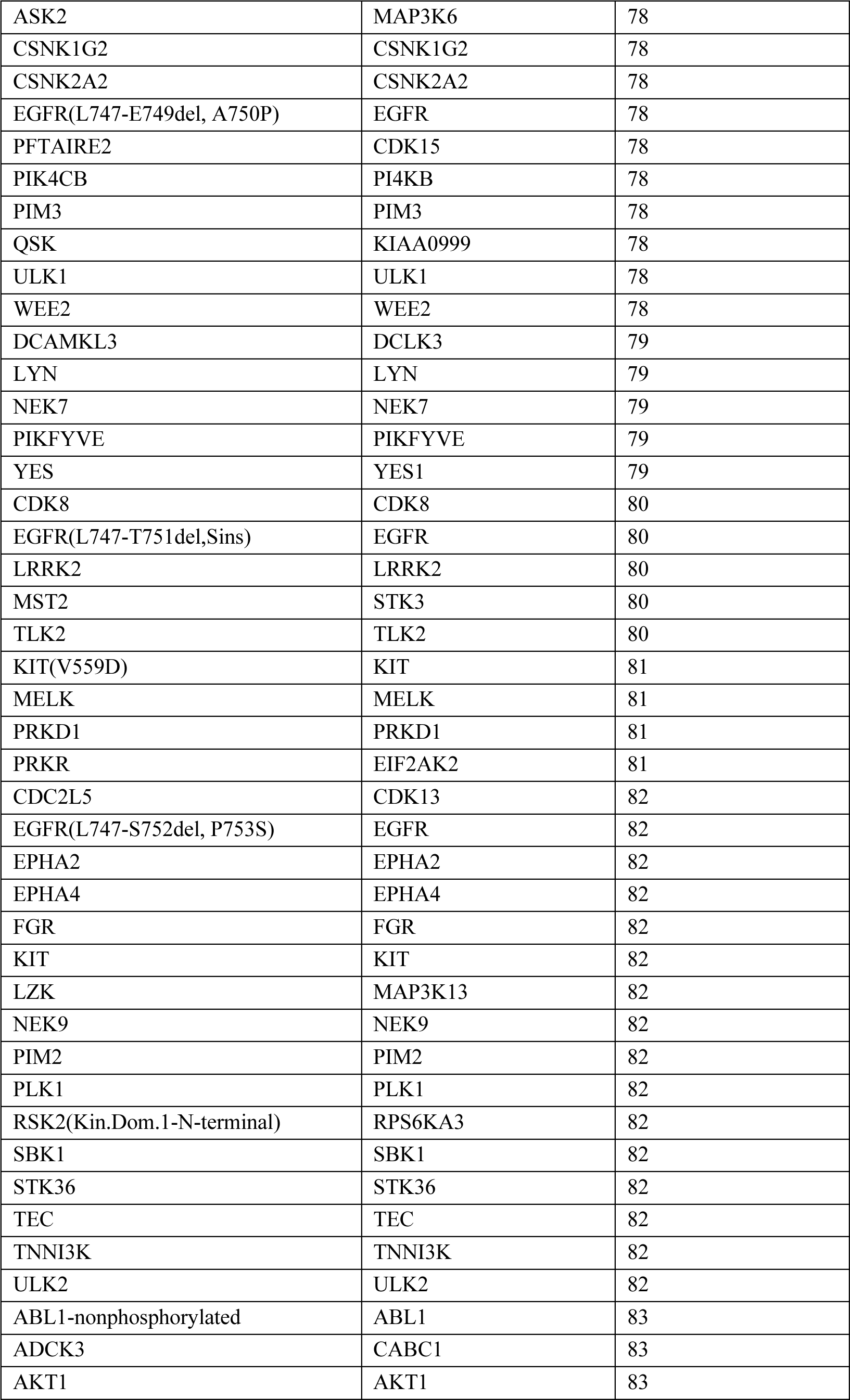
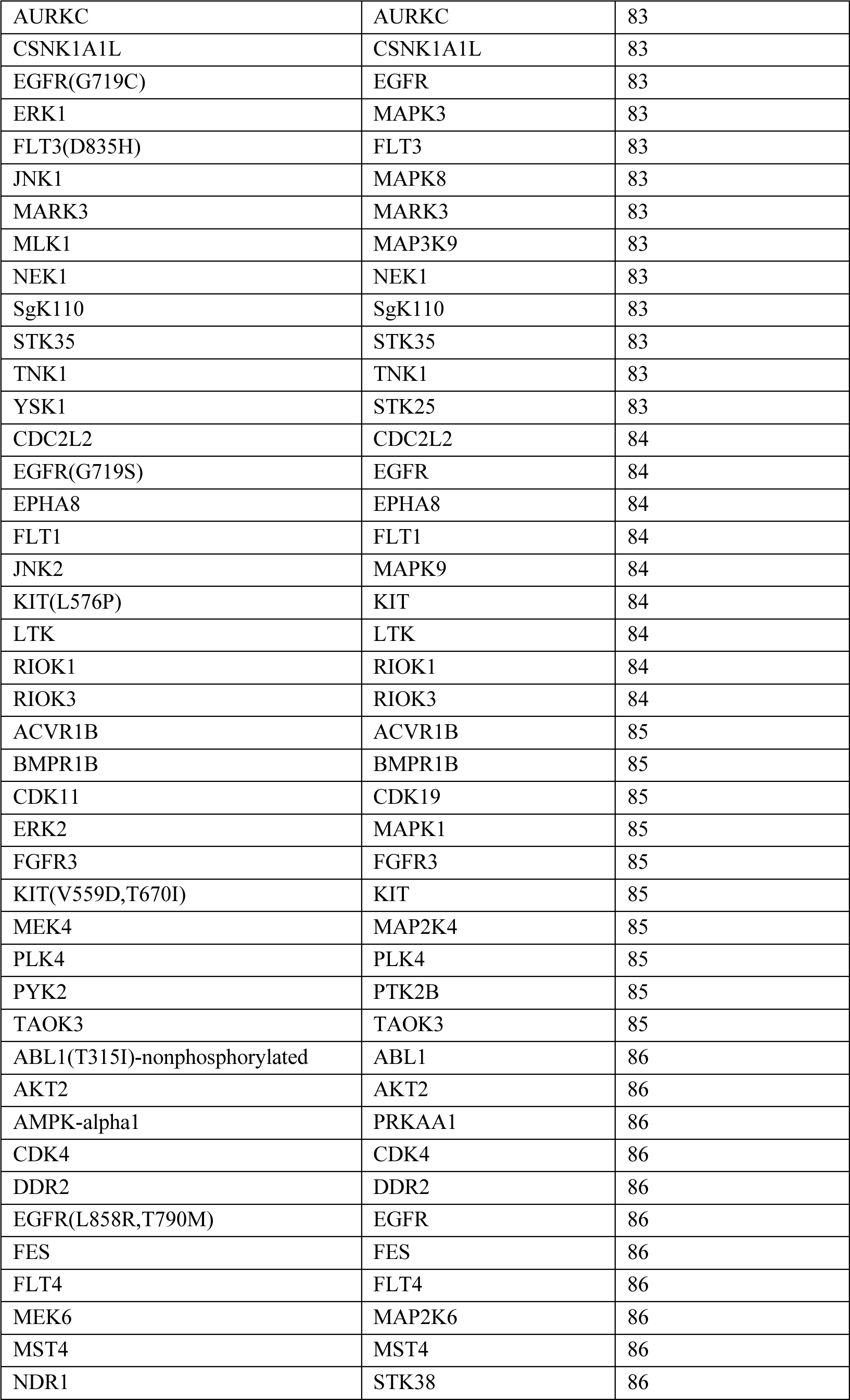
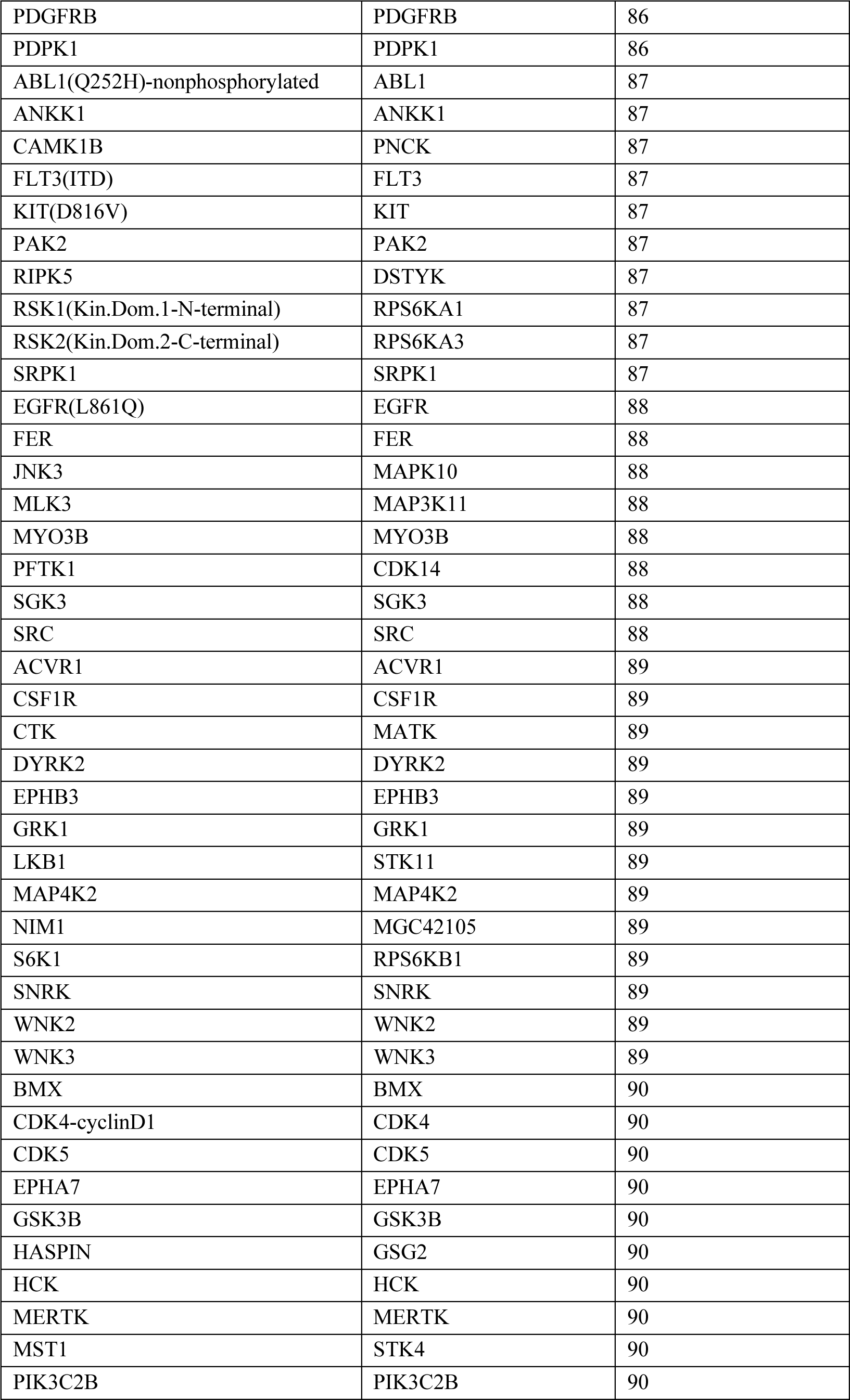
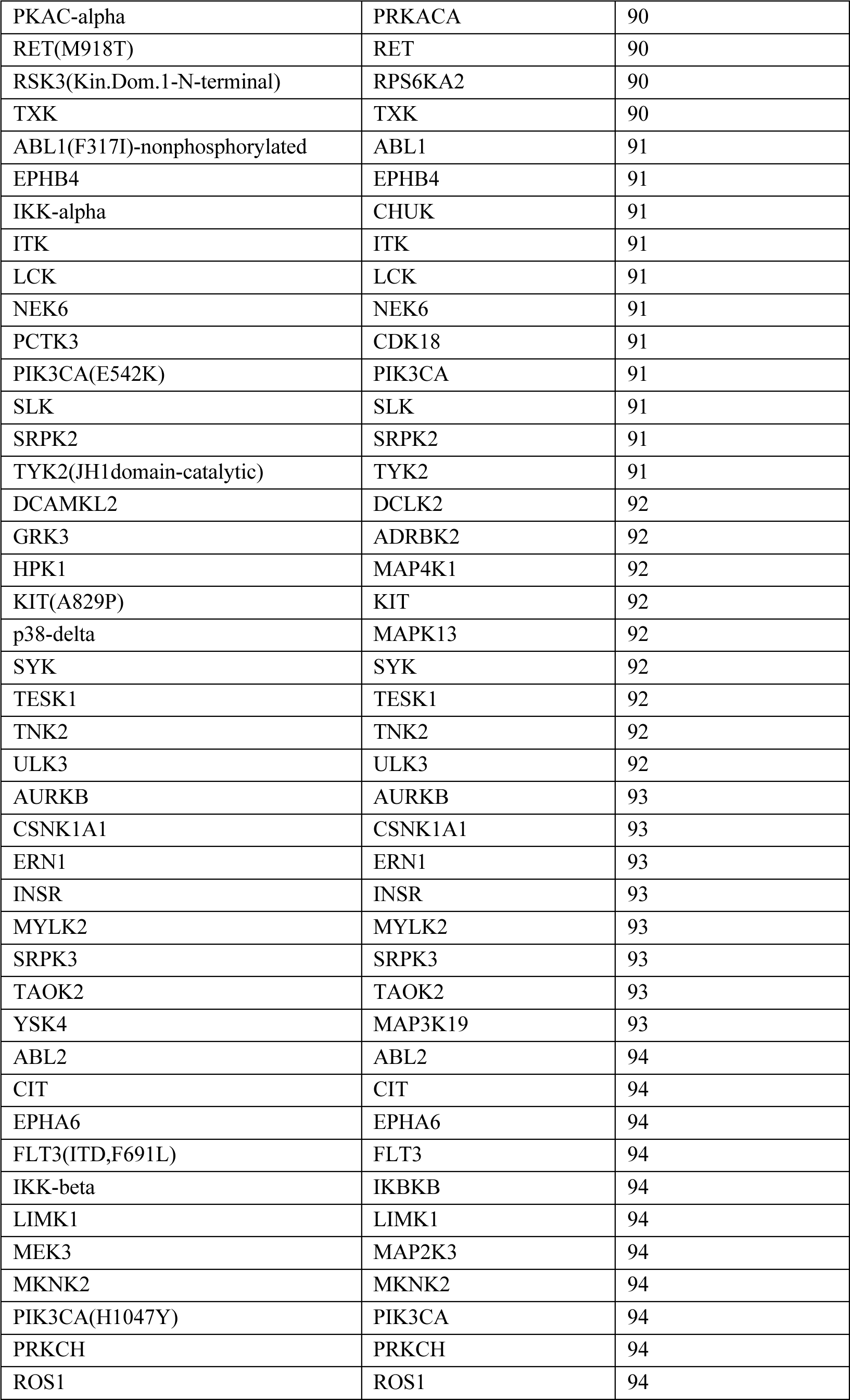
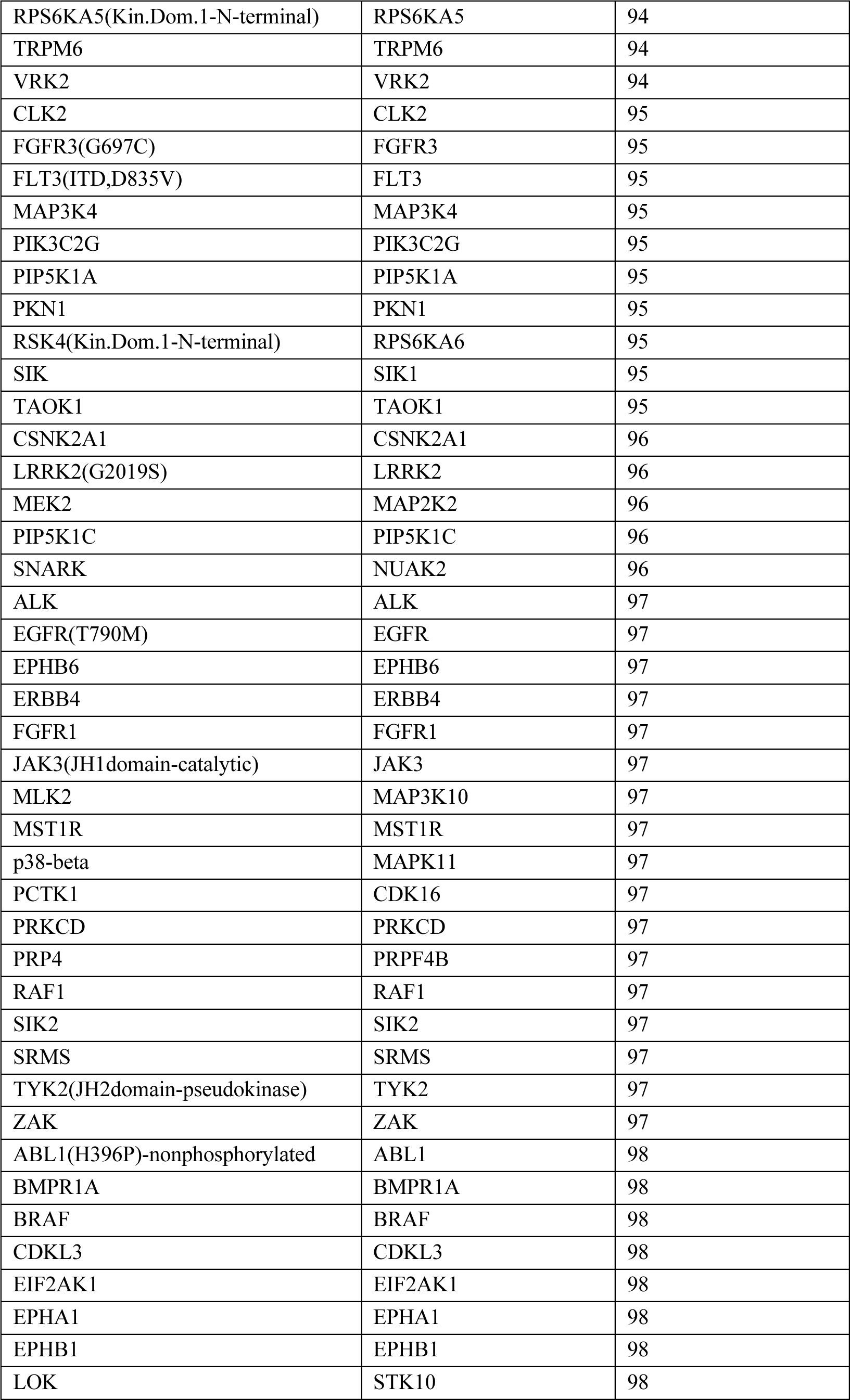
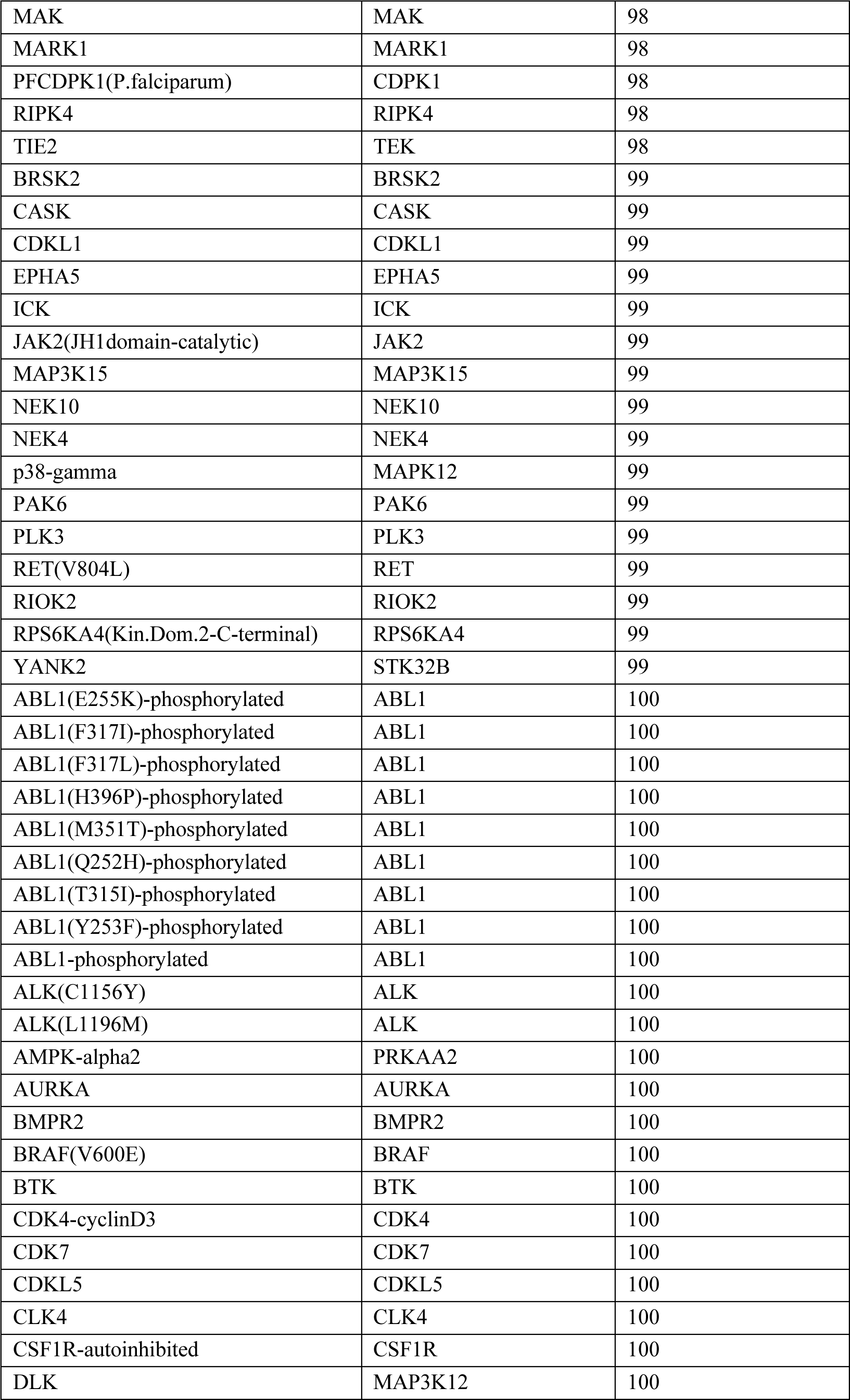
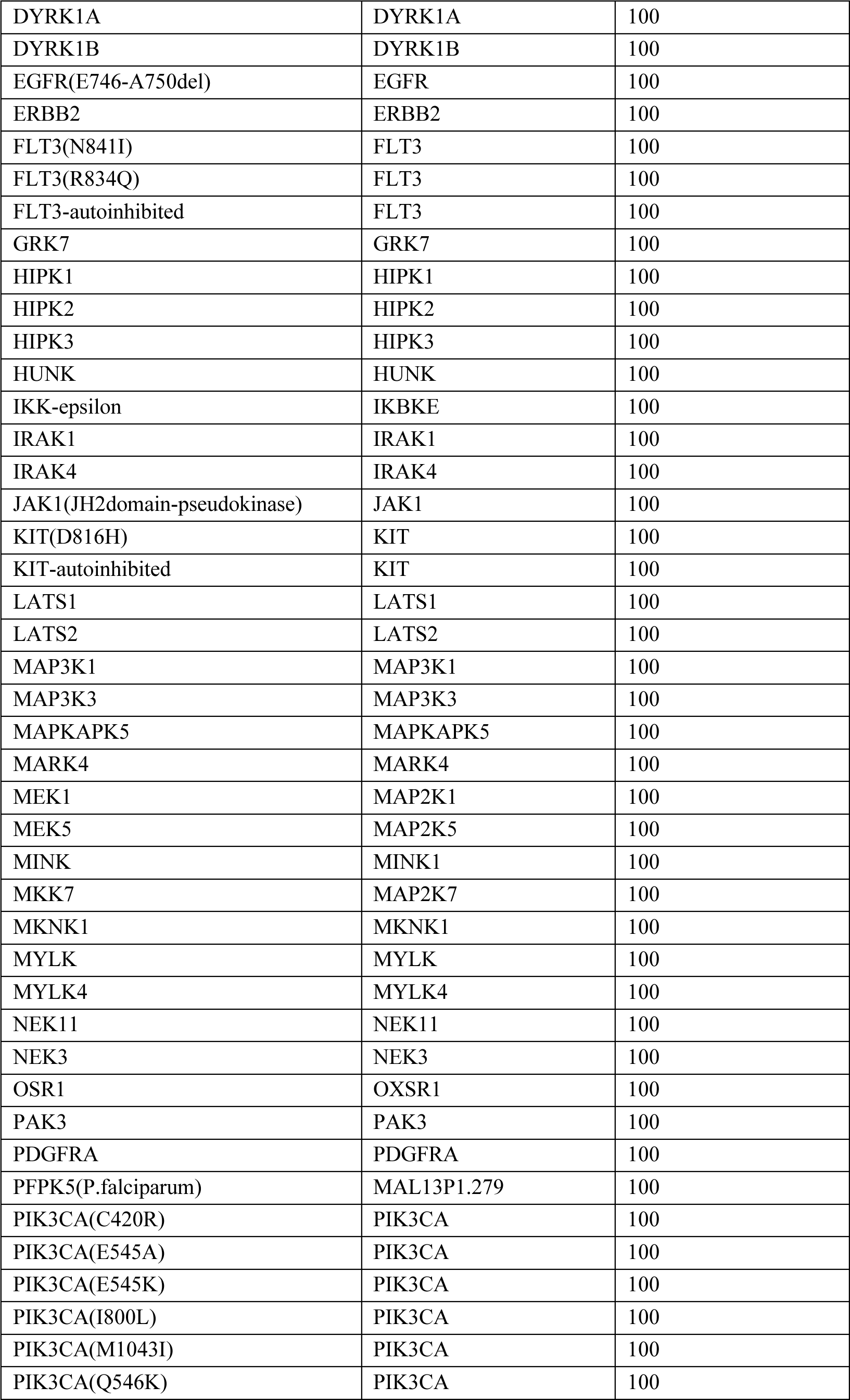
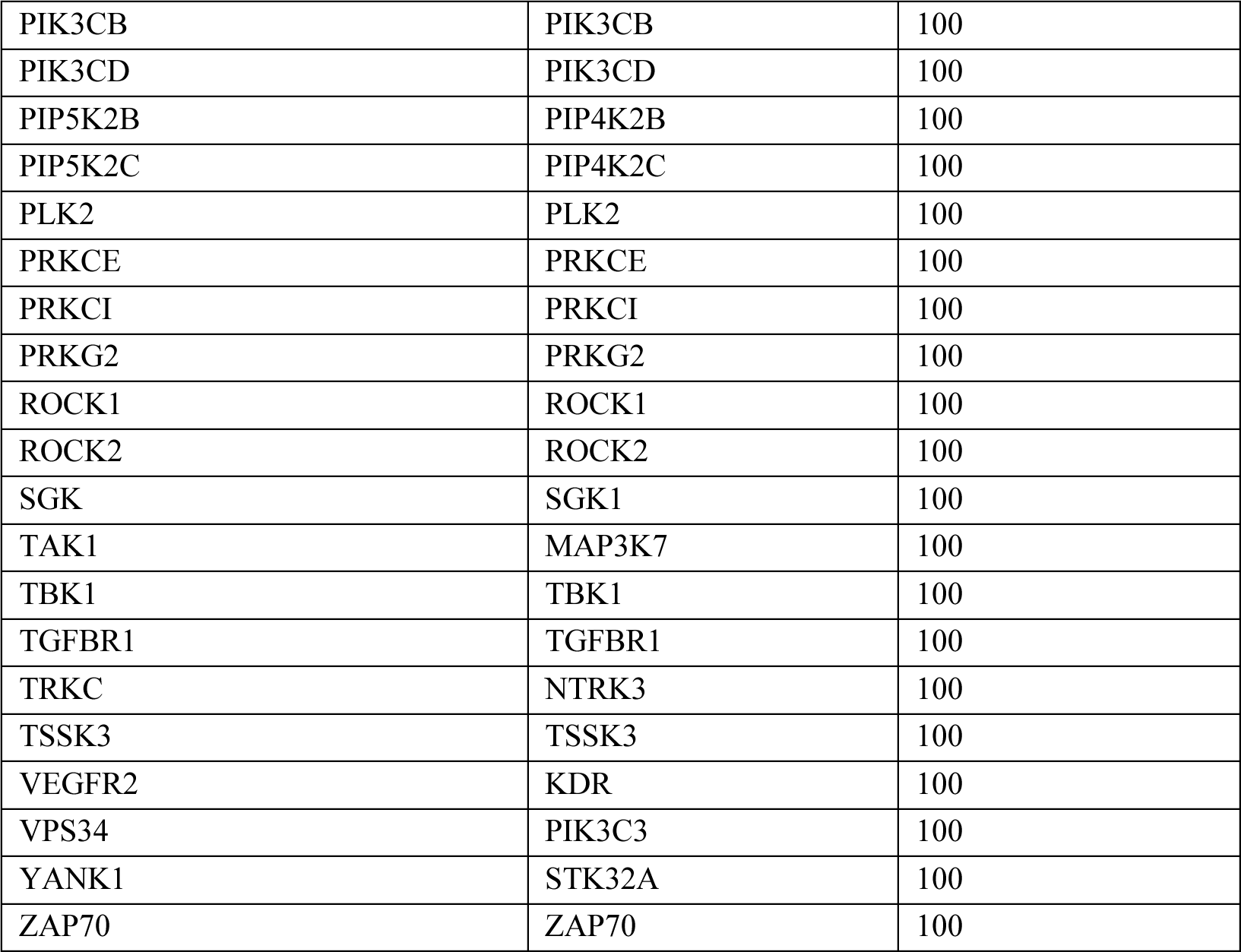
KINOMEscan Profiling of BJG-05-039 @ 10μM.

Table S3. Proteomic data of BJG-05-039 @ 10μM for 5 hours in MOLT4 cells.

(See attachment)

### LEAD CONTACT AND MATERIALS AVAILABILITY

Further information and request for resources and reagents should be directed to and will be fulfilled by the Lead Contact, Jonathan Chernoff. Requested compounds will be provided following completion of an MTA.

#### General Chemistry Methods

Analytical grade solvents and commercially available reagents were purchased from commercial sources and used directly without further purification unless otherwise stated. Thin-layer chromatography (TLC) was carried out on Merck 60 F254 precoated, glass silica plates which were visualized by ultraviolet light. Experiments were conducted under ambient conditions unless otherwise stated. ^1^H-NMR, ^13^C-NMR, and ^19^F-NMR spectra were recorded at room temperature using a Bruker 500 (^1^H-NMR at 500 MHz, ^13^C-NMR at 125 MHz, and ^19^F-NMR at 471 MHz). Chemical shifts are reported in ppm with reference to solvent signals [^1^H-NMR: CDCl3 (7.26 ppm), DMSO-*d*6 (2.50 ppm); ^13^C-NMR: CDCl3 (77.16 ppm), DMSO-*d*6 (39.52 ppm)]. Signal patterns are indicated as s, singlet; br s, broad singlet; d, doublet; t, triplet, q, quartet; p, pentet; and m, multiplet. Mass spectrometry (MS) analysis was obtained on a Waters Acquity UPLC-MS system using electrospray ionization (ESI) in positive ion mode, reporting the molecular ion [M+H]^+^, [M+Na]^+^, or a suitable fragment ion. Flash chromatography purification was conducted using an ISCO CombiFlash RF+ with RediSep Rf silica cartridges. Preparative reverse-phase HPLC purification was conducted using a Waters model 2545 pump and 2489 UV/Vis detector using SunFire Prep C18 5 μm columns (18x100 mm, 20 mL/min flow rate; 30x250 mm, 40 mL/min flow rate), and a gradient solvent system of water (0.035% TFA)/methanol (0.035% TFA) or water (0.035% TFA)/acetonitrile (0.035% TFA).

### Experimental Procedures and Characterizations

#### General Synthetic Scheme

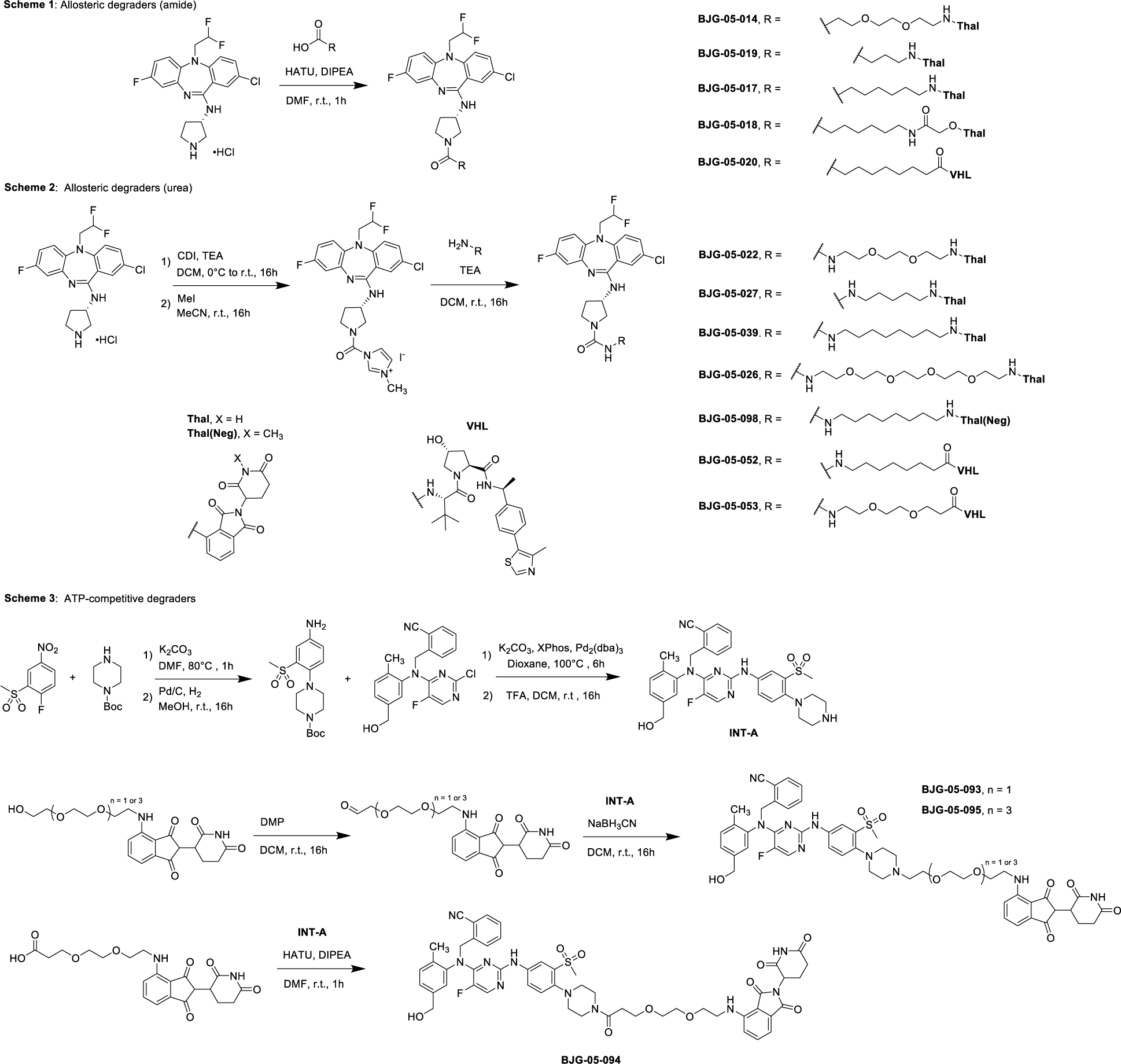

**Synthetic Scheme 1**: Allosteric degraders (amide exit vector)

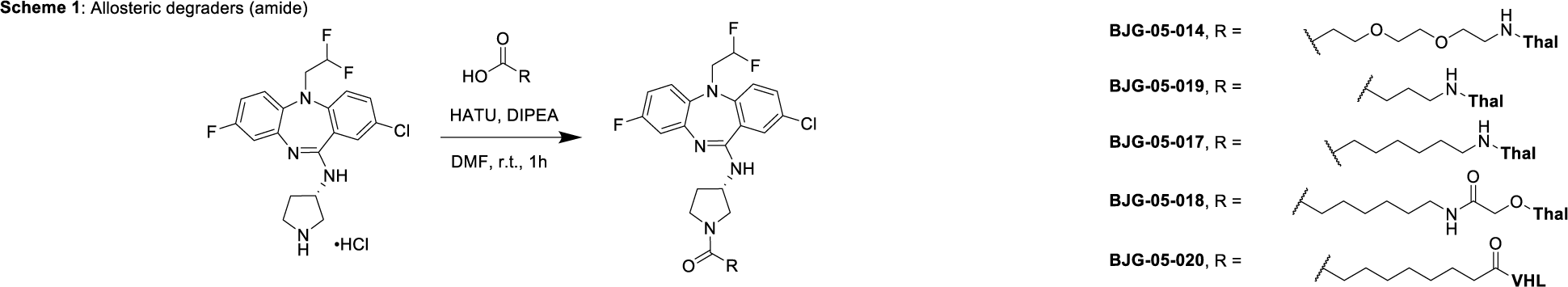

(S)-2-chloro-5-(2,2-difluoroethyl)-8-fluoro-N-(pyrrolidin-3-yl)-5H-dibenzo[b,e][1,4]diazepin- 11-amine (HCl salt)

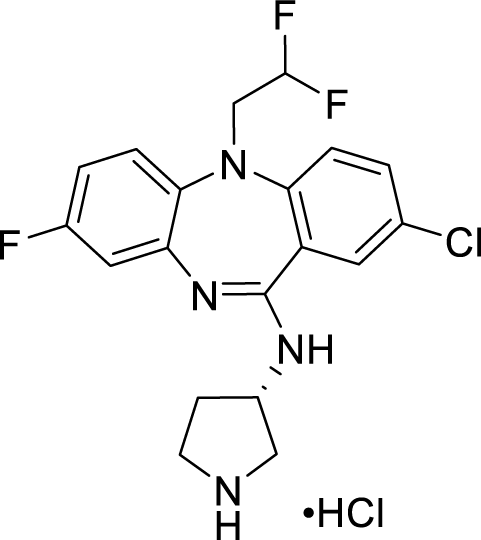

The compound was prepared according to the reported procedure.^1^

4-((2-(2-(3-((S)-3-((2-chloro-5-(2,2-difluoroethyl)-8-fluoro-5H-dibenzo[b,e][1,4]diazepin-11- yl)amino)pyrrolidin-1-yl)-3-oxopropoxy)ethoxy)ethyl)amino)-2-(2,6-dioxopiperidin-3- yl)isoindoline-1,3-dione **BJG-05-014**

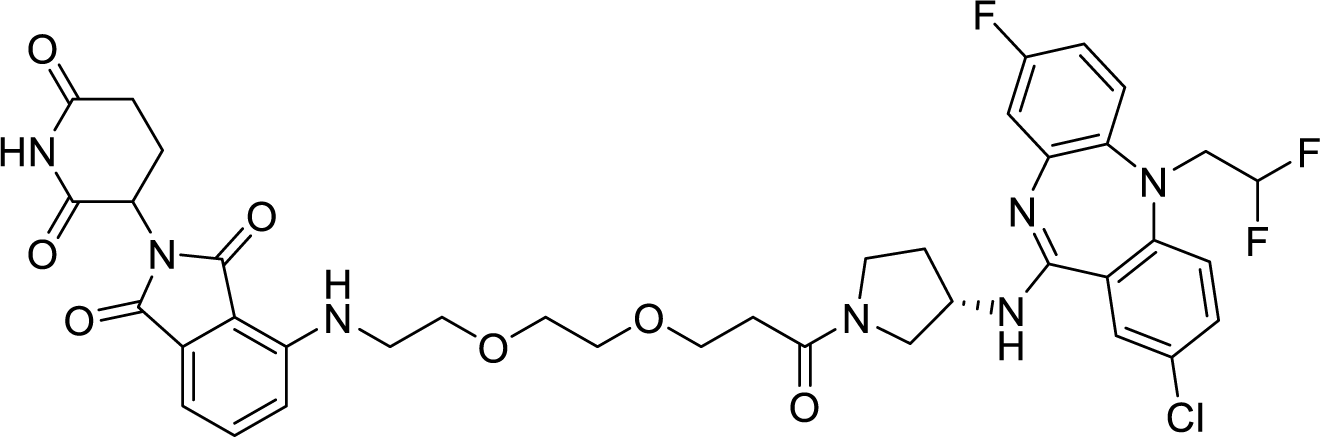

To a solution of solution of 3-(2-(2-((2-(2,6-dioxopiperidin-3-yl)-1,3-dioxoisoindolin-4- yl)amino)ethoxy)ethoxy)propanoic acid (13.0 mg, 0.030 mmol, 1 equv) in N, N-dimethylformamide (0.6 mL), disopropylamine (20.9 uL, 0.120 mmol, 4 equiv) and HATU (11.4 mg, 0.030 mmol, 1 equiv) were added. After stirring the mixture at room temperature for 5 minutes, (S)-2-chloro-5-(2,2-difluoroethyl)-8-fluoro-N-(pyrrolidin-3-yl)-5H- dibenzo[b,e][1,4]diazepin-11-amine (HCl salt) (12.9 mg, 0.030 mmol, 1 equiv) was added. After 2 hours of stirring, the reaction mixture was diluted with 1.0 mL of N, N-dimethylformamide and purified by reverse-phase prep HPLC (95-15% H2O/MeOH, 40 mL/min, 45 min). Lyophilization from H2O/MeCN provided the title compound as a yellow powder (3.8 mg, 16% yield TFA salt).

^1^H NMR (500 MHz, DMSO-*d*6) δ 11.09 (s, 1H), 7.79 – 7.39 (m, 4H), 7.32 (s, 1H), 7.14 (ddd, *J* = 9.6, 6.2, 3.8 Hz, 1H), 7.07 – 6.78 (m, 3H), 6.60 (t, *J* = 6.3 Hz, 1H), 6.03 (t, *J* = 55.8 Hz, 1H), 5.05 (dt, *J* = 12.8, 4.7 Hz, 1H), 4.71 – 4.52 (m, 1H), 4.23 (dd, *J* = 49.1, 14.0 Hz, 2H), 3.89 – 3.51 (m, 14H), 2.88 (td, *J* = 15.5, 14.2, 4.6 Hz, 1H), 2.63 – 2.52 (m, 2H), 2.35 – 2.09 (m, 1H), 2.07 –1.94 (m, 2H). (Signals broadened due to rotational isomerism). LRMS (ESI) calculated for [M+H]^+^ 810.26, found 809.71.

4-((7-((S)-3-((2-chloro-5-(2,2-difluoroethyl)-8-fluoro-5H-dibenzo[b,e][1,4]diazepin-11- yl)amino)pyrrolidin-1-yl)-7-oxoheptyl)amino)-2-(2,6-dioxopiperidin-3-yl)isoindoline-1,3-dione **BJG-05-017**

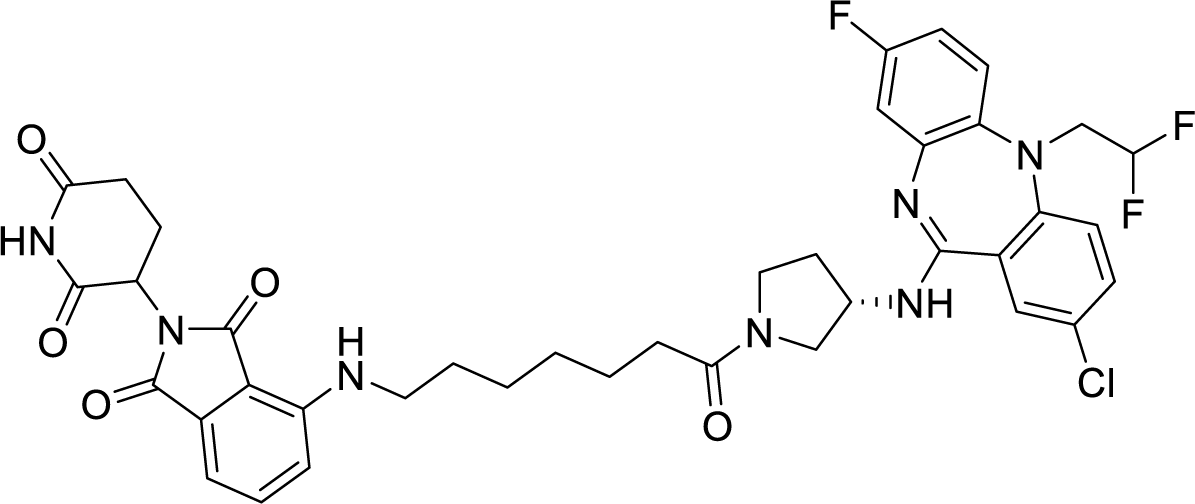

To a solution of solution of 7-((2-(2,6-dioxopiperidin-3-yl)-1,3-dioxoisoindolin-4- yl)amino)heptanoic acid (12.0 mg, 0.030 mmol, 1 equv) in N, N-dimethylformamide (0.6 mL), disopropylamine (20.9 uL, 0.120 mmol, 4 equiv) and HATU (11.4 mg, 0.030 mmol, 1 equiv) were added. After stirring the mixture at room temperature for 5 minutes, (S)-2-chloro-5-(2,2- difluoroethyl)-8-fluoro-N-(pyrrolidin-3-yl)-5H-dibenzo[b,e][1,4]diazepin-11-amine (HCl salt) (12.9 mg, 0.030 mmol, 1 equiv) was added. After 2 hours of stirring, the reaction mixture was diluted with 1.0 mL of N, N-dimethylformamide and purified by reverse-phase prep HPLC (95- 15% H2O/MeOH, 40 mL/min, 45 min). Lyophilization from H2O/MeCN provided the title compound as a yellow powder (2.1 mg, 9% yield TFA salt).

^1^H NMR (500 MHz, DMSO-*d*6) δ 11.08 (s, 1H), 7.62 – 7.44 (m, 3H), 7.43 – 7.26 (m, 2H), 7.17 –7.03 (m, 2H), 7.01 (d, *J* = 7.0 Hz, 1H), 6.77 – 6.61 (m, 2H), 6.52 (p, *J* = 6.0 Hz, 1H), 5.97 (tt, *J* = 55.7, 3.8 Hz, 1H), 5.11 – 4.98 (m, 1H), 4.63 – 4.46 (m, 1H), 4.25 – 3.98 (m, 2H), 3.75 – 3.59 (m, 1H), 3.58 – 3.34 (m, 5H), 2.88 (ddd, *J* = 17.4, 13.6, 5.4 Hz, 1H), 2.64 – 2.52 (m, 2H), 2.31 – 1.92 (m, 5H), 1.64 – 1.42 (m, 5H), 1.41 – 1.16 (m, 7H). 6:1 mix of rotamers. LRMS (ESI) calculated for [M+H]^+^ 778.26, found 777.71. N-(7-((S)-3-((2-chloro-5-(2,2-difluoroethyl)-8-fluoro-5H-dibenzo[b,e][1,4]diazepin-11- yl)amino)pyrrolidin-1-yl)-7-oxoheptyl)-2-((2-(2,6-dioxopiperidin-3-yl)-1,3-dioxoisoindolin-4- yl)oxy)acetamide **BJG-05-018**

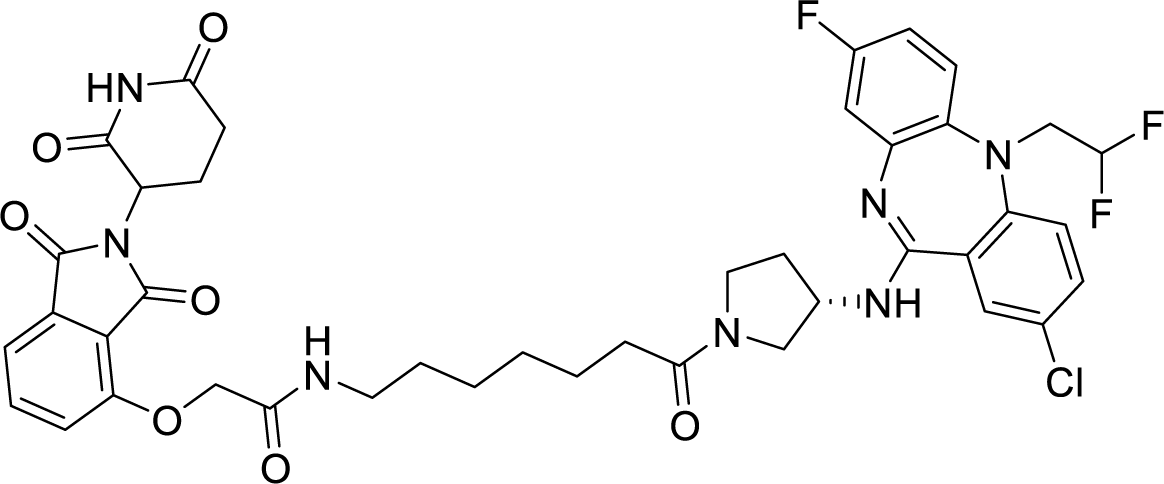

To a solution of solution of 7-(2-((2-(2,6-dioxopiperidin-3-yl)-1,3-dioxoisoindolin-4- yl)oxy)acetamido)heptanoic acid (13.8 mg, 0.030 mmol, 1 equv) in N, N-dimethylformamide (0.6 mL), disopropylamine (20.9 uL, 0.120 mmol, 4 equiv) and HATU (11.4 mg, 0.030 mmol, 1 equiv) were added. After stirring the mixture at room temperature for 5 minutes, (S)-2-chloro-5- (2,2-difluoroethyl)-8-fluoro-N-(pyrrolidin-3-yl)-5H-dibenzo[b,e][1,4]diazepin-11-amine (HCl salt) (12.9 mg, 0.030 mmol, 1 equiv) was added. After 2 hours of stirring, the reaction mixture was diluted with 1.0 mL of N, N-dimethylformamide and purified by reverse-phase prep HPLC (95-15% H2O/MeOH, 40 mL/min, 45 min). Lyophilization from H2O/MeCN provided the title compound as a yellow powder (2.5 mg, 10% yield TFA salt).

^1^H NMR (500 MHz, DMSO-*d*6) δ 11.11 (s, 1H), 7.97 – 7.88 (m, 1H), 7.85 – 7.78 (m, 1H), 7.61 – 7.46 (m, 3H), 7.42 – 7.28 (m, 3H), 7.11 (t, *J* = 7.7 Hz, 1H), 6.77 – 6.62 (m, 2H), 5.99 (tt, *J* = 55.1, 3.8 Hz, 1H), 5.12 (ddd, *J* = 12.6, 5.4, 2.2 Hz, 1H), 4.77 (s, 2H), 4.65 – 4.46 (m, 1H), 4.26 – 3.97 (m, 2H), 3.75 – 3.59 (m, 1H), 3.57 – 3.35 (m, 5H), 3.12 (dq, *J* = 13.2, 6.3 Hz, 2H), 2.90 (ddd, *J* = 19.0, 13.7, 5.4 Hz, 1H), 2.64 – 2.52 (m, 2H), 2.30 – 1.94 (m, 5H), 1.56 – 1.35 (m, 4H), 1.34 – 1.21 (m, 5H). LRMS (ESI) calculated for [M+H]^+^ 836.27, found 835.61. 4-((4-((S)-3-((2-chloro-5-(2,2-difluoroethyl)-8-fluoro-5H-dibenzo[b,e][1,4]diazepin-11- yl)amino)pyrrolidin-1-yl)-4-oxobutyl)amino)-2-(2,6-dioxopiperidin-3-yl)isoindoline-1,3-dione **BJG-05-019**

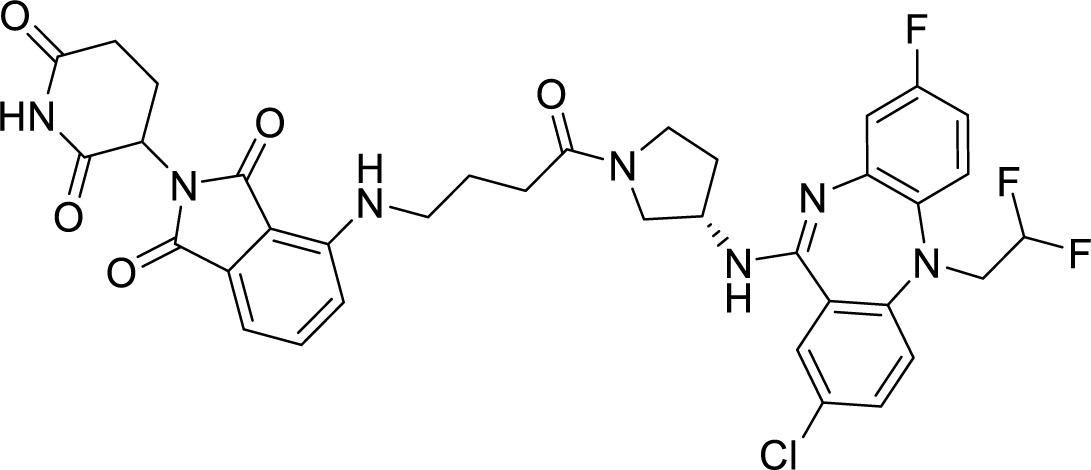

To a solution of solution of 4-((2-(2,6-dioxopiperidin-3-yl)-1,3-dioxoisoindolin-4- yl)amino)butanoic acid (14.4 mg, 0.040 mmol, 1 equv) in N, N-dimethylformamide (0.8 mL), disopropylamine (27.9 uL, 0.160 mmol, 4 equiv) and HATU (15.2 mg, 0.040 mmol, 1 equiv) were added. After stirring the mixture at room temperature for 5 minutes, (S)-2-chloro-5-(2,2- difluoroethyl)-8-fluoro-N-(pyrrolidin-3-yl)-5H-dibenzo[b,e][1,4]diazepin-11-amine (HCl salt) (17.2 mg, 0.040 mmol, 1 equiv) was added. After 2 hours of stirring, the reaction mixture was diluted with 1.0 mL of N, N-dimethylformamide and purified by reverse-phase prep HPLC (95- 15% H2O/MeOH, 40 mL/min, 45 min). Lyophilization from H2O/MeCN provided the title compound as a yellow powder (4.1 mg, 14% yield TFA salt).

^1^H NMR (500 MHz, DMSO-*d*6) δ 11.09 (s, 1H), 7.80 – 7.29 (m, 5H), 7.23 – 7.15 (m, 1H), 7.14 – 6.97 (m, 2H), 6.68 (s, 1H), 6.09 (t, *J* = 55.0 Hz, 1H), 5.04 (dt, *J* = 12.9, 5.3 Hz, 1H), 4.65 (d, *J* = 26.9 Hz, 1H), 4.41 – 4.12 (m, 2H), 3.41 – 3.28 (m, 2H), 2.96 – 2.78 (m, 1H), 2.64 – 2.52 (m, 2H), 2.44 – 1.96 (m, 5H), 1.90 – 1.74 (m, 2H). (Signals broadened due to rotational isomerism). LRMS (ESI) calculated for [M+H]^+^ 736.21, found 735.71.

(2S,4R)-1-((S)-2-(9-((S)-3-((2-chloro-5-(2,2-difluoroethyl)-8-fluoro-5H- dibenzo[b,e][1,4]diazepin-11-yl)amino)pyrrolidin-1-yl)-9-oxononanamido)-3,3- dimethylbutanoyl)-4-hydroxy-N-((S)-1-(4-(4-methylthiazol-5-yl)phenyl)ethyl)pyrrolidine-2- carboxamide **BJG-05-020**

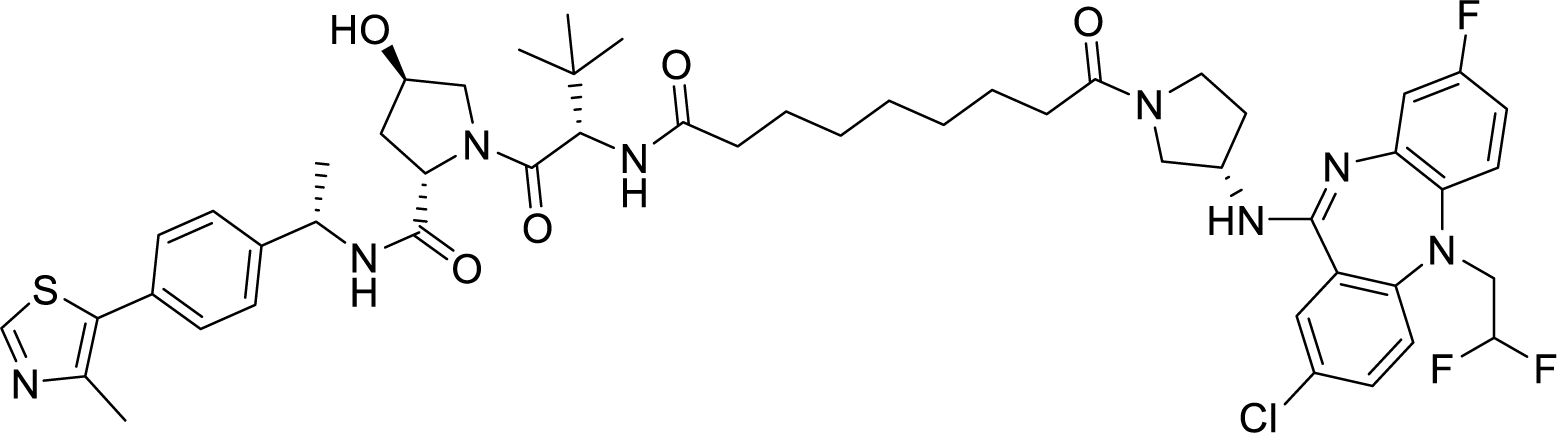

To a solution of solution of 9-(((S)-1-((2S,4R)-4-hydroxy-2-(((S)-1-(4-(4-methylthiazol-5- yl)phenyl)ethyl)carbamoyl)pyrrolidin-1-yl)-3,3-dimethyl-1-oxobutan-2-yl)amino)-9- oxononanoic acid (9.2 mg, 0.015 mmol, 1 equv) in N, N-dimethylformamide (0.3 mL), disopropylamine (10.5 uL, 0.060 mmol, 4 equiv) and HATU (5.7 mg, 0.015 mmol, 1 equiv) were added. After stirring the mixture at room temperature for 5 minutes, (S)-2-chloro-5-(2,2- difluoroethyl)-8-fluoro-N-(pyrrolidin-3-yl)-5H-dibenzo[b,e][1,4]diazepin-11-amine (HCl salt) (6.5 mg, 0.015 mmol, 1 equiv) was added. After 2 hours of stirring, the reaction mixture was diluted with 1.0 mL of N, N-dimethylformamide and purified by reverse-phase prep HPLC (95- 5% H2O/MeOH, 40 mL/min, 45 min). Lyophilization from H2O/MeCN provided the title compound as a white powder (2.7 mg, 18% yield TFA salt).

^1^H NMR (500 MHz, DMSO-*d*6) δ 8.98 (s, 1H), 8.37 (d, *J* = 8.0 Hz, 1H), 7.77 (d, *J* = 9.1 Hz, 1H), 7.74 – 7.46 (m, 3H), 7.46 – 7.30 (m, 5H), 7.26 – 6.91 (m, 1H), 6.10 (t, *J* = 55.3 Hz, 1H), 4.91 (p, *J* = 7.0 Hz, 1H), 4.73 – 4.58 (m, 1H), 4.51 (dd, *J* = 9.3, 5.2 Hz, 1H), 4.42 (t, *J* = 8.1 Hz, 1H), 3.63 – 3.55 (m, 4H), 2.45 (s, 4H), 2.31 – 1.94 (m, 7H), 1.83 – 1.75 (m, 1H), 1.56 – 1.40 (m, 5H), 1.37 (d, *J* = 6.9 Hz, 3H), 1.32 – 1.16 (m, 7H), 0.93 (s, 9H). LRMS (ESI) calculated for [M+H]^+^ 991.42, found 990.73.

**Synthetic Scheme 2**: Allosteric degraders (urea exit vector)

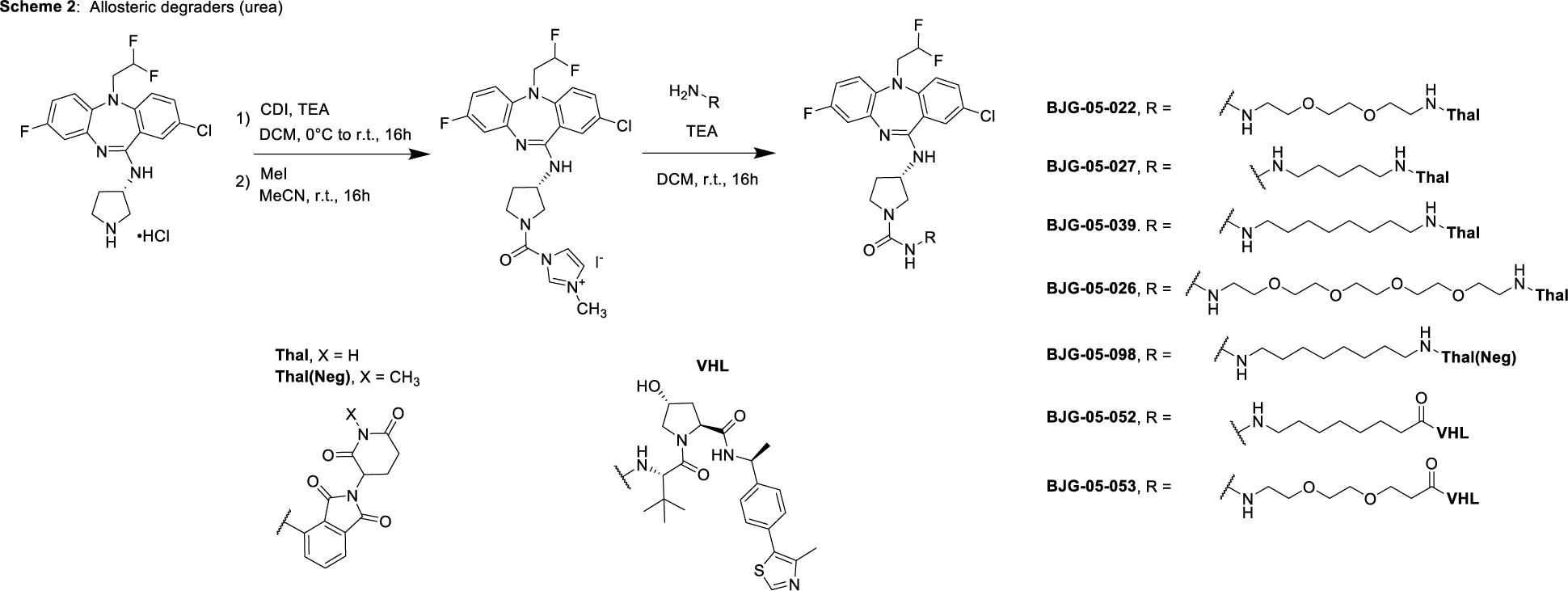

(S)-1-(3-((2-chloro-5-(2,2-difluoroethyl)-8-fluoro-5H-dibenzo[b,e][1,4]diazepin-11- yl)amino)pyrrolidine-1-carbonyl)-3-methyl-1H-imidazol-3-ium iodide

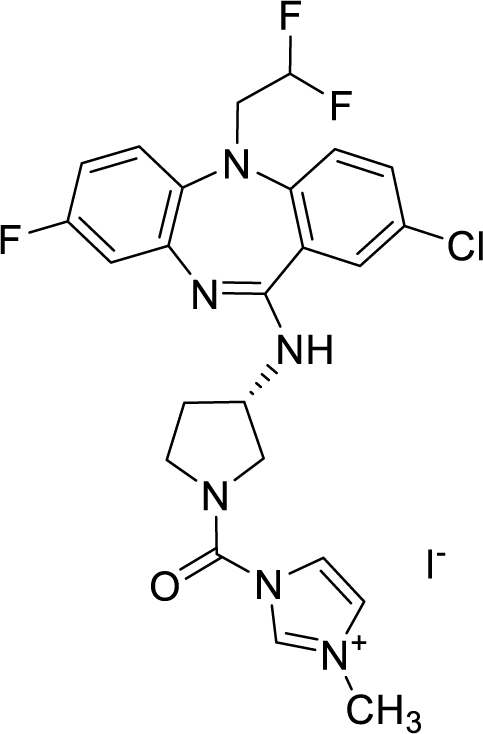

A solution of carbonyldiimidazole (50 mg, 0.31 mmol, 1.1 equiv) in dichloromethane (1.4 mL) was cooled to 0 °C at which (S)-2-chloro-5-(2,2-difluoroethyl)-8-fluoro-N-(pyrrolidin-3-yl)-5H- dibenzo[b,e][1,4]diazepin-11-amine (HCl salt) (120.8 mg, 0.28 mmol, 1.0 equiv) was added followed by addition of triethylamine (39 uL, 0.28 mmol, 1 equiv). The mixture was then removed from ice and allowed to stir at room temperature overnight. The reaction was diluted with 7 mL of water and organic layer was then extracted with dichloromethane (5x2 mL). The combined organic layers were washed with brine, dried over MgSO4, filtered, and concentrated to provide a clear yellow oil (60 mg). Product was shown as the major component via mass spec: LRMS (ESI) calculated for [M+H]^+^ 489.13, found 488.90. The crude product was moved forward without purification.

To a solution of 50 mg of crude residue in acetonitrile (1.4 mL), iodomethane (70 uL, 1.12 mmol, 4 equiv) was added at room temperature and stirred overnight. The reaction mixture was concentrated to provide the titled compound as a yellow oil (62.5 mg, 99% yield). The product was shown as the major component via mass spec, LRMS (ESI) calculated for [M+H]^+^ 504.16, found 502.78, and moved on to the next step without purification. (3S)-3-((2-chloro-5-(2,2-difluoroethyl)-8-fluoro-5H-dibenzo[b,e][1,4]diazepin-11-yl)amino)-N- (2-(2-(2-((2-(2,6-dioxopiperidin-3-yl)-1,3-dioxoisoindolin-4- yl)amino)ethoxy)ethoxy)ethyl)pyrrolidine-1-carboxamide **BJG-05-022**

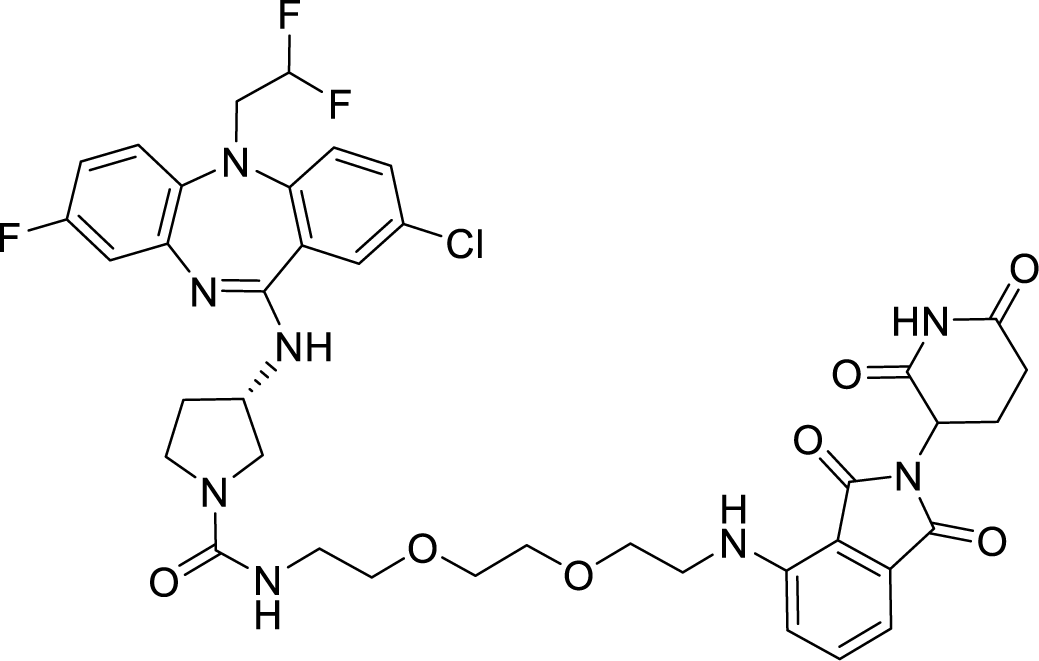

To a solution of solution of 4-((2-(2-(2-aminoethoxy)ethoxy)ethyl)amino)-2-(2,6-dioxopiperidin- 3-yl)isoindoline-1,3-dione (15.6 mg, 0.030 mmol, 1 equiv) and (S)-1-(3-((2-chloro-5-(2,2- difluoroethyl)-8-fluoro-5H-dibenzo[b,e][1,4]diazepin-11-yl)amino)pyrrolidine-1-carbonyl)-3- methyl-1H-imidazol-3-ium iodide (18.9 mg, 0.030 mmol, 1 equv) in dichloromethane (0.3 mL), triethylamine (10.5 uL, 0.075 mmol, 2.5 equiv) was added. After 16 hours of stirring, the reaction mixture was rotovapped and diluted with 1.0 mL of N, N-dimethylformamide and purified by reverse-phase prep HPLC (95-15% H2O/MeOH, 40 mL/min, 45 min). Lyophilization from H2O/MeCN provided the title compound as a yellow powder (5.1 mg, 21% yield TFA salt).

^1^H NMR (500 MHz, DMSO-*d*6) δ 11.09 (s, 1H), 7.69 (s, 1H), 7.58 (ddd, *J* = 8.5, 7.0, 1.5 Hz, 2H), 7.49 (s, 1H), 7.39 (s, 1H), 7.27 – 6.98 (m, 4H), 6.60 (s, 1H), 6.33 – 5.89 (m, 2H), 5.06 (dd, *J* = 12.6, 5.4 Hz, 1H), 4.60 (s, 1H), 4.40 – 4.14 (m, 2H), 4.10 (d, *J* = 1.1 Hz, 1H), 3.57 – 3.29 (m, 12H), 3.18 (dq, *J* = 12.1, 6.1 Hz, 2H), 2.88 (ddd, *J* = 16.7, 13.6, 5.4 Hz, 1H), 2.62 – 2.52 (m, 2H), 2.34 – 1.95 (m, 3H). LRMS (ESI) calculated for [M+H]^+^ 825.26, found 824.71. (3S)-3-((2-chloro-5-(2,2-difluoroethyl)-8-fluoro-5H-dibenzo[b,e][1,4]diazepin-11-yl)amino)-N- (14-((2-(2,6-dioxopiperidin-3-yl)-1,3-dioxoisoindolin-4-yl)amino)-3,6,9,12- tetraoxatetradecyl)pyrrolidine-1-carboxamide **BJG-05-026**

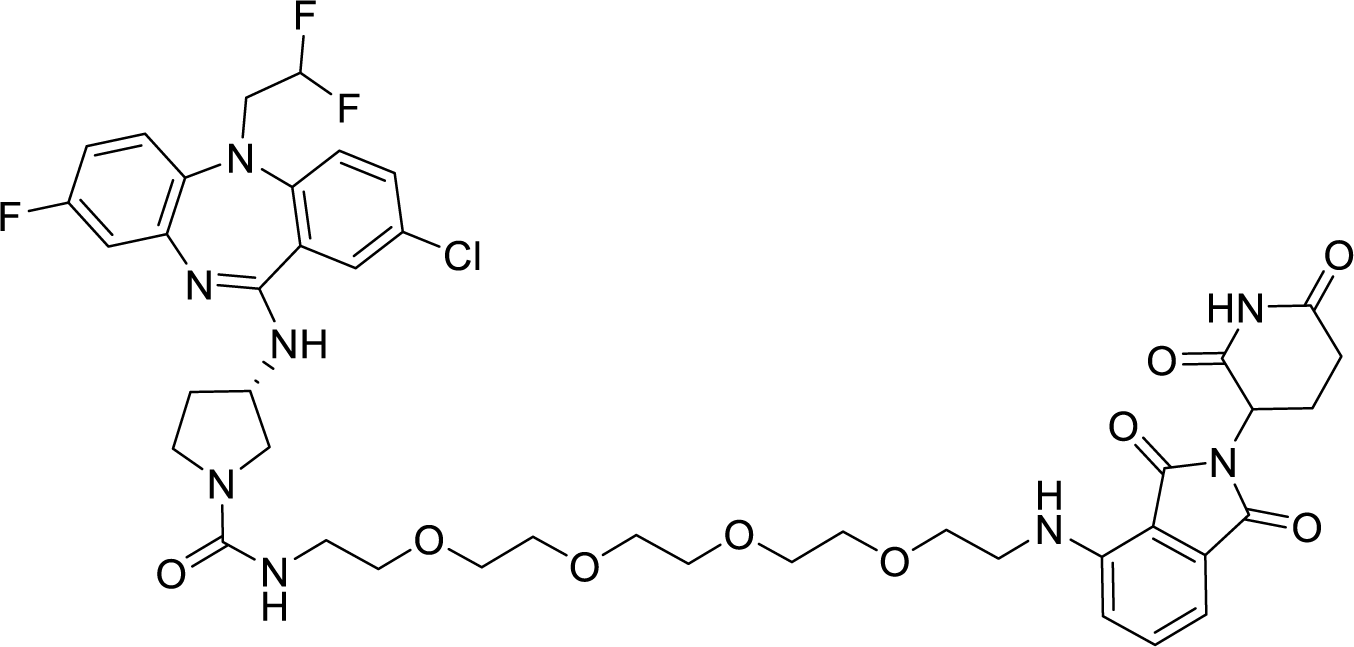

To a solution of solution of 4-((14-amino-3,6,9,12-tetraoxatetradecyl)amino)-2-(2,6- dioxopiperidin-3-yl)isoindoline-1,3-dione (23 mg, 0.030 mmol, 1 equiv) and (S)-1-(3-((2-chloro- 5-(2,2-difluoroethyl)-8-fluoro-5H-dibenzo[b,e][1,4]diazepin-11-yl)amino)pyrrolidine-1- carbonyl)-3-methyl-1H-imidazol-3-ium iodide (18.9 mg, 0.030 mmol, 1 equv) in dichloromethane (0.3 mL), triethylamine (10.5 uL, 0.075 mmol, 2.5 equiv) was added. After 16 hours of stirring, the reaction mixture was rotovapped and diluted with 1.0 mL of N, N-dimethylformamide and purified by reverse-phase prep HPLC (95-15% H2O/MeOH, 40 mL/min, 45 min). Lyophilization from H2O/MeCN provided the title compound as a yellow powder (4.8 mg, 18% yield TFA salt).

^1^H NMR (500 MHz, DMSO-*d*6) δ 11.09 (s, 1H), 7.67 – 7.46 (m, 3H), 7.44 – 7.24 (m, 2H), 7.18 – 7.07 (m, 2H), 7.04 (d, *J* = 7.0 Hz, 1H), 6.79 – 6.63 (m, 2 H), 6.60 (t, *J* = 5.9 Hz, 1H), 6.21 – 5.82 (m, 2H), 5.05 (dd, *J* = 12.7, 5.5 Hz, 1H), 4.53 (t, *J* = 4.9 Hz, 1H), 4.28 – 3.93 (m, 2H), 3.64 – 3.34 (m, 20H), 3.24 – 3.10 (m, 2H), 2.88 (ddd, *J* = 16.7, 13.7, 5.4 Hz, 1H), 2.62 – 2.51 (m, 2H), 2.24 – 1.87 (m, 3H). LRMS (ESI) calculated for [M+H]^+^ 913.31, found 912.62. (3S)-3-((2-chloro-5-(2,2-difluoroethyl)-8-fluoro-5H-dibenzo[b,e][1,4]diazepin-11-yl)amino)-N- (5-((2-(2,6-dioxopiperidin-3-yl)-1,3-dioxoisoindolin-4-yl)amino)pentyl)pyrrolidine-1- carboxamide **BJG-05-027**

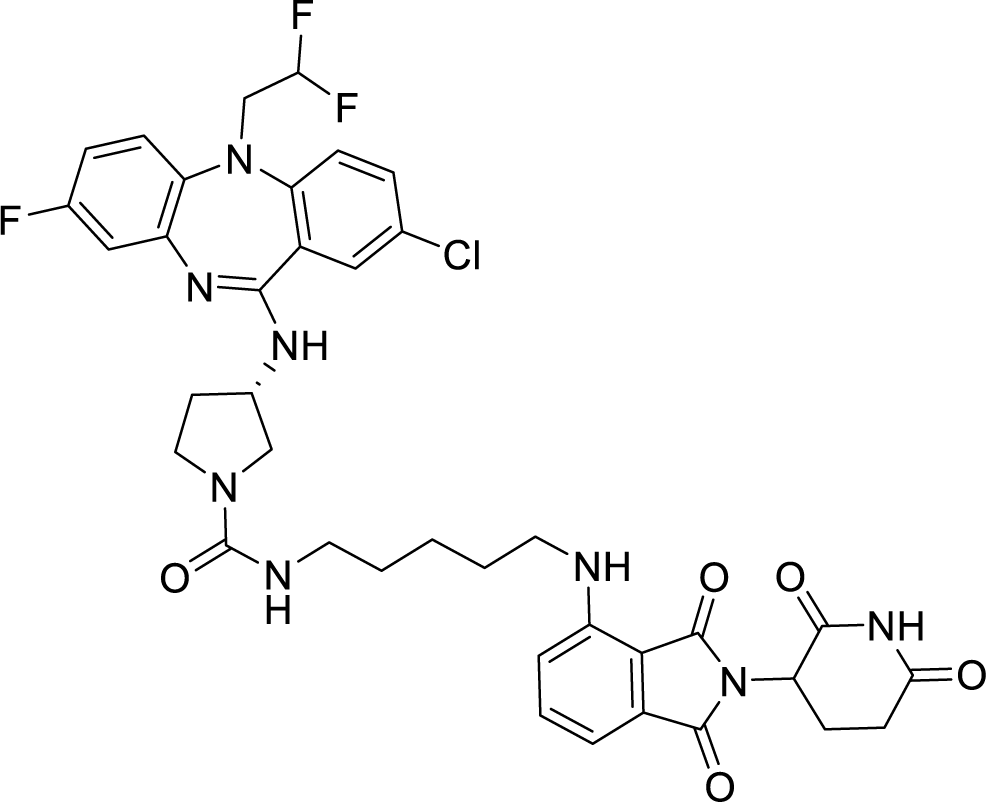

To a solution of solution of 4-((5-aminopentyl)amino)-2-(2,6-dioxopiperidin-3-yl)isoindoline- 1,3-dione (15.1 mg, 0.030 mmol, 1 equiv) and (S)-1-(3-((2-chloro-5-(2,2-difluoroethyl)-8-fluoro- 5H-dibenzo[b,e][1,4]diazepin-11-yl)amino)pyrrolidine-1-carbonyl)-3-methyl-1H-imidazol-3-ium iodide (18.9 mg, 0.030 mmol, 1 equv) in dichloromethane (0.3 mL), triethylamine (10.5 uL, 0.075 mmol, 2.5 equiv) was added. After 16 hours of stirring, the reaction mixture was rotovapped and diluted with 1.0 mL of N, N-dimethylformamide and purified by reverse-phase prep HPLC (95-15% H2O/MeOH, 40 mL/min, 45 min). Lyophilization from H2O/MeCN provided the title compound as a yellow powder (3.8 mg, 16% yield TFA salt).

^1^H NMR (500 MHz, DMSO-*d*6) δ 11.09 (s, 1H), 7.70 (s, 1H), 7.65 – 7.54 (m, 2H), 7.53 – 7.29 (m, 2H), 7.12 – 6.98 (m, 4H), 6.92 (dq, *J* = 7.5, 0.8 Hz, 1H), 6.57 – 6.41 (m, 2H), 6.30 – 5.92 (m, 2H), 5.05 (ddd, *J* = 12.8, 5.6, 1.3 Hz, 1H), 4.67 (s, 1H), 4.33 (s, 4H), 3.74 – 3.64 (m, 1H), 3.36 (dd, *J* = 8.7, 5.8 Hz, 2H), 3.31 – 3.24 (m, 2H), 3.14 – 2.97 (m, 4H), 2.88 (ddd, *J* = 16.8, 13.8, 5.4 Hz, 1H), 2.62 – 2.52 (m, 2H), 2.02 (ddd, *J* = 13.1, 6.5, 4.2 Hz, 2H), 1.58 (h, *J* = 7.0 Hz, 2H), 1.45 (dq, *J* = 11.3, 7.2 Hz, 2H), 1.34 (dq, *J* = 15.6, 8.0, 7.0 Hz, 2H). LRMS (ESI) calculated for [M+H]^+^ 779.26, found 778.71.

(3S)-3-((2-chloro-5-(2,2-difluoroethyl)-8-fluoro-5H-dibenzo[b,e][1,4]diazepin-11-yl)amino)-N- (8-((2-(2,6-dioxopiperidin-3-yl)-1,3-dioxoisoindolin-4-yl)amino)octyl)pyrrolidine-1- carboxamide **BJG-05-039**

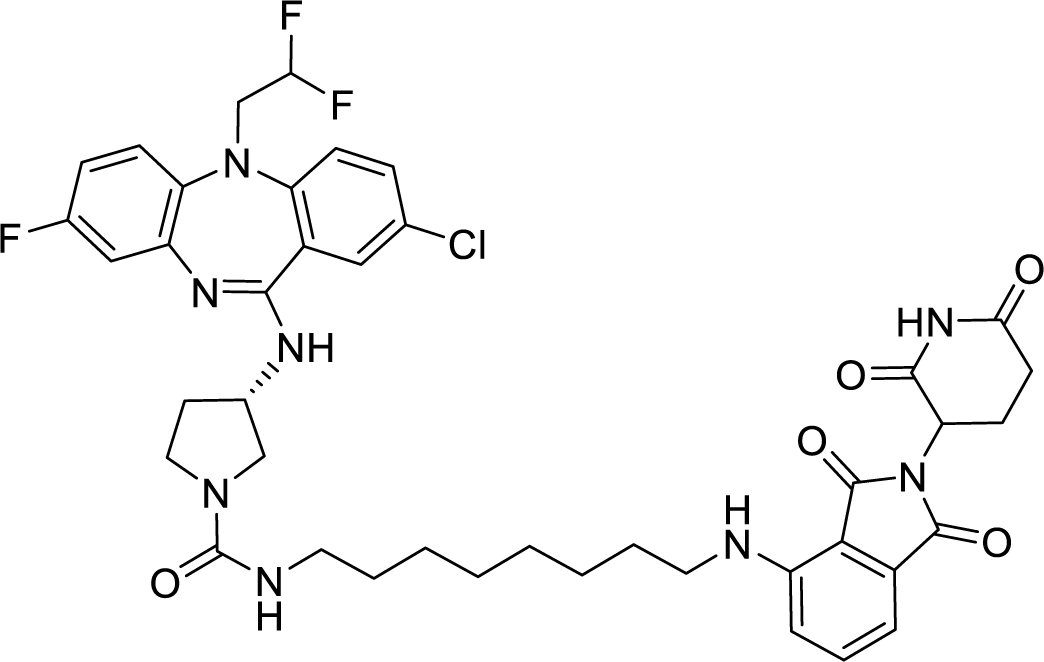

To a solution of solution of 4-((8-aminooctyl)amino)-2-(2,6-dioxopiperidin-3-yl)isoindoline-1,3- dione (20.4 mg, 0.040 mmol, 1 equiv) and (S)-1-(3-((2-chloro-5-(2,2-difluoroethyl)-8-fluoro-5H- dibenzo[b,e][1,4]diazepin-11-yl)amino)pyrrolidine-1-carbonyl)-3-methyl-1H-imidazol-3-ium iodide (25.2 mg, 0.040 mmol, 1 equv) in dichloromethane (0.5 mL), triethylamine (13.9 uL, 0.100 mmol, 2.5 equiv) was added. After 16 hours of stirring, the reaction mixture was rotovapped and diluted with 1.0 mL of N, N-dimethylformamide and purified by reverse-phase prep HPLC (95-15% H2O/MeOH, 40 mL/min, 45 min). Lyophilization from H2O/MeCN provided the title compound as a yellow powder (7.8 mg, 24% yield TFA salt).

^1^H NMR (500 MHz, DMSO-*d*6) δ 11.09 (s, 1H), 7.71 (d, *J* = 9.2 Hz, 1H), 7.64 – 7.49 (m, 3H), 7.48 – 7.35 (m, 1H), 7.16 – 7.08 (m, 2H), 7.02 (d, *J* = 7.0 Hz, 1H), 6.51 (s, 1H), 6.25 – 5.96 (m, 2H), 5.05 (dd, *J* = 12.7, 5.4 Hz, 1H), 4.68 (s, 1H), 4.44 – 4.12 (m, 2H), 3.68 (ddd, *J* = 14.0, 11.1, 6.1 Hz, 1H), 3.39 – 3.31 (m, 2H), 3.28 (t, *J* = 7.5 Hz, 3H), 3.05 – 2.95 (m, 2H), 2.88 (ddd, *J* = 16.8, 13.7, 5.4 Hz, 1H), 2.64 – 2.52 (m, 2H), 2.39 – 2.15 (m, 2H), 2.09 – 1.96 (m, 2H), 1.61 – 1.51 (m, 2H), 1.45 – 1.21 (m, 10H). LRMS (ESI) calculated for [M+H]^+^ 821.21, found 821.15. (S)-3-((2-chloro-5-(2,2-difluoroethyl)-8-fluoro-5H-dibenzo[b,e][1,4]diazepin-11-yl)amino)-N-(8-(((S)-1-((2S,4R)-4-hydroxy-2-(((S)-1-(4-(4-methylthiazol-5- yl)phenyl)ethyl)carbamoyl)pyrrolidin-1-yl)-3,3-dimethyl-1-oxobutan-2-yl)amino)-8- oxooctyl)pyrrolidine-1-carboxamide **BJG-05-052**

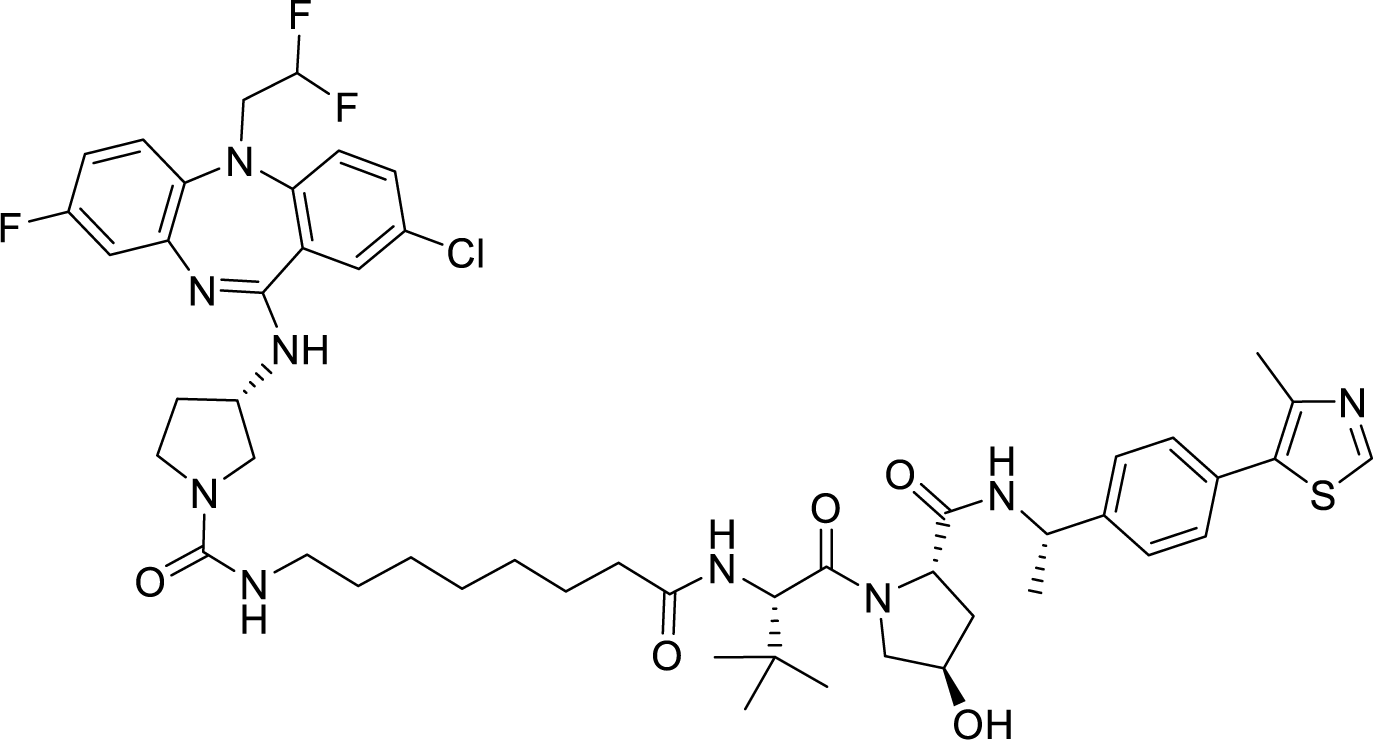

To a solution of solution of (2S,4R)-1-((S)-2-(8-aminooctanamido)-3,3-dimethylbutanoyl)-4- hydroxy-N-((S)-1-(4-(4-methylthiazol-5-yl)phenyl)ethyl)pyrrolidine-2-carboxamide (20.9 mg, 0.030 mmol, 1 equiv) and (S)-1-(3-((2-chloro-5-(2,2-difluoroethyl)-8-fluoro-5H- dibenzo[b,e][1,4]diazepin-11-yl)amino)pyrrolidine-1-carbonyl)-3-methyl-1H-imidazol-3-ium iodide (18.9 mg, 0.030 mmol, 1 equv) in dichloromethane (0.3 mL), triethylamine (10.5 uL, 0.075 mmol, 2.5 equiv) was added. After 16 hours of stirring, the reaction mixture was rotovapped and diluted with 1.0 mL of N, N-dimethylformamide and purified by reverse-phase prep HPLC (95-15% H2O/MeOH, 40 mL/min, 45 min). Lyophilization from H2O/MeCN provided the title compound as a white powder (4.3 mg, 14% yield TFA salt).

^1^H NMR (500 MHz, DMSO-*d*6) δ 8.98 (s, 1H), 8.37 (dd, *J* = 7.9, 1.6 Hz, 1H), 7.77 (dd, *J* = 9.3, 1.9 Hz, 1H), 7.66 – 7.50 (m, 2H), 7.47 – 7.31 (m, 6H), 6.77 (s, 1H), 6.23 – 5.81 (m, 2H), 5.09 (s, 1H), 4.91 (p, *J* = 7.1 Hz, 1H), 4.56 (s, 1H), 4.51 (dd, *J* = 9.4, 2.6 Hz, 1H), 4.42 (td, *J* = 8.0, 1.8 Hz, 1H), 4.35 – 3.98 (m, 3H), 3.72 – 3.53 (m, 3H), 3.52 – 3.43 (m, 1H), 2.99 (dtd, *J* = 17.5, 11.4, 10.9, 7.3 Hz, 2H), 2.45 (d, *J* = 0.9 Hz, 3H), 2.29 – 1.94 (m, 5H), 1.79 (ddd, *J* = 12.9, 8.5, 4.7 Hz, 1H), 1.56 – 1.32 (m, 8H), 1.23 (t, *J* = 6.6 Hz, 8H), 0.93 (s, 9H). LRMS (ESI) calculated for [M+H]^+^ 1006.43, found 1005.73. (S)-3-((2-chloro-5-(2,2-difluoroethyl)-8-fluoro-5H-dibenzo[b,e][1,4]diazepin-11-yl)amino)-N- (2-(2-(3-(((S)-1-((2S,4R)-4-hydroxy-2-(((S)-1-(4-(4-methylthiazol-5-yl)phenyl)ethyl)carbamoyl)pyrrolidin-1-yl)-3,3-dimethyl-1-oxobutan-2-yl)amino)-3- oxopropoxy)ethoxy)ethyl)pyrrolidine-1-carboxamide **BJG-05-053**

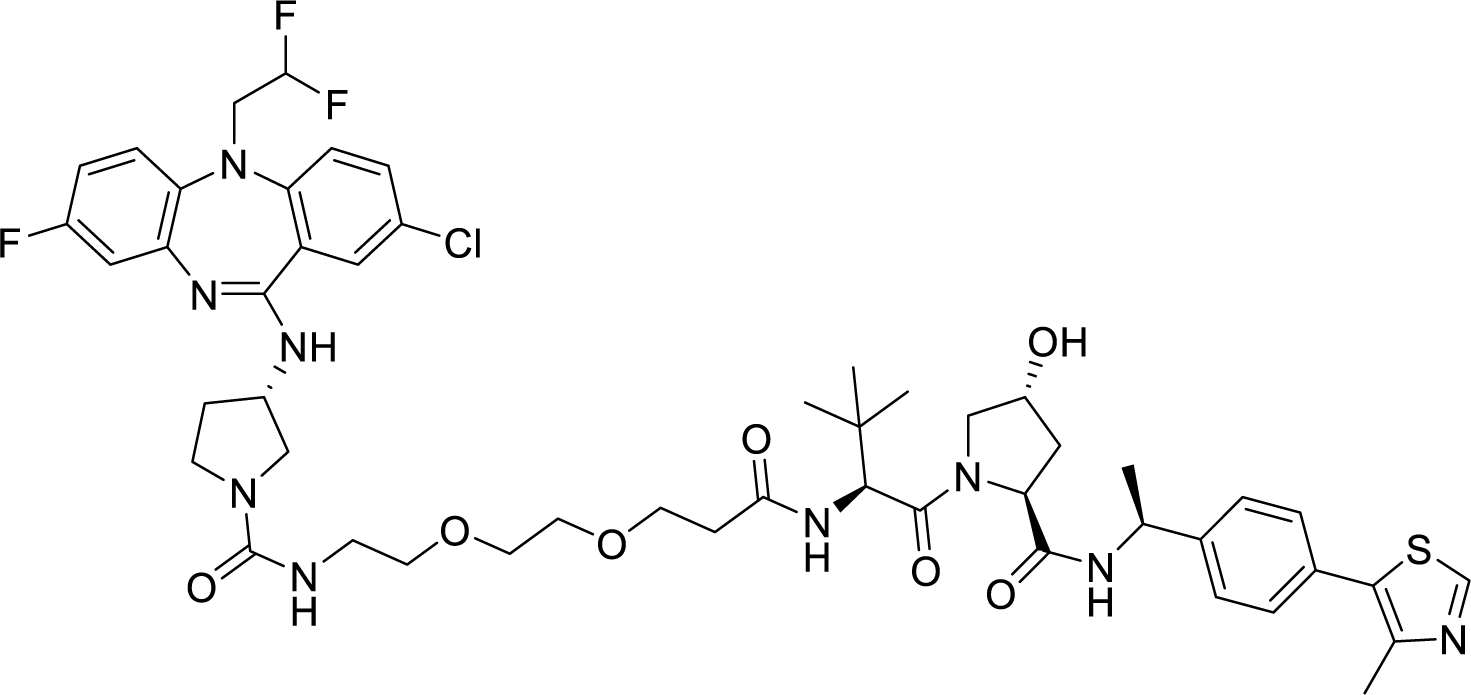

To a solution of solution of (2S,4R)-1-((S)-2-(3-(2-(2-aminoethoxy)ethoxy)propanamido)-3,3- dimethylbutanoyl)-4-hydroxy-N-((S)-1-(4-(4-methylthiazol-5-yl)phenyl)ethyl)pyrrolidine-2- carboxamide (21.5 mg, 0.030 mmol, 1 equiv) and (S)-1-(3-((2-chloro-5-(2,2-difluoroethyl)-8- fluoro-5H-dibenzo[b,e][1,4]diazepin-11-yl)amino)pyrrolidine-1-carbonyl)-3-methyl-1H- imidazol-3-ium iodide (18.9 mg, 0.030 mmol, 1 equv) in dichloromethane (0.3 mL), triethylamine (10.5 uL, 0.075 mmol, 2.5 equiv) was added. After 16 hours of stirring, the reaction mixture was rotovapped and diluted with 1.0 mL of N, N-dimethylformamide and purified by reverse-phase prep HPLC (95-15% H2O/MeOH, 40 mL/min, 45 min). Lyophilization from H2O/MeCN provided the title compound as a white powder (4.9 mg, 16% yield TFA salt).

^1^H NMR (500 MHz, DMSO-*d*6) δ 8.98 (d, *J* = 1.1 Hz, 1H), 8.38 (d, *J* = 7.7 Hz, 1H), 7.86 (dd, *J* = 9.3, 1.9 Hz, 1H), 7.66 (s, 1H), 7.60 – 7.26 (m, 7H), 6.33 – 5.87 (m, 2H), 4.91 (p, *J* = 7.0 Hz, 1H), 4.60 (s, 1H), 4.53 (dd, *J* = 9.3, 2.1 Hz, 1H), 4.42 (t, *J* = 8.1 Hz, 1H), 4.37 – 4.10 (m, 3H), 3.64 – 3.57 (m, 6H), 3.43 – 3.26 (m, 7H), 3.22 – 3.12 (m, 2H), 2.45 (d, *J* = 1.2 Hz, 3H), 2.40 –2.09 (m, 3H), 2.06 – 1.95 (m, 2H), 1.79 (ddd, *J* = 12.9, 8.5, 4.7 Hz, 1H), 1.37 (d, *J* = 7.0 Hz, 3H), 0.93 (s, 9H). (PEG and pyrrolidine methylene protons obscured by water peak). LRMS (ESI) calculated for [M+H]^+^ 1024.41, found 1023.79.

(3S)-3-((2-chloro-5-(2,2-difluoroethyl)-8-fluoro-5H-dibenzo[b,e][1,4]diazepin-11-yl)amino)-N- (8-((2-(1-methyl-2,6-dioxopiperidin-3-yl)-1,3-dioxoisoindolin-4-yl)amino)octyl)pyrrolidine-1- carboxamide **BJG-05-098** (Negative control of BJG-03-039)

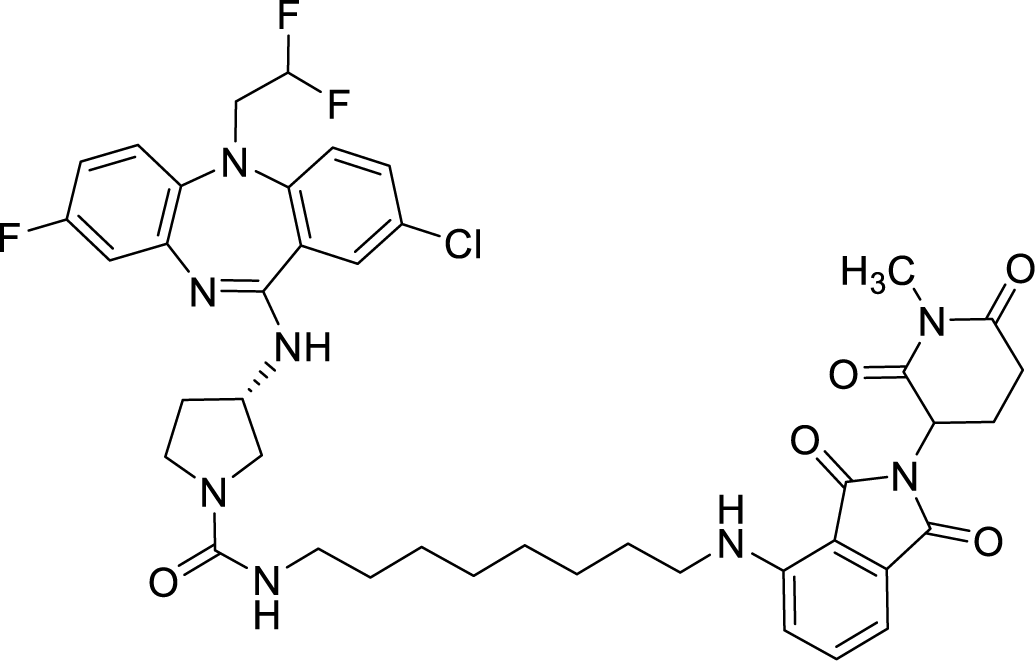

To a solution of solution of 4-((8-aminooctyl)amino)-2-(1-methyl-2,6-dioxopiperidin-3- yl)isoindoline-1,3-dione (17.4 mg, 0.030 mmol, 1 equiv) and (S)-1-(3-((2-chloro-5-(2,2- difluoroethyl)-8-fluoro-5H-dibenzo[b,e][1,4]diazepin-11-yl)amino)pyrrolidine-1-carbonyl)-3- methyl-1H-imidazol-3-ium iodide (18.9 mg, 0.030 mmol, 1 equv) in dichloromethane (0.3 mL), triethylamine (10.5 uL, 0.075 mmol, 2.5 equiv) was added. After 16 hours of stirring, the reaction mixture was rotovapped and diluted with 1.0 mL of N, N-dimethylformamide and purified by reverse-phase prep HPLC (95-15% H2O/MeOH, 40 mL/min, 45 min). Lyophilization from H2O/MeCN provided the title compound as a yellow powder (7.1 mg, 28% yield TFA salt).

1H NMR (500 MHz, DMSO-d6) δ 7.61 – 7.45 (m, 3H), 7.39 – 7.28 (m, 2H), 7.13 – 7.05 (m, 2H), 7.02 (d, J = 7.0 Hz, 1H), 6.76 – 6.62 (m, 2H), 6.52 (t, J = 5.9 Hz, 1H), 6.12 – 5.83 (m, 2H), 5.11 (dt, J = 13.1, 5.3 Hz, 1H), 4.51 (td, J = 10.4, 9.6, 4.8 Hz, 1H), 4.26 – 3.95 (m, 2H), 3.60 (ddd, J = 27.3, 10.5, 6.3 Hz, 1H), 3.49 – 3.33 (m, 1H), 3.29 – 3.16 (m, 3H), 3.01 (s, 3H), 3.00 –2.89 (m, 2H), 2.75 (ddd, J = 17.1, 4.5, 2.6 Hz, 1H), 2.59 – 2.52 (m, 1H), 2.22 – 1.90 (m, 3H),1.55 (q, J = 7.3 Hz, 2H), 1.45 – 1.14 (m, 12H). LRMS (ESI) calculated for [M+H]^+^ 835.32,found 834.71.

**Synthetic Scheme 3**: ATP-competitive degraders

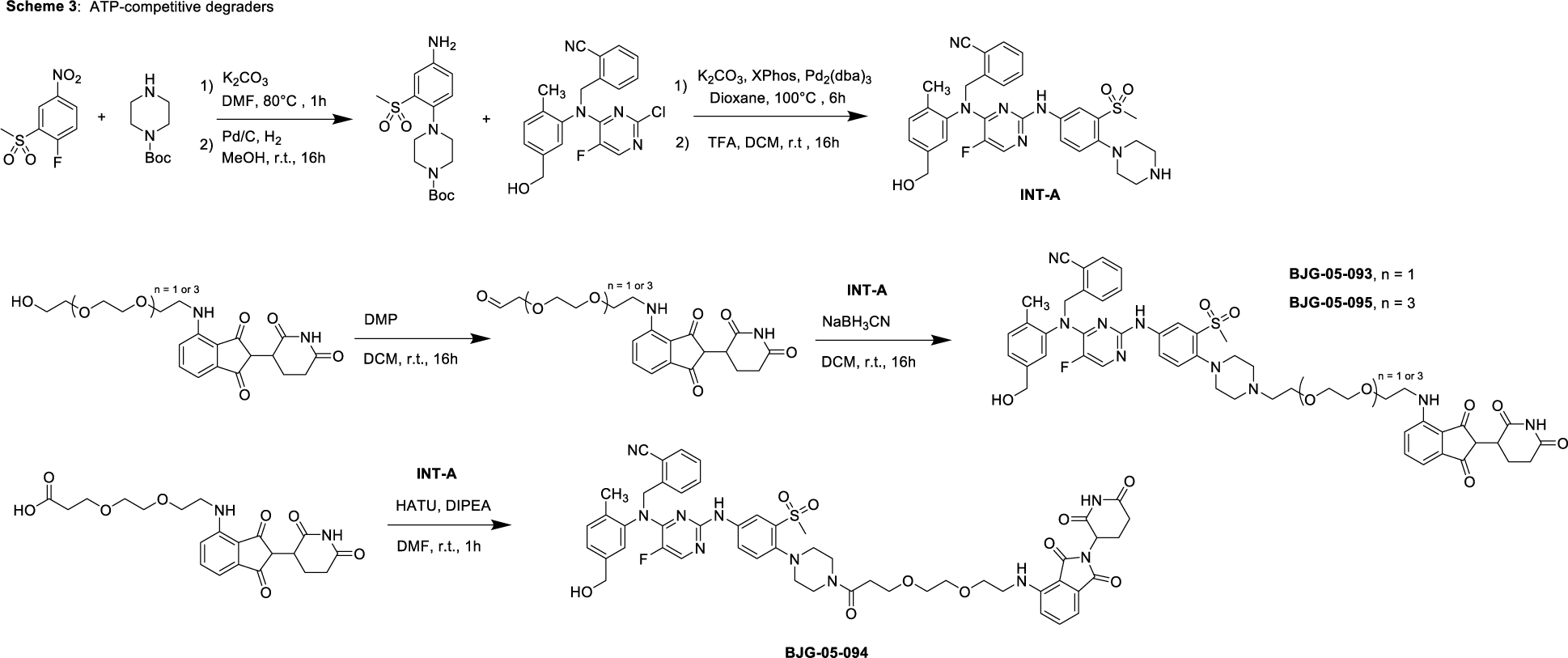

Tert-butyl 4-(4-amino-2-(methylsulfonyl)phenyl)piperazine-1-carboxylate

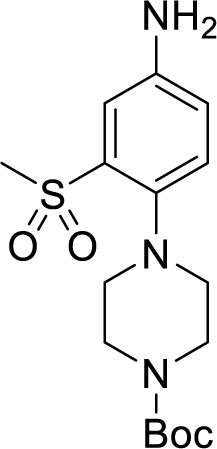

A suspension of 1-fluoro-2-(methylsulfonyl)-4-nitrobenzene (767 mg, 3.5 mmol, 1.0 equiv), tert-butyl piperazine-1-carboxylate (1.956 g, 10.5 mmol, 3.0 equiv), and potassium carbonate (967 mg, 7.0 mmol, 2.0 equiv) in N, N-dimethylformamide (7.0 ml) was stirred at 80°C for 1 hour.

The reaction mixture was diluted with ethyl acetate (100 ml), washed with 0.5M HCl, followed by H2O (3x). The organic fractions were combined, washed with brined, dried over MgSO4, filtered and concentrated in vacuo, yielding 1.355g of yellow solid (∼93% purity by NMR). To a suspension of the crude solids in methanol (25 ml), 10 wt % Pd/C (355 mg, 0.35 mmol, 0.1 equiv) was added. The reaction mixture was sparged with H2 gas for 7 minutes then stirred at room temperature for 16 hours. The mixture was filtered through celite, washing with MeOH (50 ml), and filtrate was concentrated to afford the title compound as a light brown powder (1.248 g, 99% yield, >90 % purity by UPLC-MS). LRMS: [M+H]^+^ found 355.97 2-(((5-fluoro-2-((3-(methylsulfonyl)-4-(piperazin-1-yl)phenyl)amino)pyrimidin-4-yl)(5- (hydroxymethyl)-2-methylphenyl)amino)methyl)benzonitrile

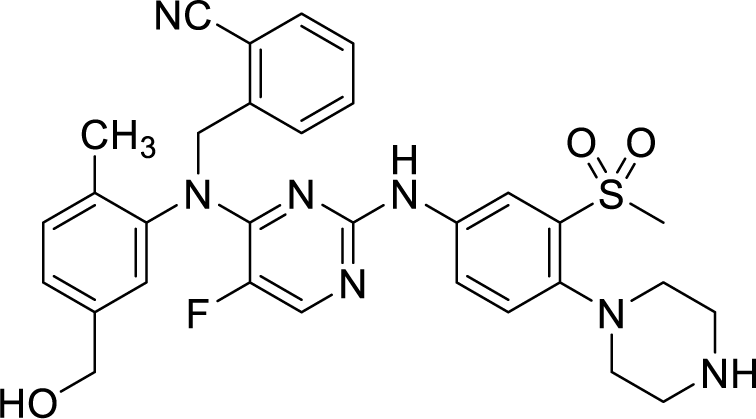

A suspension of 2-(((2-chloro-5-fluoropyrimidin-4-yl)(5-(hydroxymethyl)-2- methylphenyl)amino)methyl)benzonitrile (synthesized according to literature^2^) (765 mg, 2.0 mmol, 1 equiv), tert-butyl 4-(4-amino-2-(methylsulfonyl)phenyl)piperazine-1-carboxylate (781 mg, 2.2 mmol, 1.1 equiv), Tris(dibenzylideneacetone)dipalladium(0) (91.6 mg, 0.10 mmol, 0.05 equiv), XPhos (95.3 mg, 0.20 mmol, 0.10 equiv), and potassium carbonate (829 mg, 6.0 mmol, 3.0 equiv) in dioxane (20 ml) was sparged with N2 for 10 minutes. The reaction mixture was then stirred at 100°C for 6 hours. After cooling to room temperature, the mixture was filtered through celite, washing with ethyl acetate (50 ml), and concentrated in vacuo. The crude material was purified via silica gel chromatography (30% -> 100% ethyl acetate/hexanes) to afford 1.025g of the boc-protected intermediate (beige solids, LRMS: [M+H]^+^ found 701.90). To a suspension of boc-protected intermediate in dichloromethane (5 ml), TFA (1 ml) was added. The reaction mixture was stirred at room temperature for 16 hours. The reaction mixture was concentrated in vacuo to afford the title compound (1.563 g, 73% yield over 2 steps). LRMS: [M+H]^+^ found 601.81) 2-(((2-((4-(4-(2-(2-(2-((2-(2,6-dioxopiperidin-3-yl)-1,3-dioxoisoindolin-4- yl)amino)ethoxy)ethoxy)ethyl)piperazin-1-yl)-3-(methylsulfonyl)phenyl)amino)-5- fluoropyrimidin-4-yl)(5-(hydroxymethyl)-2-methylphenyl)amino)methyl)benzonitrile **BJG-05- 093**

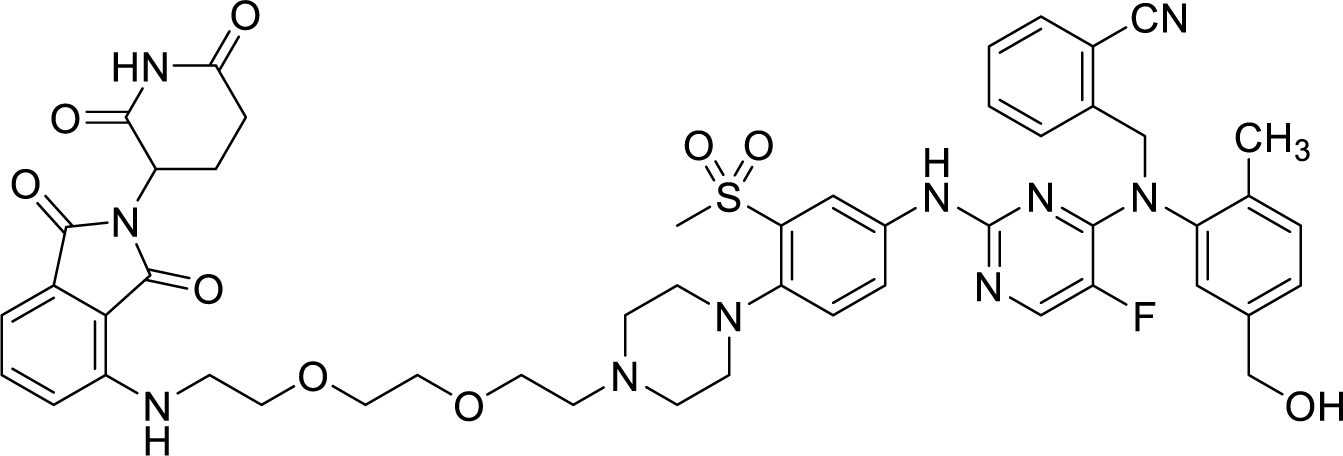

To a solution of solution of 2-(2,6-dioxopiperidin-3-yl)-4-((2-(2-(2- hydroxyethoxy)ethoxy)ethyl)amino)isoindoline-1,3-dione (20.3 mg, 0.050 mmol, 1 equiv) in dichloromethane (1.0 mL), dess-martin periodane (31.8 mg, 0.075 mmol, 1.5 equiv) was added. After 16 hours of stirring at room temperature, the reaction mixture was diluted with 5.0 mL of dichloromethane, filtered and concentrated. To a solution of the crude aldehyde and 2-(((5- fluoro-2-((3-(methylsulfonyl)-4-(piperazin-1-yl)phenyl)amino)pyrimidin-4-yl)(5- (hydroxymethyl)-2-methylphenyl)amino)methyl)benzonitrile (24.9 mg, 0.030 mmol, 1 equiv) in methanol (0.60 ml), sodium cyanoborohydride (5.6 mg, 0.090 mmol, 3 equiv) was added. After 16 hours of stirring at room temperature, the reaction mixture was diluted with 2.0 mL of methanol, filtered and purified by reverse-phase prep HPLC (95-15% H2O/MeOH, 40 mL/min, 45 min). Lyophilization from H2O/MeCN provided the title compound as a yellow powder (8.8 mg, 18% yield TFA salt).

^1^H NMR (500 MHz, DMSO-*d*6) δ 11.09 (s, 1H), 9.65 (s, 1H), 8.29 (s, 1H), 8.02 (d, *J* = 5.6 Hz, 1H), 7.78 (d, *J* = 7.7 Hz, 2H), 7.71 – 7.60 (m, 2H), 7.58 (dd, *J* = 8.6, 7.1 Hz, 1H), 7.45 (td, *J* = 7.5, 1.5 Hz, 1H), 7.31 (d, *J* = 8.8 Hz, 1H), 7.23 – 7.12 (m, 3H), 7.06 – 6.99 (m, 2H), 6.61 (t, *J* = 5.8 Hz, 1H), 5.52 (s, 1H), 5.12 (t, *J* = 5.6 Hz, 1H), 5.05 (dd, *J* = 12.8, 5.4 Hz, 1H), 4.37 (d, *J* = 5.5 Hz, 2H), 3.67 – 3.44 (m, 11H), 3.30 (s, 3H), 2.94 – 2.80 (m, 3H), 2.62 – 2.51 (m, 3H), 2.03 (s, 3H), 2.01 – 1.98 (m, 1H). LRMS (ESI) calculated for [M+H]^+^ 989.37, found 988.73. 2-(((2-((4-(4-(3-(2-(2-((2-(2,6-dioxopiperidin-3-yl)-1,3-dioxoisoindolin-4- yl)amino)ethoxy)ethoxy)propanoyl)piperazin-1-yl)-3-(methylsulfonyl)phenyl)amino)-5- fluoropyrimidin-4-yl)(5-(hydroxymethyl)-2-methylphenyl)amino)methyl)benzonitrile **BJG-05- 094**

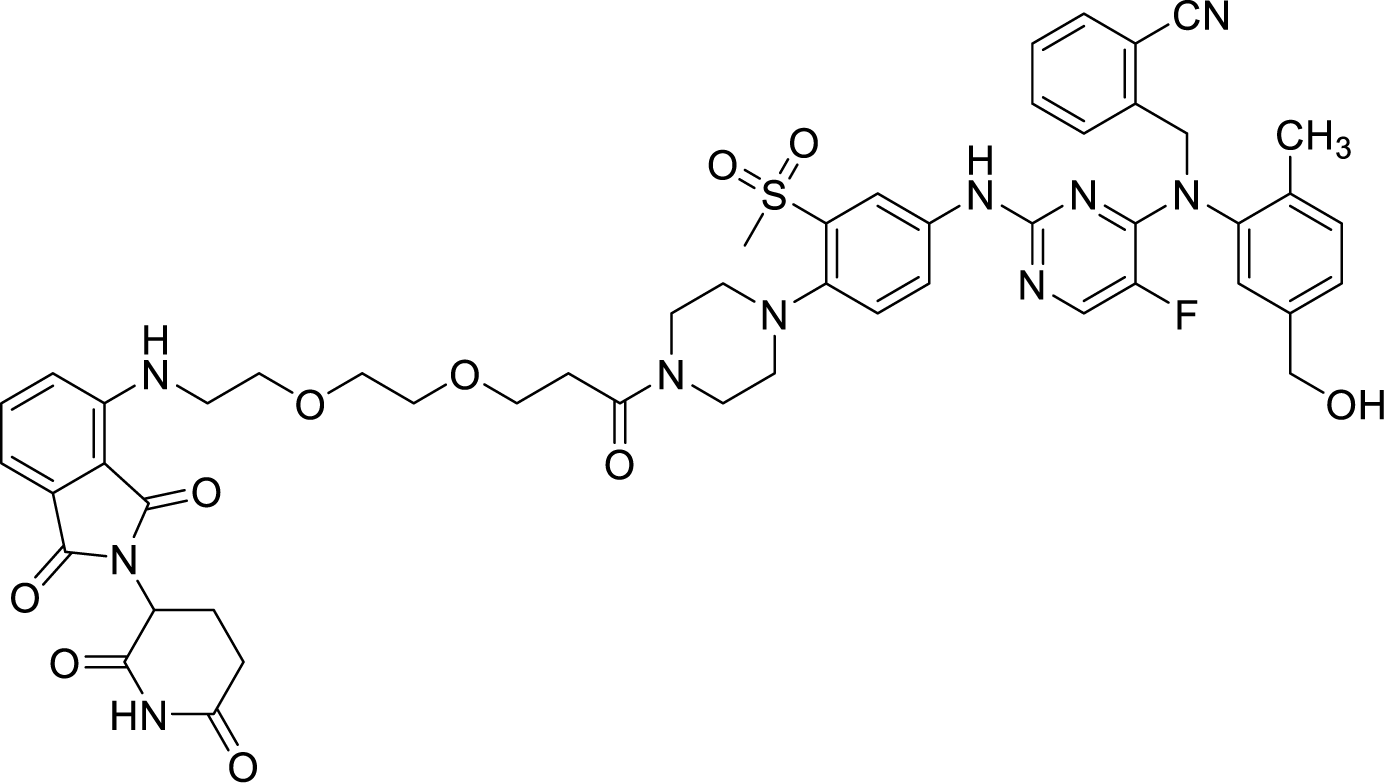

To a solution of solution of 3-(2-(2-((2-(2,6-dioxopiperidin-3-yl)-1,3-dioxoisoindolin-4- yl)amino)ethoxy)ethoxy)propanoic acid (20.7 mg, 0.040 mmol, 1 equiv) in N, N-dimethylformamide (0.5 mL), disopropylamine (34.8 uL, 0.200 mmol, 5 equiv) and HATU (15.2 mg, 0.040 mmol, 1 equiv) were added. After stirring the mixture at room temperature for 5 minutes, 2-(((5-fluoro-2-((3-(methylsulfonyl)-4-(piperazin-1-yl)phenyl)amino)pyrimidin-4-yl)(5- (hydroxymethyl)-2-methylphenyl)amino)methyl)benzonitrile (33.2 mg, 0.040 mmol, 1 equiv) was added. After 2 hours of stirring, the reaction mixture was diluted with 1.0 mL of N, N-dimethylformamide and purified by reverse-phase prep HPLC (95-15% H2O/MeOH, 40 mL/min, 45 min). Lyophilization from H2O/ACN provided the title compound as a yellow powder (11.7 mg, 29% yield TFA salt).

^1^H NMR (500 MHz, DMSO-*d*6) δ 11.09 (s, 1H), 9.66 (s, 1H), 8.31 (d, *J* = 2.6 Hz, 1H), 8.02 (d, *J* = 5.6 Hz, 1H), 7.78 (dd, *J* = 7.8, 1.3 Hz, 2H), 7.70 – 7.61 (m, 2H), 7.57 (dd, *J* = 8.6, 7.0 Hz, 1H), 7.45 (td, *J* = 7.5, 1.4 Hz, 1H), 7.30 (d, *J* = 8.7 Hz, 1H), 7.23 – 7.10 (m, 3H), 7.06 – 6.97 (m, 2H), 6.61 (t, *J* = 5.8 Hz, 1H), 5.52 (s, 1H), 5.11 (t, *J* = 5.7 Hz, 1H), 5.05 (dd, *J* = 12.7, 5.4 Hz, 1H), 4.36 (d, *J* = 5.6 Hz, 2H), 3.69 – 3.60 (m, 4H), 3.60 – 3.50 (m, 5H), 3.46 (q, *J* = 5.6 Hz, 2H), 2.94 – 2.77 (m, 5H), 2.63 – 2.51 (m, 4H), 2.06 – 1.97 (m, 4H). LRMS (ESI) calculated for [M+H]^+^ 1017.36, found 1016.63. 2-(((2-((4-(4-(14-((2-(2,6-dioxopiperidin-3-yl)-1,3-dioxoisoindolin-4-yl)amino)-3,6,9,12- tetraoxatetradecyl)piperazin-1-yl)-3-(methylsulfonyl)phenyl)amino)-5-fluoropyrimidin-4-yl)(5- (hydroxymethyl)-2-methylphenyl)amino)methyl)benzonitrile **BJG-05-095**

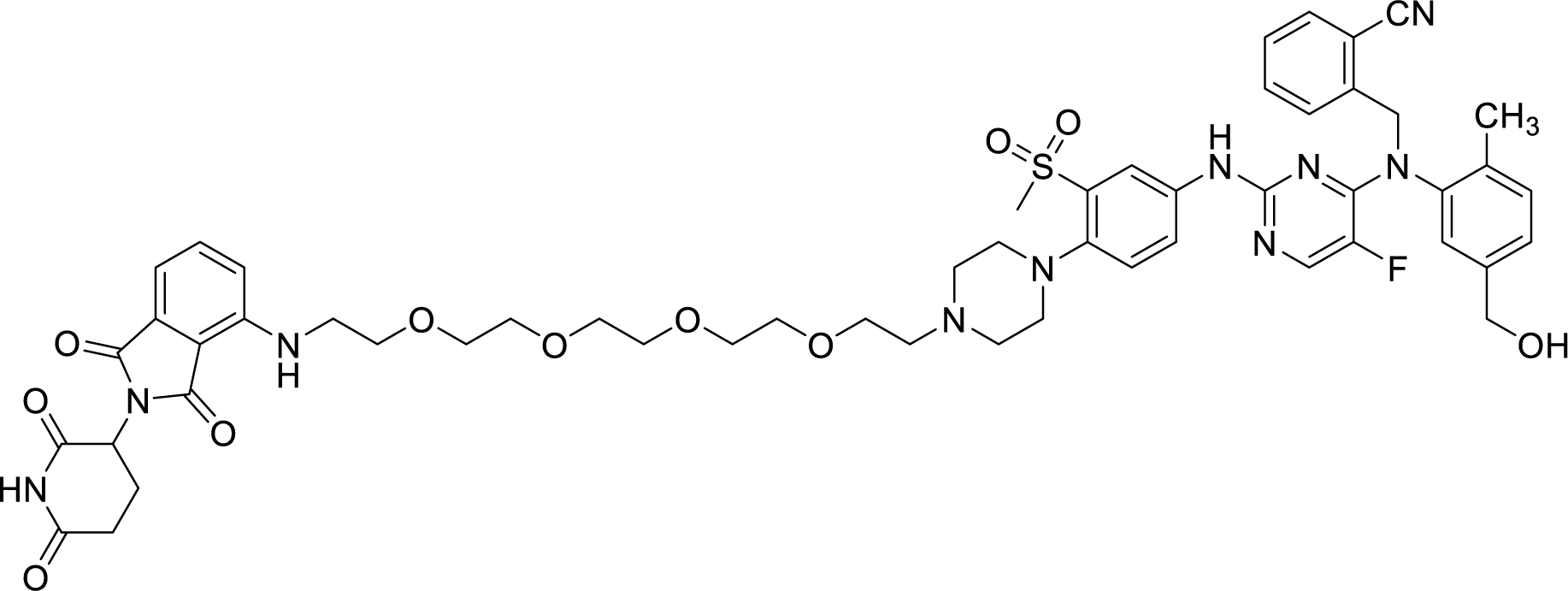

To a solution of solution of 2-(2,6-dioxopiperidin-3-yl)-4-((14-hydroxy-3,6,9,12- tetraoxatetradecyl)amino)isoindoline-1,3-dione (39.5 mg, 0.080 mmol, 1 equiv) in dichloromethane (1.6 mL), dess-martin periodane (50.9 mg, 0.120 mmol, 1.5 equiv) was added. After 16 hours of stirring at room temperature, the reaction mixture was diluted with 5.0 mL of dichloromethane, filtered and concentrated. To a solution of the crude aldehyde and 2-(((5- fluoro-2-((3-(methylsulfonyl)-4-(piperazin-1-yl)phenyl)amino)pyrimidin-4-yl)(5- (hydroxymethyl)-2-methylphenyl)amino)methyl)benzonitrile (66.4 mg, 0.080 mmol, 1 equiv) in methanol (1.0 ml), sodium cyanoborohydride (15.1 mg, 0.240 mmol, 3 equiv) was added. After 16 hours of stirring at room temperature, the reaction mixture was diluted with 2.0 mL of methanol, filtered and purified by reverse-phase prep HPLC (95-15% H2O/MeOH, 40 mL/min, 45 min). Lyophilization from H2O/MeCN provided the title compound as a yellow powder (9.6 mg, 11% yield TFA salt).

^1^H NMR (500 MHz, DMSO-*d*6) δ 11.09 (s, 1H), 9.63 (s, 1H), 8.27 (d, J = 2.6 Hz, 1H), 8.01 (d, J = 5.6 Hz, 1H), 7.95 (dd, J = 7.9, 1.2 Hz, 1H), 7.81 – 7.71 (m, 2H), 7.70 – 7.60 (m, 3H), 7.57 (dd, J = 8.6, 7.0 Hz, 1H), 7.45 (td, J = 7.5, 1.3 Hz, 2H), 7.33 (d, J = 8.8 Hz, 1H), 7.23 – 7.15 (m, 3H), 7.13 (d, J = 8.6 Hz, 1H), 7.06 – 6.95 (m, 2H), 6.60 (t, J = 5.8 Hz, 1H), 5.51 (s, 1H), 5.27 – 4.92 (m, 3H), 4.37 (s, 2H), 3.61 (t, J = 5.4 Hz, 2H), 3.59 – 3.42 (m, 20H), 2.95 – 2.81 (m, 5H), 2.62 – 2.55 (m, 1H), 2.07-1.97 (br s, 4H). LRMS (ESI) calculated for [M+H]^+^ 1077.42, found 1076.65.

